# Nuclear envelope rupture in cardiomyocytes orchestrates early transcriptomic changes and immune activation in *LMNA*-related dilated cardiomyopathy that are reversed by LINC complex disruption

**DOI:** 10.1101/2024.06.11.598511

**Authors:** Noam Zuela-Sopilniak, Julien L.P. Morival, Ioannis Ntekas, Margaret A. Elpers, Rohit Agarwal, Sarah J. Henretta, Jacob D. Odell, Iwijn De Vlaminck, Jan Lammerding

**Affiliations:** Weill Institute for Cell and Molecular Biology, Cornell University, Ithaca, NY, USA; Meinig School of Biomedical Engineering, Cornell University, Ithaca, NY, USA

**Keywords:** Laminopathy, nuclear damage, transcriptomics, cell-cell-communication

## Abstract

Mutations in the *LMNA* gene, which encodes the nuclear envelope (NE) proteins lamins A and C, cause dilated cardiomyopathy (*LMNA*-DCM) and other diseases. The pathogenic mechanisms for *LMNA*-DCM remain poorly understood, limiting current treatment options and leading to high mortality amongst patients. We developed a mouse model with inducible, cardiomyocyte-specific *Lmna* deletion (cKO) and performed comprehensive bulk, single-nucleus, and spatial transcriptomic analyses across disease progression. Our analysis identified key disease-driving genes involved in cellular responses to DNA damage, cytosolic pattern recognition receptor signaling, and innate immunity that originated from two disease-specific cardiomyocyte subpopulations. Spatial mapping revealed aberrant interactions between these cardiomyocytes, fibroblasts, and immune cells, contributing to tissue-wide transcriptional changes in cKO hearts. Concurrent cardiomyocyte-specific disruption of the LINC complex, which transmits cytoskeletal forces to the nucleus, substantially reduced NE rupture in cKO cardiomyocytes, normalized expression of more than half of the dysregulated genes, and dramatically improved cardiac function and survival in cKO mice. These findings suggest that NE rupture in cKO cardiomyocytes triggers cytosolic DNA sensing pathways and maladaptive cell-cell communication with fibroblasts and immune cells, leading to fibrosis and inflammation driving *LMNA*-DCM pathogenesis.

## Introduction

Lamins A and C (lamin A/C) are nuclear intermediate filaments that provide structural support to the nucleus and regulate numerous cellular processes, such as chromatin organization, nuclear pore positioning, signal transduction, transcriptional regulation, and DNA damage repair^1–3^. Inherited or de novo mutations in the *LMNA* gene cause a group of disorders known as laminopathies, and although lamins A and C are expressed in nearly all tissues, most *LMNA* mutations primarily affect striated muscle and tendons. *LMNA*-related dilated cardiomyopathy (*LMNA*-DCM) is the most prevalent laminopathy and has a worse clinical prognosis compared to other congenital forms of DCM^4–7^, frequently resulting in sudden cardiac death, even in the absence of systolic dysfunction^5^. Currently, no effective therapies exist for *LMNA*-DCM. Furthermore, the molecular pathogenic mechanism for *LMNA*-DCM remains largely unresolved, presenting a major hurdle in developing effective treatment strategies.

A major question in the study of striated muscle laminopathies is whether the tissue-specific phenotypes arise from disrupted lamina-chromatin interactions and altered transcription factor binding (“gene regulation hypothesis”) or from impaired structural function of the nuclear lamina that results in NE rupture in mechanically stressed tissues (“structural hypothesis”)^3,8^. In support of the structural hypothesis, *LMNA* mutations linked to striated muscle disease often impair lamin filament assembly, resulting in mechanically fragile nuclei and increased incidence of NE rupture^9–11^. NE rupture, although typically repaired rapidly^12,13^, can lead to DNA damage and activate cytosolic DNA sensing pathways, such as cGAS/STING^9,14^, which might trigger additional pathogenic signaling cascades that lead to cardiac dysfunction. However, it is unclear if NE ruptures are indeed the drivers for cardiac functional decline.

Previous studies using mouse models of laminopathies and late-stage patient tissue have identified several signaling pathways known to be misregulated in *LMNA*-DCM, including AMP-activated protein kinase (AMPK), myocardin-related transcription factor (MRTF), extracellular signal-regulated kinase 1/2 (ERK1/2), protein kinase C (AKT)/mammalian target of rapamycin (mTOR), Wnt/β-Catenin, transforming growth factor-β (TGF-β), and various DNA damage response pathways^15,16^. Yet, despite various attempts to target these altered signaling pathways in preclinical models, none of the approaches have been curative, and most have shown only modest effects on survival^16^. These limited curative outcomes raise the question whether the identified dysregulated pathways are indeed disease-causing or arise as downstream consequences of the failing heart or other disease-associated processes.

Here, we addressed these questions by performing a multi-level transcriptomic analysis in left ventricle tissue obtained from an inducible *LMNA*-DCM mouse model (cKO). By examining transcriptional changes occurring early in disease, which become more pronounced with disease progression, we identified the activation of cGAS-independent cytosolic PRR pathways, which likely respond to exposure of genomic DNA to the cytoplasm upon NE rupture^17,18^, triggering innate immune activation and cardiac inflammation. Single-nucleus RNA sequencing (snRNA-seq) identified two disease-specific cardiomyocyte subpopulations that were responsible for these tissue-level transcriptomic changes in cKO hearts. Single-nucleus and spatial transcriptomic analyses indicated that these cardiomyocytes activate fibroblasts and innate immune cells. Supporting a crucial role of NE rupture in driving pathogenic transcriptomic changes in cKO hearts, concurrent cardiomyocyte-specific disruption of the LINC complex^9,10^, which reduces the physical stress on diseased cardiomyocytes^10,19–22^, significantly reduced NE ruptures and rescued the expression of approximately half of the dysregulated genes in cKO hearts, including many of those involved in PRR and inflammation. Intriguingly, LINC complex disruption resulted in near complete rescue of cardiac function and extended survival of cKO mice from weeks to over one year. Together, our findings offer new insight into the molecular mechanisms driving *LMNA*-DCM and offer new targets to focus on for future therapeutic studies.

## Results

### Cardiomyocyte-specific depletion of lamin A/C in adult mice causes severe dilated cardiomyopathy

We generated cKO mice using a tamoxifen-responsive Cre driven by the α-myosin heavy chain promoter and a floxed *Lmna* gene. The cKO mice offer several key advantages for identifying disease-driving mechanisms. Unlike complete lamin A/C deletion, this model leads to a reduction in lamin A/C protein levels mirroring that observed in many clinical cases of *LMNA*-DCM^44^. In addition, inducing lamin A/C depletion in adulthood allowed us to disentangle the effects of lamin A/C on cardiac development from those on cardiac function. The cKO mice also reduce confounding effects present in many other *LMNA*-DCM animal models, such as variability in disease onset and severity, involvement of multiple tissues, and differences in animal size and motility across genotypes^45–47^. Following tamoxifen treatment of adult cKO mice (Figure 1A), we observed a ≈50-70% reduction in lamin A levels in cardiomyocytes by 11 days post-injection (dpi), compared to tamoxifen-treated littermate controls that expressed the MerCreMer recombinase but lacked the floxed *Lmna* alleles (cWT) (Figure 1B-C) and cKO mice treated with vehicle-only (Suppl. Figure S1A). Lamin A levels in non-cardiomyocytes were not affected (Figure 1B; Suppl. Figure S1B), as expected. Immunofluorescence staining for lamin A/C showed a similar trend to lamin A, albeit with a smaller effect size, likely due to weaker antibody labeling (Suppl. Figure S1B-C). Since the anti-lamin A antibody produced better labeling in cardiac tissue sections, and results obtained with this antibody matched those of a recent study in the same genetic model^48^, subsequent analyses are based on immunofluorescence labeling for lamin A. Remarkably, even the partial depletion of lamin A/C in tamoxifen-treated cKO mice was sufficient to trigger rapid development of severe DCM (Figure 1D). Histological analysis demonstrated increased cardiac fibrosis in cKO hearts at 11-dpi, followed by substantial left ventricular enlargement of cKO hearts by 25-dpi (Figure 1D-E; Suppl. Figure S1D). Echocardiographic analysis revealed a slight reduction in cardiac function in tamoxifen-treated cKO mice at 11-dpi that progressively worsened (Figure 1F; Suppl. Figure S1E-H). In contrast, tamoxifen-treated cWT controls and vehicle-only treated cKO mice maintained normal cardiac morphology and function (Figure 1D-F; Suppl. Figure S1D-H).

**Figure 1:**
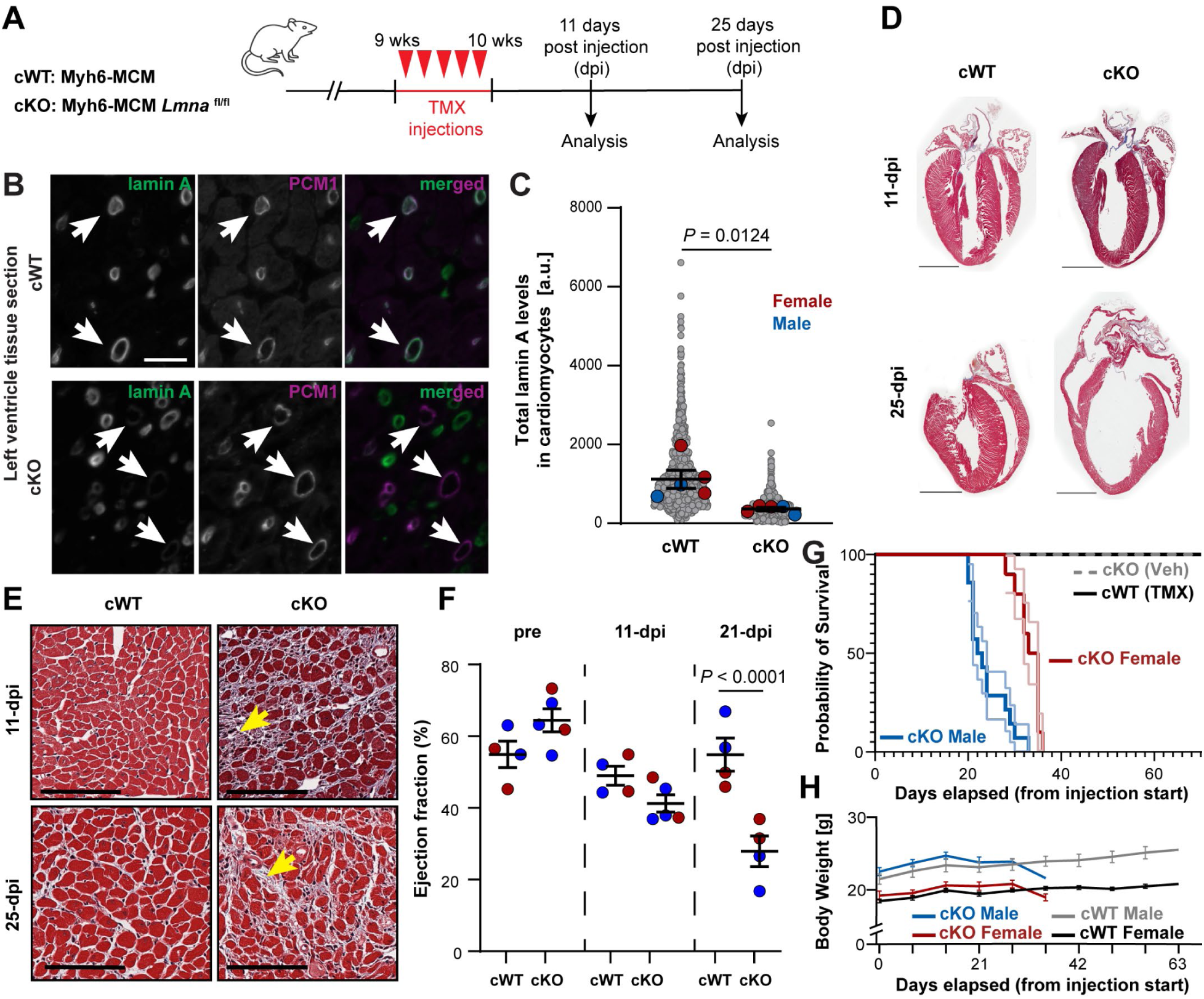
Cardiac-specific lamin A depletion in adult mice causes rapid onset dilated cardiomyopathy. **(A)** Experimental timeline for tamoxifen-induced *Lmna* deletion with echocardiography and tissue collection at indicated time points. **(B)** Representative immunofluorescence images of cardiac tissues from cWT (top) and cKO (bottom) mice labeled for lamin A (green) and PCM1 (magenta), a cardiomyocyte-specific nuclear marker. Scale bar 20 μm. Arrows indicate representative cardiomyocytes. **(C)** Quantification of total lamin A levels in cardiomyocytes. Larger colored circles represent average values for individual male (blue) or female (red) mice. Smaller gray circles represent individual cardiomyocytes. *N* = 5 animals per genotype. Statistics are based on the mean values of individual animals. **(D-E)** Masson’s trichrome staining of heart sections **(D)** showing progressive thinning of the left ventricle free wall and enlargement of cKO hearts. Scale bar: 2 mm, and **(E)** increased collagen deposition (yellow arrows) in cKO hearts at both 11- and 25-dpi. Scale bar: 100 μm. **(F)** Left ventricular ejection fraction (EF) measured by echocardiography. Symbols represent %EF of individual (red for female and blue for male) mice. Error bars represent mean ± s.e.m. *N* = 4-5 animals per genotype. **(G)** Kaplan-Meier survival plot for tamoxifen-treated cWT mice (*N* = 14 males and 11 females), vehicle-only treated cKO mice (*N* = 10 males and 2 females) and tamoxifen-treated cKO mice (*N* = 14 males and 10 females). **(H)** Body weight measurements performed during the survival study on the mice in panel (G) before (day 0) and after tamoxifen injections.

Consistent with the severely impaired cardiac function, tamoxifen-treated cKO mice had substantially reduced survival compared to cWT and vehicle-only controls (Figure 1G). Male cKO mice died by 22-dpi, whereas female cKO mice lived until 34-dpi, despite similar depletion of lamin A as the male mice (Figure 1C). This sex-specific difference in survival is consistent with previous reports in other *LMNA*-DCM mouse models^49^. Tamoxifen-treated cWT controls did not exhibit premature death (Figure 1G), indicating that the Cre recombinase had no significant cardiotoxic effects at the administered tamoxifen dose. Intriguingly, whereas other striated laminopathy mouse models commonly show loss of body weight and reduced motility^50^, cKO mice died suddenly, without preceding weight loss (Figure 1H) or other external indications. This sudden cardiac death mirrors symptoms in *LMNA*-DCM patients^51^. Collectively, these findings indicate that even partial reduction of lamin A/C protein levels in cardiomyocytes results in severe and rapid-onset DCM^6,44,52,53^.

### Transcriptomic profiling reveals early and progressive gene dysregulation in cKO hearts

To comprehensively characterize the transcriptional changes resulting from cardiac-specific *Lmna* deletion, we performed bulk RNA-seq, snRNA-seq, and spatially resolved RNA sequencing on left ventricular tissue from tamoxifen-treated cKO mice and cWT controls at 11- and 25-dpi (Figure 2A). These timepoints correspond to early disease onset and severe cardiac dysfunction, respectively (Figure 1D-F). We initially focused on bulk RNA-seq data to capture broad transcriptional shifts in *LMNA*-DCM cardiac tissue and to identify key pathways for deeper investigation at the cellular and spatial level. Transcriptional differences between cKO and cWT hearts were already detectable at 11-dpi and became more pronounced by 25-dpi (Figure 2B). The number of differentially expressed genes (DEGs) increased from 86 at 11-dpi to 2,951 at 25-dpi (Suppl. Figure S2A). The majority of DEGs showed increased expression in the cKO samples (Suppl. Figure S2B), suggesting that cKO hearts have a progressive activation of transcriptional programs. Despite the sex-specific differences in survival (Figure 1G), we did not detect any statistically significant transcriptional differences between male and female cKO mice (Suppl. Figure S3A). Consequently, we combined data from male and female mice in subsequent analyses to increase their statistical power.

**Figure 2:**
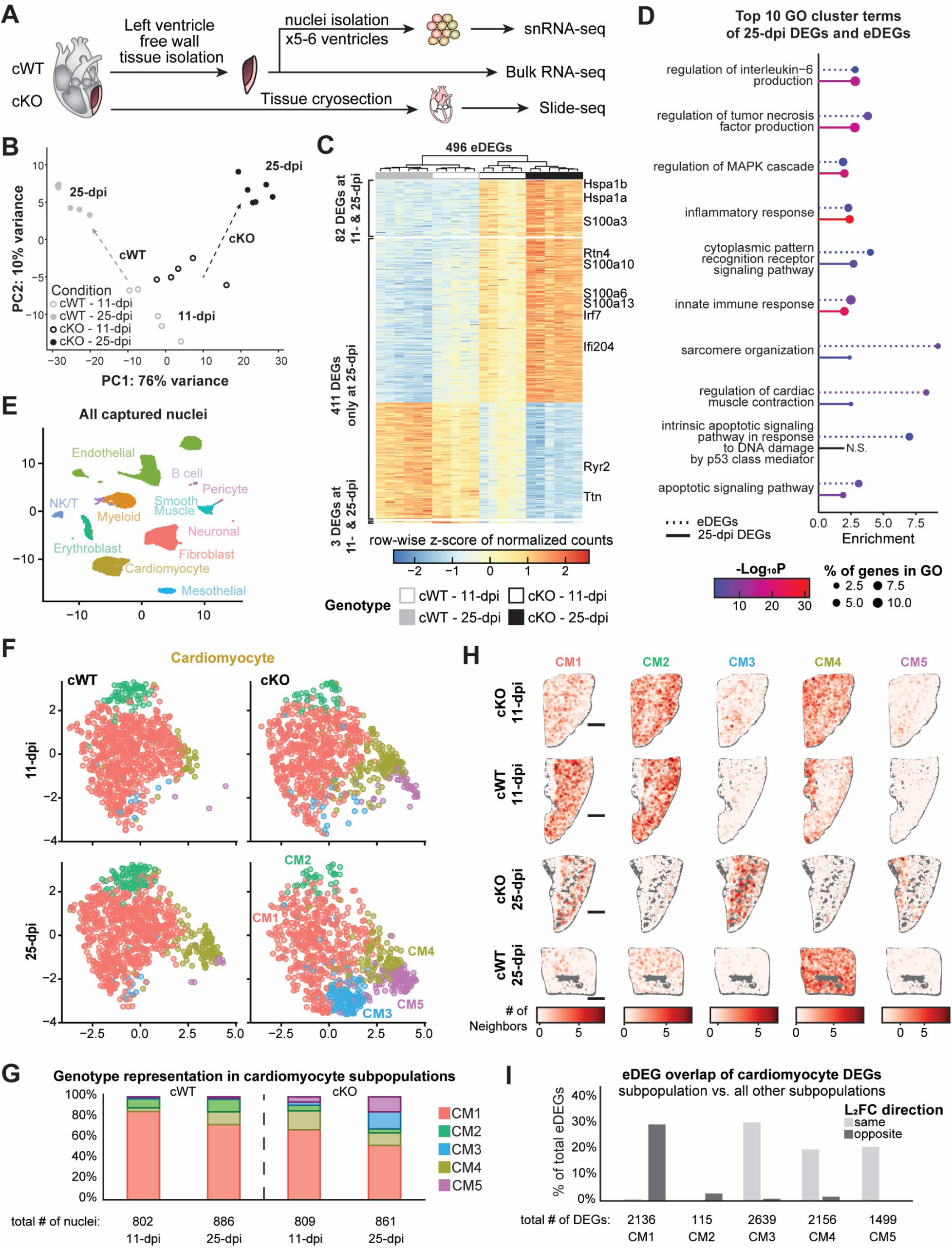
*LMNA*-DCM-associated eDEGs are mainly enriched in two disease-specific cardiomyocyte subpopulations. **(A)** Experimental design for transcriptomic analyses of cKO and cWT hearts, including bulk RNA-seq (*N* = 5-6 animals per genotype), snRNA-seq (*N* = 5-6 animals per genotype), and spatial transcriptomic (*N* = 1 animals per genotype). **(B)** Principal components analysis (PCA) of bulk RNA-seq samples showing progressive genotype-dependent separation of samples from 11- to 25-dpi (arrows). **(C)** Heatmap of row-wise z-score normalized read counts for eDEGs. Genes meeting DEG criteria at both timepoints (85 genes, top and bottom sections) or emerging by 25-dpi (411 genes, middle section) are indicated. Samples are hierarchically clustered. Select genes identified throughout the study are highlighted. **(D)** Gene Ontology (GO) cluster terms enrichment of eDEGs (dashed) and 25-dpi DEGs (solid). N.S. = Not-significant. **(E)** UMAP of integrated snRNA-seq nuclei from cWT (left) and cKO (right) hearts, colored by cell type and shaded by timepoint (11-dpi, light; 25-dpi, dark). **(F)** UMAP of cardiomyocyte nuclei, colored by subpopulation (CM1-CM5), across genotype (left and right) and timepoint (top and bottom). **(G)** Relative composition of cardiomyocyte subpopulations shown as the percentage of total cardiomyocyte nuclei sequenced per condition. Total number of sequenced nuclei are indicated for each subgroup. **(H)** Spatial neighborhood enrichment maps of cardiomyocyte subpopulations in left ventricular sections from cKO and cWT heart at 11- and 25-dpi. Red intensity indicates local enrichment. Scale bars: 500 μm. **(I)** Percentage of eDEGs overlapping with subpopulation-specific DEGs. Bar color indicates concordant (both up- or down-regulated; light gray) or discordant (one up- and one down-regulated; dark gray) fold change direction of corresponding eDEG and DEG. The total number of DEGs is indicated below each condition.

Given the only subtle transcriptional differences present at 11-dpi (Figure 2B), we hypothesized that some early changes in gene expression that are statistically significant may fall below our stringent threshold criteria for DEGs, (i.e., |log_2_ fold change | ≥ 1), yet still contribute to disease pathogenesis. To identify which dysregulated genes are likely to be involved in disease progression, we developed an analysis pipeline that uncovered 496 genes that were already statistically significant at 11-dpi and whose change in expression increased further over time, reaching the |log_2_-fold| threshold by 25-dpi (Suppl. Figure S2A). We termed these genes ‘emerging DEGs’ (eDEGs). Hierarchical clustering of samples based on eDEG expression created distinct separation based on genotype and timepoint (Figure 2C), demonstrating that our methodology enables the enhanced detection of genes already dysregulated early in disease and thus likely involved in disease driving processes. In contrast, we identified only 12 genes that were statistically significant at 11-dpi but reversed the direction of change (e.g., from up- to down-regulated) at 25-dpi (Suppl. Figure S2C-D).

To uncover disease-driving biological processes mediated by the eDEGs, we performed comparative Gene Ontology (GO) analysis comparing eDEGs with DEGs identified at 25-dpi (Figure 2D)^36,54^. Among the top 10 enriched GO cluster terms in both gene sets were several processes previously linked to lamin A/C deficiency, such as increased MAPK activation^55^ and apoptosis^56^. Additionally, eDEGs were enriched for terms associated with inflammation, innate immunity, and cardiac function.

### Two cardiomyocyte subpopulations predominantly account for eDEG expression in cKO hearts

To distinguish the individual transcriptional contribution of different cell types to the overall response, we next focused on the snRNA-seq dataset. Cell type annotation identified many of the cell types expected in left ventricular tissue^38^ (Figure 2E). Cardiomyocyte nuclei clustered into five individual subpopulations, two of which, CM3 and CM5, were represented only in cKO hearts and progressively expanded over time (Figure 2F-G), mirroring the increased transcriptional divergence between cKO and cWT control hearts (Figure 2B). Spatial transcriptomics analysis enabled us to map the identified cardiomyocyte subpopulations based on their snRNA-seq transcriptional profiles and to validate that the CM3 and CM5 subpopulations are present in the cKO hearts at both 11-and 25-dpi, but not in the cWT hearts (Figure 2H, Suppl. Figure S4).

We next compared the transcriptomic profile of individual cell subpopulations identified in the snRNA-seq analysis to the eDEGs from the bulk RNA-seq analysis. Remarkably, the cardiomyocyte subpopulations CM3 and CM5 together accounted for over 40% of all eDEGs (Figure 2I, Suppl. Figure S5A-B), despite comprising only 11% of total cardiomyocyte nuclei (Figure 2F). The eDEGs detected in the CM3 and CM5 subpopulations enriched for GO terms involving immunity, apoptosis, and cardiac function (Suppl. Figure S5C). These findings suggest that the identified disease-specific cardiomyocyte CM3 and CM5 subpopulations are involved in promoting *LMNA*-DCM pathogenesis.

### cKO cardiomyocytes exhibit transcriptomic evidence of cytosolic DNA sensing pathway activation

Cells from human patients and mouse models of striated laminopathies frequently show signs of nuclear damage, including chromatin protrusions and NE ruptures^9,45,48,57–59^. The NE ruptures can be visualized by transient accumulation of barrier-to-autointegration factor (BAF) at the NE^60^, where BAF contributes to NE repair^12,13^. Cardiomyocytes of cKO mice, but not cWT controls, frequently exhibited NE ruptures (Figure 3A, Suppl. Figure S6). Consistent with the expected leakage of nuclear DNA into the cytoplasm upon NE rupture^61,62^, eDEGs were significantly enriched for the GO term “cytoplasmic pattern recognition receptor signaling pathway”. This GO term encompasses molecules involved in cytosolic DNA sensing pathways, including cyclic GMP-AMP synthase (cGAS), a major cytoplasmic DNA sensors, and its target STING, which acts as a central node for downstream activation of NF-κB and interferon-regulatory factors (IRFs) in the inflammatory response pathway^17,63^. Ingenuity Pathway Analysis (IPA) revealed that many of the top 25 predicted upstream regulators of the eDEGs are components of the cytosolic DNA-sensing pathway, including STING1, IRF3, IRF7, MyD88, and NF-κB (NFKB) (Table 1). Although GO term analysis of CM3 and CM5 associated eDEGs did not find the same cytosolic DNA sensing pathway enrichment (Suppl. Figure S5C), IPA did identify several upstream regulators involved in the cytosolic DNA-sensing signaling cascade, including Toll-like receptor-9 (TLR9), p53 (TP53), Interleukin-6 (IL6), NF-κB (NFkB), Interleukin-1 (IL1A/B), MyD88, and interferon-γ (IFNG) (Figure 3B). Furthermore, both the single-nucleus and spatial data revealed an enrichment of the GO terms “inflammatory response” and “regulation of defense” (Figure 3 C-E, Suppl. Figure S7A-B), which include cytosolic DNA-sensing molecules, within the CM3 and CM5 subpopulations.

**Figure 3:**
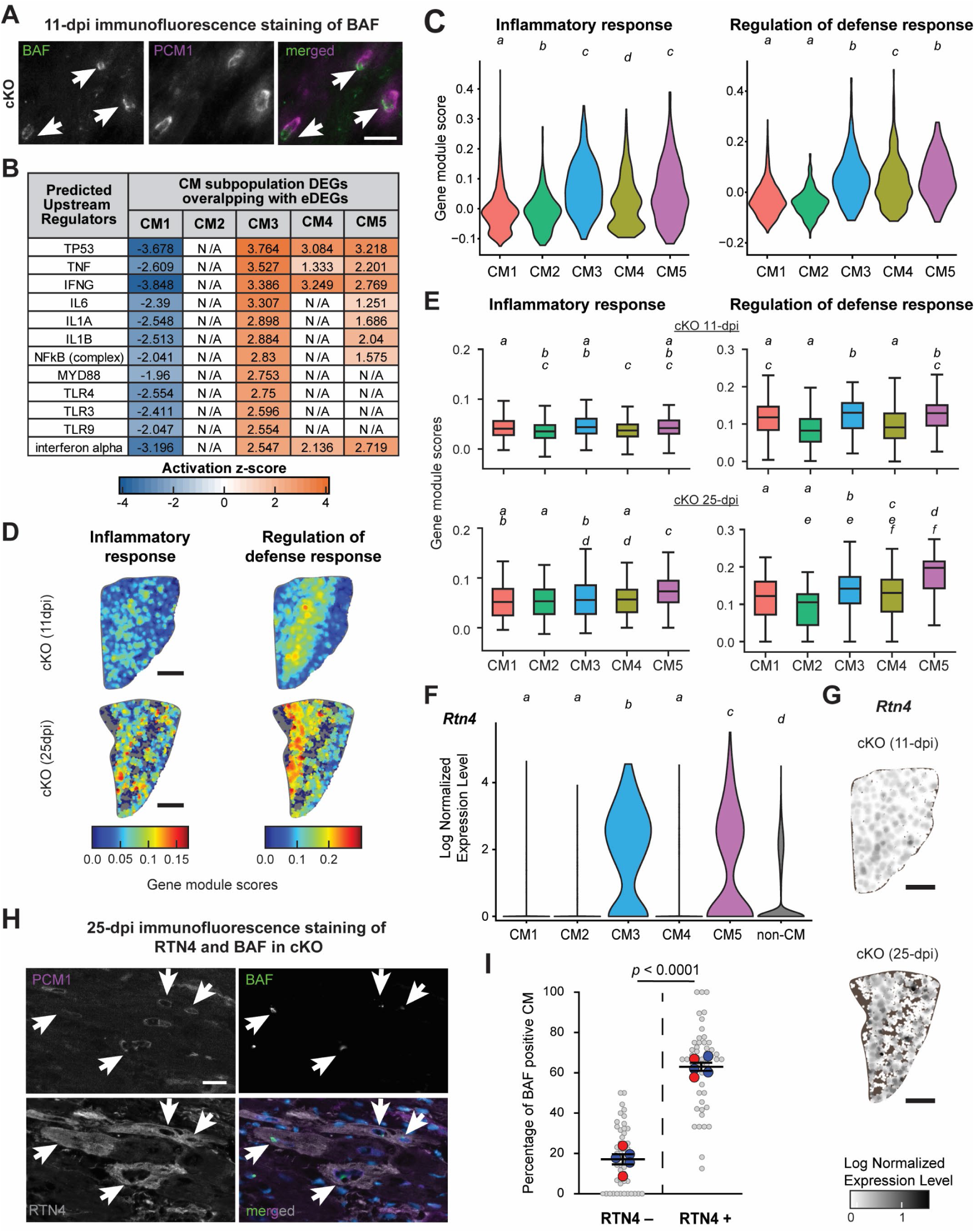
cKO hearts exhibit transcriptomic evidence of cytosolic DNA-sensing pathway activation. **(A)** Immunofluorescence images of cardiac tissues from cKO mice at 11-dpi labeled for BAF (green) to indicate NE rupture, and PCM1 (magenta) to identify cardiomyocytes. Scale bar: 20 μm. Arrows indicate representative cardiomyocytes. **(B)** Predicted activation of the top 40 upstream regulators involved in DNA damage and cytosolic DNA sensing pathways identified using IPA based on cardiomyocyte subpopulation (CM1-CM5)-specific DEGs that overlap eDEGs. N/A indicates the upstream regulator was not predicted. **(C)** Gene module scores for GO terms “inflammatory response” (left) and “regulation of defense” (right) for individual cardiomyocyte subpopulations in the integrated snRNA-seq dataset. **(D)** Spatial distribution and **(E)** quantification of GO term “Inflammatory response” (left) and “Regulation of defense” (right) gene module scores across the **(D)** left ventricle and (**E**) individual cardiomyocyte subpopulations at 11- and 25-dpi in cKO hearts. **(F)** Normalized *Rtn4* expression across CM1-CM5 and all non-cardiomyocyte cells in the integrated snRNA-seq dataset. **(G)** Spatial expression of *Rtn4* in cKO left ventricles at 11-dpi (left) and 25-dpi (right). **(H)** Immunofluorescence labeling for RTN4 (gray), BAF (green), PCM1 (magenta) and nuclei using DAPI (blue) for cKO left ventricles at 25-dpi. Scale bar: 20 μm **(I)** Quantification of BAF-positive nuclei in RTN4-negative (left) and RTN4 positive (right) cardiomyocytes in cKO left ventricles at 25-dpi. Larger colored circles represent average values for individual male (blue) or female (red) mice. *N* = 5 animals. Statistical analysis performed by *t*-test **(C, E, F)** Statistical annotation: Groups that share a letter are not significantly different; different letters indicate *P* < 0.05. Tests: pair-wise Wilcoxon rank sum with Benjamini-Hochberg correction **(C)** and Kruskal–Wallis tests followed by pairwise Mann–Whitney U tests with Benjamini-Hochberg *p*-value adjustment **(E,F)**.

**Table 1:**
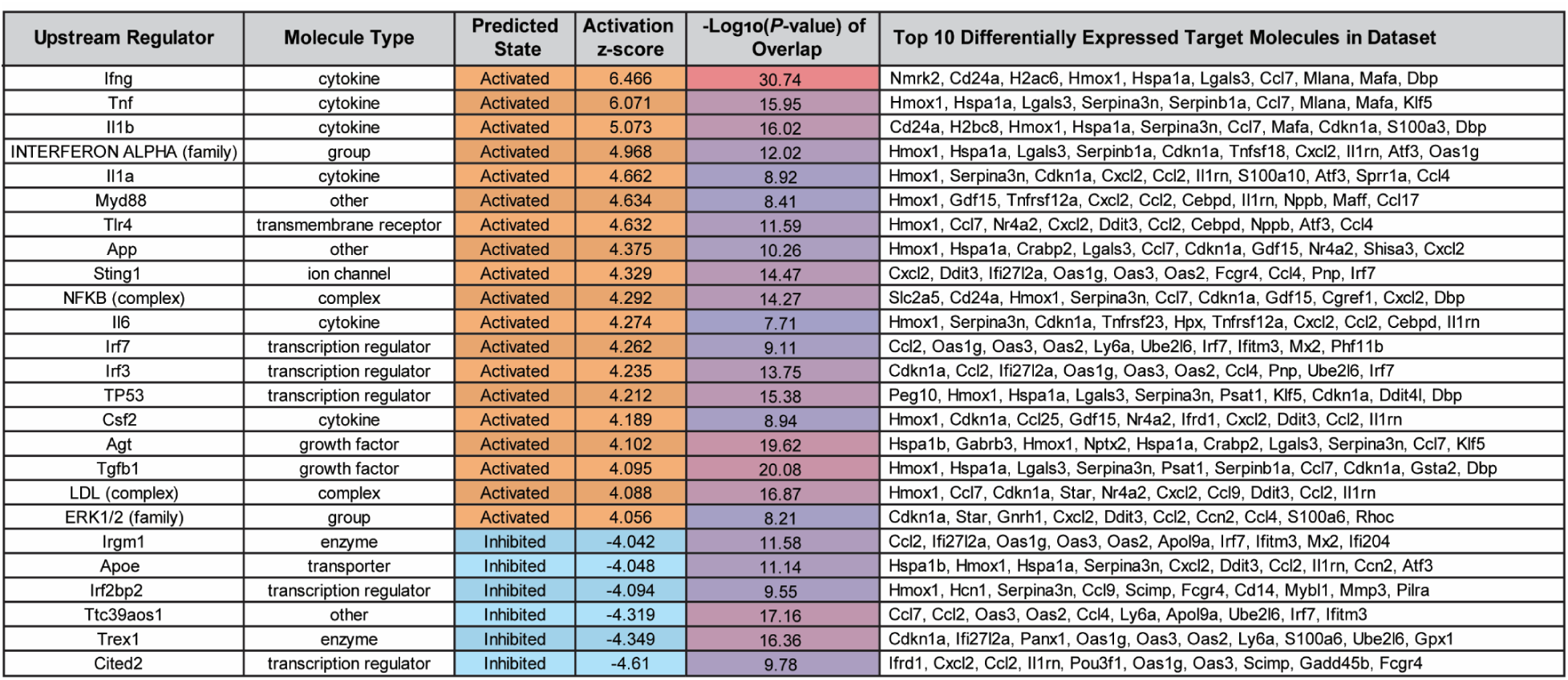
Cytosolic DNA sensing signaling proteins were predicted as upstream regulators of eDEGs. Top 25 upstream regulators, identified by Ingenuity Pathway Analysis (IPA), predicted to be activated or inhibited based on 11-dpi eDEG expression differences between cKO and cWT samples. Top 10 most differentially expressed eDEGs targeted by predicted upstream regulators are displayed.

Despite the significant activation of cytosolic DNA-sensing pathways, only a very small fraction of cardiomyocytes expressed *cGAS* or *Sting* (0.35% and 0.2%, respectively), and neither was significantly expressed in the CM3 and CM5 subpopulations in the snRNA-seq dataset (Suppl. Figure S8A-D). Although snRNA-seq has limited sensitivity to lowly expressed transcripts and therefore cannot definitively exclude *cGas* or *Sting* expression, the rare detection frequency observed is consistent with recent reports that cGAS is essentially absent in cardiomyocytes, and that deletion of either *cGas* or *Sting* does not rescue cardiomyopathy in *LMNA*-DCM mouse models^38,48,64^. The lack of *cGAS*/*STING* activation prompted us to look for alternative cytosolic DNA sensing processes in cardiomyocytes that could contribute to the observed inflammatory response. Among the candidates, *Rtn4* stood out as a highly upregulated eDEG expressed in the CM3 and CM5 subpopulations based on both snRNA-seq (Figure 3F) and spatial transcriptomic analyses (Figure 3G, Suppl. Figure S7C). RTN4 is a resident endoplasmic reticulum (ER) transmembrane protein involved in the trafficking of toll-like receptor (TLRs), particularly TLR9, from the ER to endolysosomes, where cytosolic nucleic acid-sensing TLRs can initiate a signaling cascade for downstream innate immune response, independent of cGAS/STING^17,18,65^. We confirmed significantly higher levels of RTN4 in cKO hearts compared to cWT controls at 25-dpi by immunofluorescence (Suppl. Figure S9) and western blot analysis (Suppl. Figure S10A). We also noticed the upregulation of several genes downstream of TLR activation in the CM3 and CM5 subpopulations, including *Traf3, Tkb1, Irak4 and Map3k7* (Suppl. Figure S11).

These results suggest that the disease-specific CM3/CM5 subpopulations represent cardiomyocytes with damaged nuclei. Indeed, despite the transient nature of NE rupture^11,66,67^, we captured these events through co-staining for the cardiomyocyte marker PCM1, RTN4 (representing CM3/CM5 populations), and BAF (representing nuclei with NE rupture) in cKO hearts. RTN4-positve cells were highly enriched for BAF-positive nuclei (64% compared to 17% in RNT4-negatve cells), indicating a strong association between NE rupture and increased RTN4 levels (Figure 3H-I). In addition to TLR-associated signaling, we found evidence of STING-independent cytosolic DNA sensing in CM3 and CM5 cardiomyocytes, including upregulation of *Dhx9* (Suppl. Figure S8E-F), a helicase known to bind to DNA and activate NF-κB and TGF-β signaling^68^, and *Hmgb1* (Suppl. Figure S11), which binds to cytosolic DNA, acting as a TLR accessory protein^68^. Furthermore, many cardiomyocytes, particularly in the CM3 subpopulations, expressed interferon gamma inducible protein 16 (*IFI16*, also known as *Ifi204*) (Suppl. Figure S8G-H), a cytoplasmic DNA sensor that can work independently of STING to induce formation of inflammasome complexes and release of cytokines^69^.

Collectively, these findings suggest that the disease-specific subpopulations CM3/CM5 represent cardiomyocytes in which NE rupture triggers activation of cytosolic DNA-sensing pathways, leading to the production of cytokines, interferons, and other inflammatory signaling pathways. Supporting this interpretation, cardiac samples from patients with *LMNA*-DCM show similar upregulation of several genes involved in PRR, including *RTN4*, *MAP3K7*, and *HMGB1*, in comparison to healthy controls (Suppl. Figure S12).

### cKO hearts show progressive innate immune cell activation

Cytosolic DNA-sensing pathways can activate transcription factors such as NF-κB and IRF, leading to their nuclear translocation and subsequent upregulation of pro-inflammatory cytokines and type I interferons, including INF-α^17^ and INF-γ^70,71^. In parallel, studies on patient samples and murine *LMNA*-DCM models^38,48^ found that in response to injury cardiomyocytes release cytokines and damage-associated molecular patterns (DAMPs), which recruit immune cells and promote inflammation and cardiac remodeling^72,73^. Consistent with these previous findings, our analysis of GO cluster terms among the eDEGs and 25-dpi DEGs revealed an enrichment for immune response and inflammation (Figure 2D), particularly associated with innate immunity. The disease-specific CM3 and CM5 subpopulations had significantly elevated expression of cytokines (*Il34*, *Il4*, *Cxcl9*, *Ccl5*) and several DAMPs, including heat shock proteins, S100 proteins, and HMGB-1 (Suppl. Figure S13). Several of these DAMPs also appeared as eDEGs, suggesting their potential involvement in disease promoting processes.

Our snRNA-seq analysis further revealed an increase in immune cell populations within the cKO hearts compared to cWT controls at 25-dpi (Figure 4A). We confirmed an increased prevalence of immune cells (marked by the pan-leukocyte marker CD45) in cKO hearts at 25-dpi (Figure 4B). Levels of macrophages and monocytes (marked by CD68) were particularly elevated, starting at 11-dpi and progressively increased (Figure 4C), consistent with the detection of many innate immunity related eDEGs (Fig. 2D) and confirmed by western blot analysis (Suppl. Figure S10C). The immune subtypes however lacked disease-specific subpopulations but instead displayed an expansion of existing immune subpopulations in cKO hearts during disease progression, with particular shifts towards myeloid subpopulations Mye1 (22% to 32%), Mye2 (24% to 28%), and Mye4 (13% to 23%), and natural killer/T cell subpopulation NKT2 (29% to 51%) (Figure 4A, Suppl. Figure S14A). Using established gene expression markers^74^, we identified Mye2 & Mye4 as tissue resident macrophages and Mye1 as infiltrating macrophages (Figure 4D). Our data indicates that both recruitment and expansion of tissue-resident immune cells drive the evolving immune landscape, potentially amplifying inflammatory signaling and disease progression. Co-staining for the monocyte/macrophage marker CD68 and the CM3/CM5 marker RTN4 revealed increased spatial proximity between regions expressing RTN4 and CD68-positive regions in the cKO hearts compared to cWT controls (Figure 4E-F), supporting a model in which RTN4-mediated release of cytokines from cardiomyocytes with NE rupture leads to the local recruitment/expansion of innate immune cells to drive the inflammation and remodeling observed in *LMNA*-DCM.

**Figure 4:**
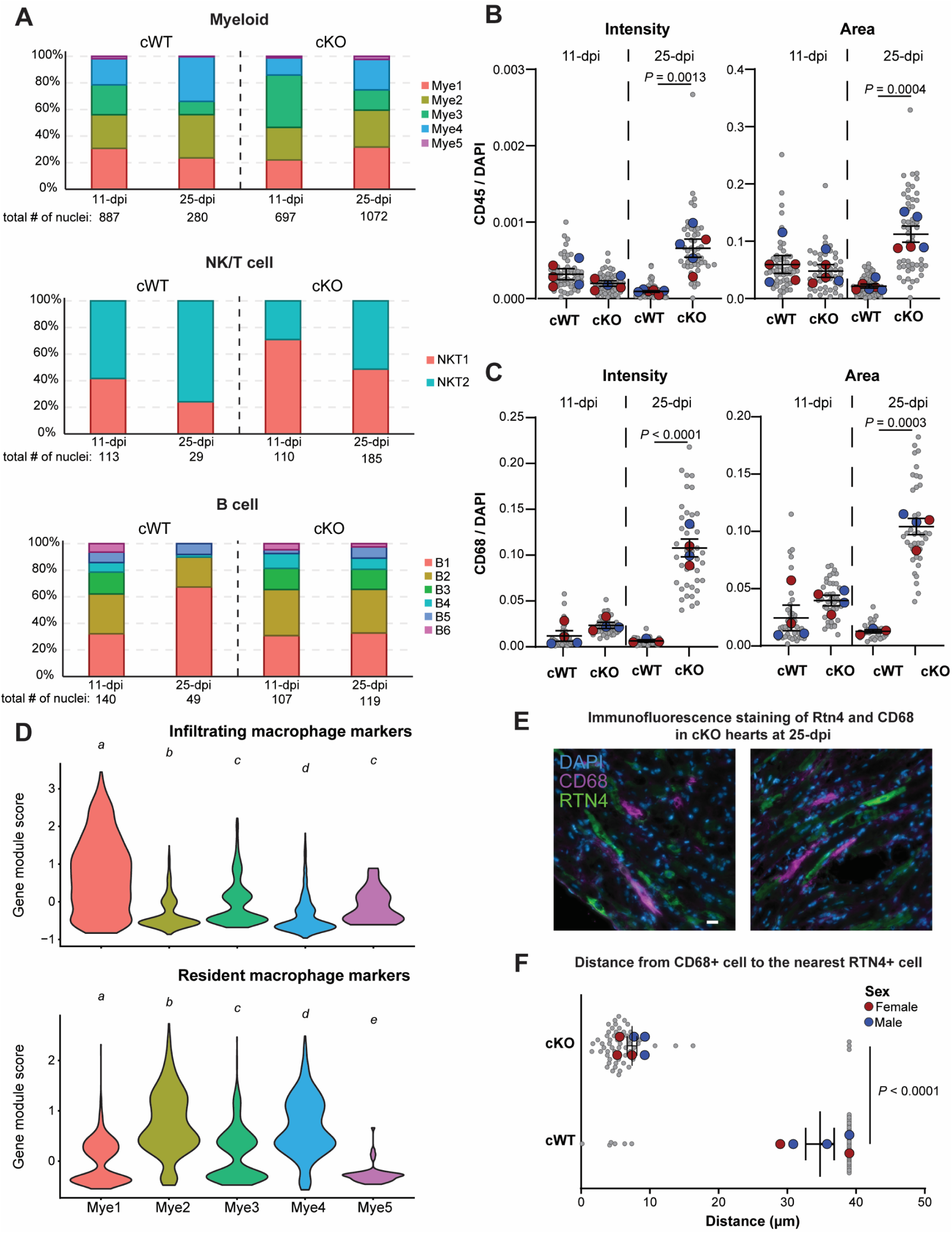
Cytoplasmic DNA sensing initiates immune infiltration in cKO hearts. **(A)** Number of nuclei captured in each genotype within individual immune cell subclusters and in total. Quantification of **(B)** CD45 and **(C)** CD68 fluorescence intensity normalized to DAPI intensity (left) and area coverage normalized to DAPI area coverage (right) for cKO and cWT cardiac tissue sections, collected at 11-dpi (left side of each graph) and 25-dpi (right side of each graph). Larger colored circles represent average values for individual male (blue) or female (red) mice. Smaller gray circles represent individual fields of view. *N* = 5-6 per genotype for CD45 and *N* = 4 animals per genotype for CD68. Statistical analysis by Tukey Honest Significant Difference of two-way ANOVA test. **(D)** Gene module scores for infiltrating (left) and resident (right) gene markers across individual myeloid subpopulations in the integrated snRNA-seq dataset. Groups that share a letter are not significantly different; different letters indicate *P* < 0.05, pair-wise Wilcoxon rank sum test with Benjamini-Hochberg *p*-value adjustment. **(E)** Immunofluorescence images of cKO cardiac tissue sections at 25-dpi stained for CD68 (magenta), RTN4 (green), and DAPI (blue). Scale bar: 20 µm. **(F)** Nearest neighbor analysis showing distances (µm) between RTN4-positive and CD68-positive regions in cKO (top) and cWT (bottom) hearts. Larger colored circles represent average values for individual male (blue) or female (red) mice. Smaller gray circles represent individual fields of view. *N* = 5-6 animals per genotype. For statistical comparison, we selected 40 µm as an arbitrary threshold for lack of spatial proximity. Statistical analysis by Tukey Honest Significant Difference of one-way ANOVA test.

Concurrently, we observed a small but statistically significant increase in *Lmna* transcript levels in cKO hearts between 11- and 25-dpi (*P* = 0.042, Suppl. Figure S3B). To determine whether this increase was due to the infiltration/expansion of immune cell (Figure 4A-D) or the loss of cardiomyocyte with low lamin A/C levels, we quantified the percentage of cardiomyocytes (PCM1-positive) amongst all detected nuclei in cardiac tissue at 11- and 25-dpi, as well as their lamin A levels, by immunofluorescence. Neither the fraction of PCM1-positive nuclei (Suppl. Figure S3C) nor the lamin A levels in cardiomyocytes (Suppl. Figure S3D) changed from 11- to 25-dpi, indicating that cardiomyocyte composition and lamin A expression remained unchanged. These findings support the interpretation that the slightly elevated *Lmna* expression in bulk tissue likely reflects infiltration/expansion of immune cells (Figure 4A-D).

### Lamin A/C depleted cardiomyocytes affect non-cardiomyocyte gene expression through altered cell-cell communication

Inflammatory cytokines and DAMPs lead to activation of immune cells^75^ and myofibroblasts^76,77^, thereby inducing further transcriptomic changes. We reasoned that non-cardiomyocyte cells would thus display disease-specific transcriptomic signatures in response to cell-cell signaling initiated by pathogenic cKO cardiomyocytes. Our snRNA-seq analysis identified several subpopulations in cKO non-cardiomyocyte cells with distinct gene expression changes (Suppl. Figure S14A-C), despite not being depleted for lamin A/C. In agreement with the general transcriptomic trend (Figure 2B), fibroblasts had substantially higher aberrant gene expression between cKO and cWT hearts at the 25-dpi (Figure 5A). In contrast, over 65% of 25-dpi DEGs in immune cells were already differentially regulated at 11-dpi (Figure 5A). These findings suggest that the contribution of fibroblasts to pathogenic gene expression takes place at an advanced disease stage, whereas immune cells gene misregulation occurs early in disease.

**Figure 5:**
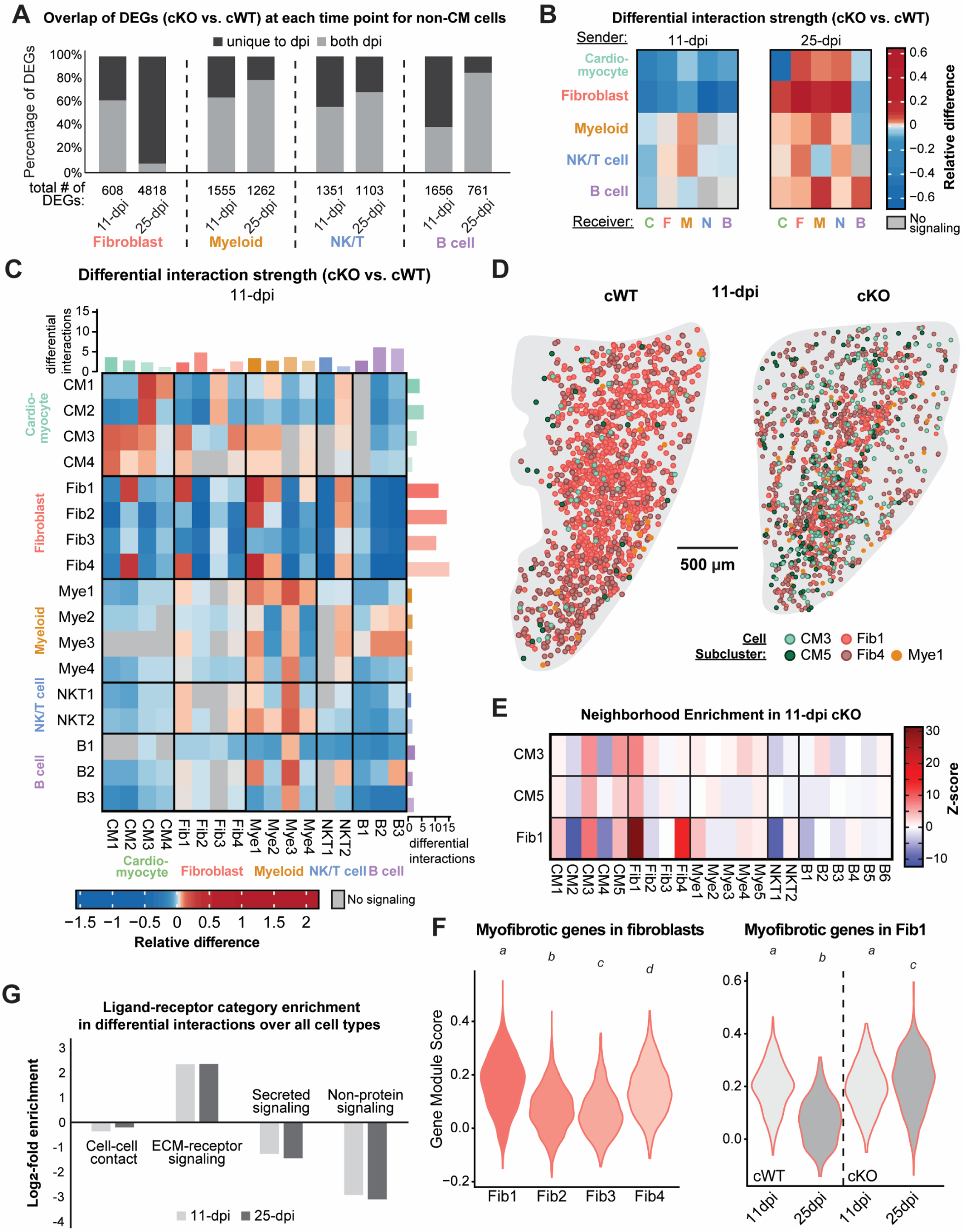
Prediction of intercellular signaling in cKO cardiomyocytes, fibroblasts, and myeloid cells reveals a fibrosis-inflammation link driven by disease-specific cardiomyocyte subpopulations. **(A)** Percentage of differentially expressed genes (DEGs) between cKO and cWT hearts at 11- and 25-dpi, for individual non-cardiomyocyte cell types (separated by dotted line). DEGs unique to (light gray) or shared between (dark gray) time points are denoted. The total DEG counts are indicated under each column. **(B-C)** Heatmap showing relative difference (cKO versus cWT) in the interaction strength of sender (rows) and receiver (columns, initial letter) cell types **(B)** and subpopulations **(C)** at 11- (left) and 25-dpi (right). **(C)** The number of differential interactions for each row and column are indicated in the bar plot. Subpopulations with insufficient nuclei for analysis were removed (i.e., CM5, Mye5, and B4-B6). **(D)** Spatial transcriptomics annotation of cardiomyocyte (CM3, light green; CM5 dark green), fibroblast (Fib1, pink; Fib4 maroon), and myeloid (Mye1, orange) subpopulations in cWT and cKO 11-dpi ventricles (tissue boundaries shown in gray) from heart slices. **(E)** Heatmap of *z*-score neighborhood enrichment for subpopulations in 11-dpi cKO ventricles based on spatial transcriptomics. **(F)** Gene module scores for a myofibrotic gene set in fibroblast subpopulations (left) and split by experimental condition for Fib1 (right, light gray - cWT, dark gray - cKO). Groups that share a letter are not significantly different; different letters indicate *P* < 0.05, Kruskal–Wallis tests followed by pairwise Mann–Whitney U tests with Benjamini-Hochberg *p*-value adjustment. **(G)** Log_2_-fold enrichment of particular ligand-receptor categories associated with differential interactions over all cell types at 11-dpi (light gray) and 25-dpi (dark gray).

Transcriptomic changes in non-cardiomyocyte cKO cells could either be the result of communication from the Lamin A/C-depleted cardiomyocytes or in response to global changes in the failing heart. To distinguish these mechanisms, we applied the CellChat^34^ algorithm to our snRNA-seq data to compare cell-cell communication between cKO and cWT hearts. Cell-cell communication was overall downregulated in cKO versus cWT hearts at 11-dpi but increased by 25-dpi (Figure 5B, Suppl. Figure S15). Fibroblasts had a particularly strong shift from down- to up-regulated interaction strength between the 11-and 25-dpi, mirroring the DEG increase over time (Figure 5A) and driving most of the outgoing signaling to innate immune cells (myeloid and NK/T cells). Myeloid and NK/T cell interactions were also upregulated at 25-dpi, further corroborating the involvement of innate immunity in disease progression. In contrast, B cells had decreased incoming communication from all other cell types, at both 11- and 25-dpi. Other cardiac cell types, including endothelial and mural cells, similarly exhibited reduced cell-cell communication with cardiomyocytes in cKO hearts at both 11- and 25-dpi (Suppl. Figure S15). This diminished interaction, particularly at 25-dpi when pathological cell-cell communication is expected to peak, suggested that these cell types do not substantially contribute to pathogenic signaling.

Whereas overall cell-cell communication was increased in cKO hearts at 25-dpi, cardiomyocyte-to-cardiomyocyte communication strength was markedly reduced in cKO versus cWT hearts at both timepoints (Figure 5B). However, CellChat analysis on individual subpopulations at 11-dpi (Figure 5C, Suppl. Figure S16) revealed that this overall reduction was primarily driven by the larger CM1 and CM2 subpopulations. In contrast, in cKO hearts, the disease-specific subpopulation CM3 had increased communication with other cardiomyocytes, as well as with myeloid subpopulations Mye1 and Mye 2 and fibroblast subpopulations Fib1 and Fib4 (Figure 5C). Although the CM5 subpopulation contained too few nuclei in the 11-dpi cWT sample to assess differential cell-cell communication, CellChat predicted interactions between CM5 and the Fib1, Mye1, and Mye2 subpopulations in the cKO samples, similar to the findings for the CM3 subpopulation (Suppl. Figure S16). Additionally, both Fib1 and Fib4 showed increased interactions with Mye1 and Mye2 in cKO hearts. This finding was further supported by our spatial transcriptomic analysis, which revealed increased spatial proximity among CM3, CM5, Fib1, Fib4, and Mye1 subpopulations in cKO compared to cWT hearts (Figure 5D-E, Suppl. Figure S17). Notably, Fib1 and Fib4 showed a particularly pronounced shift in spatial distribution at 11-dpi (Figure 5D).

Previous studies had reported the appearance of activated fibroblast (myofibroblast), primarily responsible for extracellular matrix (ECM) deposition during fibrosis, in *LMNA*-DCM hearts^78^. We determined that the cKO Fib1 fibroblast subpopulation had an expression profile that most closely matched that of myofibroblasts (Figure 5F, left), which significantly intensified over time (Figure 5F, right). Pathogenic activation of fibroblast in cKO hearts was further supported by the CellChat prediction that cardiomyocyte and non-cardiomyocyte differential interactions were largely mediated by ECM-receptor pathways (Figure 5G). These results are in agreement with recent work in another cKO mouse model, which found that cardiomyocytes activate fibroblasts via ECM-mediated signaling^48^, and suggest that fibrosis is not only a downstream consequence of cardiac dysfunction in *LMNA*-DCM but directly contributes to the disease pathogenesis

The increased cell-cell communication (Figure 5C) and spatial proximity (Figure 5D-E) between CM3, CM5, Fib1, Fib4, and Mye1 in cKO versus cWT hearts at 11-dpi suggests a pathogenic communication network between disease-specific CM3/CM5 cardiomyocytes, myofibrotic fibroblasts, and infiltrating macrophages. Collectively, our snRNA-seq, spatial transcriptomic, and immunofluorescence data indicate that cardiomyocyte and non-cardiomyocyte interactions are becoming reorganized in cKO hearts, starting in the early disease stage and progressing over time, leading to the inflammatory and fibrotic phenotypes observed in *LMNA*-DCM hearts.

### Reducing NE damage in lamin A/C-deficient cardiomyocytes partially restores gene expression and substantially ameliorates *LMNA*-DCM pathology

Our data suggest that inflammatory responses triggered by NE rupture of mechanically weaker cKO cardiomyocyte nuclei drive altered gene expression and pathogenesis. Thus, we hypothesize that reducing NE rupture should reverse many transcriptional changes—even in the continued absence of lamin A/C—and ameliorate disease progression. To test this hypothesis, we generated mice with inducible, cardiomyocyte-specific expression of a dominant negative nesprin-construct, consisting of the Klarsicht, ANC-1, Syne Homology (KASH) domain of nesprin-2 (DN KASH)^79^, and concurrent depletion of lamin A/C (Figure 6A). Expression of DN KASH displaces endogenous nesprins from the NE, resulting in LINC complex disruption and reducing cytoskeletal forces acting on the nucleus^9,10,19–22^. LINC complex disruption has previously been shown to reduce NE rupture and DNA damage in lamin A/C-deficient and *Lmna* mutant myoblasts^9^ and isolated cardiomyocytes^10^. Tamoxifen treatment of adult mice resulted in a significant reduction in the fraction of cKO cardiomyocytes with NE ruptures (BAF-positive nuclei) at 11-dpi (Figure 6B, Suppl. Figure S18). Cardiomyocyte-specific LINC complex disruption dramatically improved cardiac function in cKO animals (Figure 6C, Suppl. Figure S19). Although cardiac function was not rescued completely, cKO mice expressing DN KASH had near-normal lifespan, with survival extended from ∼30 dpi to over 400-dpi, which served as our experimental endpoint. cWT animals with cardiac LINC complex disruption had normal survival (Figure 6D).

**Figure 6:**
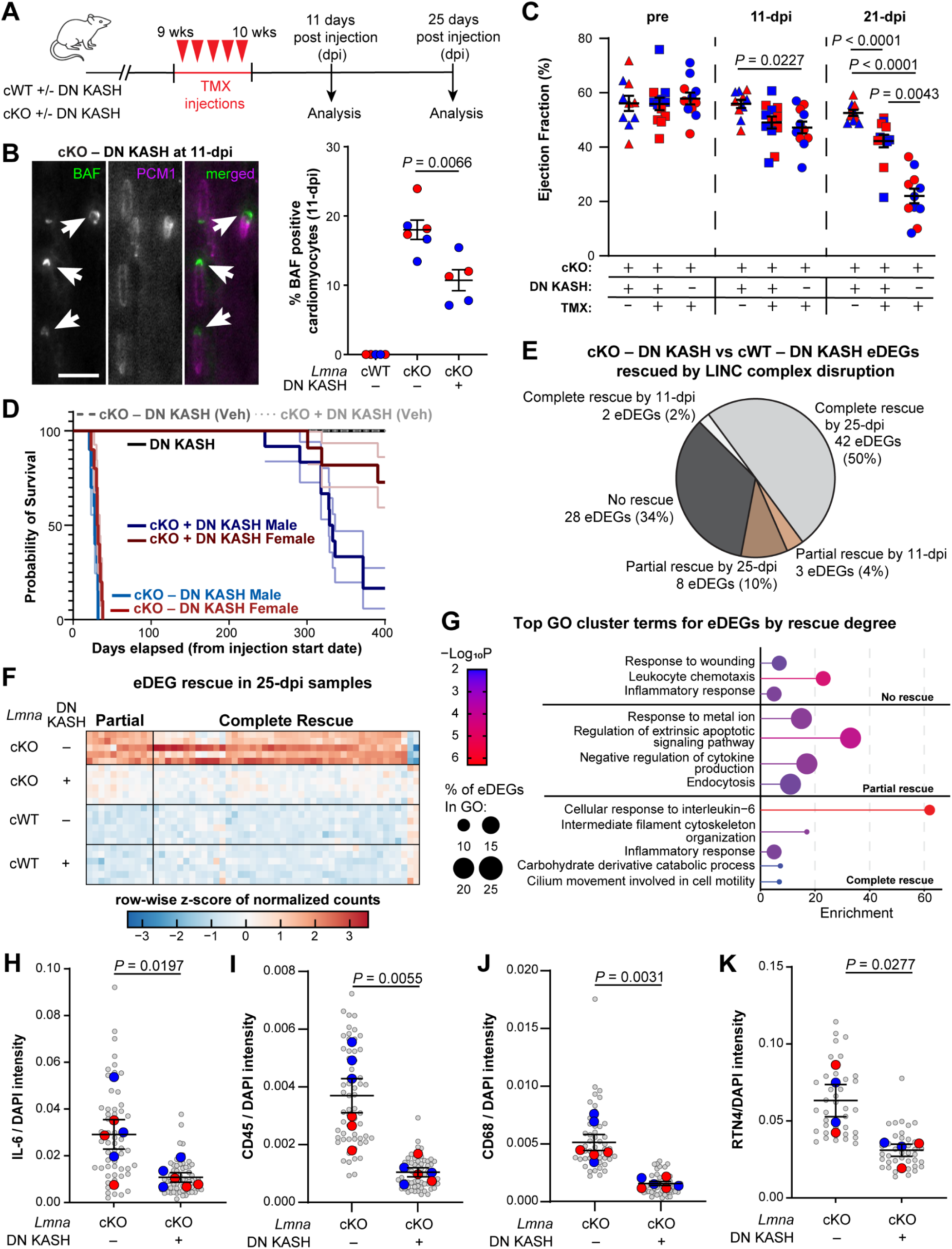
LINC complex disruption dramatically improves cardiac function and survival in cKO mice by reducing NE rupture and transcriptional activation of cytosolic DNA sensing-mediated inflammation. **(A)** Experimental timeline for concurrent induction of *Lmna* deletion and LINC complex disruption. Echocardiography and tissue collection were performed at the indicated time points. **(B)** Representative images of BAF-positive (green) cardiomyocytes (PCM1-positive, magenta) in cKO mice without LINC complex disruption (left) and quantification for cWT and cKO mice with (+ DN KASH) and without LINC complex disruption (right). Arrows indicate representative cardiomyocytes. Larger colored circles represent average values for individual male (blue) or female (red) mice. *N* = 5-6 animals per genotype. **(C)** Left ventricular ejection fraction (EF) measured before tamoxifen (TMX) injection (pre), or at 11-dpi and 25-dpi. Symbols represent %EF of individual male (blue) or female (red) mice. *N* = 10-12 animals per genotype. Error bars represent mean ± S.E.M. **(D)** Kaplan-Meier survival plot for cWT mice with LINC complex disruption (cWT + DN KASH, *10* = 5 males and 5 females), cKO mice treated with vehicle (cKO vehicle, *10* = 5 males and 5 females), cKO mice with DN KASH treated with vehicle (cKO + DN KASH vehicle, *10* = 5 males and 5 females), cKO mice with LINC complex disruption (cKO + DN KASH, *N* = 23 mice: 12 males and 11 females), and cKO mice (cKO, *N* = 20 mice: 10 males and 10 females). **(E)** Fraction of eDEGs with complete (44 eDEGs, light gray), partial (11 eDEGs, brown), or no rescue (28 eDEGs, dark gray) following LINC complex disruption. **(F)** Heatmap of row-wise *z*-score normalized read counts for eDEGs rescued by LINC complex disruption, grouped by degree of rescue (complete, right; partial, left) and sample condition. **(G)** Gene Ontology (GO) cluster terms enriched of eDEGs stratified by rescue category (none, top; partial, middle; complete bottom). Dot color indicates enrichment significance and size reflects proportion of eDEGs per term. **(H-K)** Quantification of IL-6 **(H)**, CD45 **(I)**, CD68 **(J)**, and RTN4 **(K)** intensity normalized to DAPI in cKO with (+ DN KASH) and without (– DN KASH) LINC complex disruption. Larger colored circles represent average values for individual male (blue) or female (red) mice. Smaller gray circles represent individual fields of view. *N* = 6 animals per genotype for IL-6, CD45, and CD68. *N* = 4 animals per genotype for RTN4. Statistical analysis performed by two-sided *t*-test.

To gain additional insights into the effects of LINC complex disruption, we conducted bulk RNA-seq on left ventricular tissue from cKO and cWT mice, each with and without concurrent cardiomyocyte-specific LINC complex disruption. We decided to use bulk RNA-seq to efficiently capture global transcriptional changes across the left ventricle with higher sensitivity and better gene coverage than offered by snRNA-seq and spatial approaches. We first applied our bulk RNA-seq analysis pipeline to cKO and cWT littermates lacking the DN KASH transgene (Suppl. Figure S20A). We found a progressive increase in DEGs from 11-dpi (*n* = 208) to 25-dpi (*n* = 3426) (Suppl. Figure S20A), similarly to our original cKO vs. cWT analysis (Figure 2B-C, Suppl. Figure S2A). Cross-referencing gene expression differences between cKO and cWT hearts lacking DN KASH expression at 11- and 25-dpi, identified 83 eDEGs (Suppl. Figure S20A-C). The identified eDEGs were consistently enriched for GO terms involved in inflammation, apoptosis, and innate immunity (Suppl. Figure S20D), mirroring our findings in the original cKO cohort (Figure 2). This convergence supports the view that even modest early transcriptional shifts reliably predict meaningful disease-relevant processes.

Next, we assessed the effect of LINC complex disruption on cardiac gene expression. DN KASH expression caused only minimal transcriptional changes in cWT hearts (4 DEGs detected by 25-dpi), indicating that transcriptional changes observed in cKO mice are likely disease-related and that LINC complex disruption in adult mice does not cause adverse effects. In the cKO hearts, DN KASH expression led to the majority of eDEGs (*n* = 44, 53%) and DEGs (*n* = 1786, 52%) returning to levels statistically indistinguishable from cWT controls (Figure 6E-F, Suppl. Figure S21). We designated this set of genes ‘complete rescue’. We identified an additional 11 eDEGs (13%) and 226 DEGs (7%) as ‘partial rescue’, i.e., their expression significantly differed between cKO samples with and without LINC complex disruption but did not return completely to cWT levels (Figure 6E-F, Suppl. Figure S21). A small fraction of eDEGs and DEGs were rescued already at 11-dpi, while the majority of eDEGs and DEGs reached rescue levels at 25-dpi (Figure 6E; Suppl. Fig. S21).

GO term analysis of eDEGs showed that the “partial” and “complete” rescued gene sets were enriched for apoptosis- and inflammation-related terms, whereas terms such as “inflammatory response” and “leukocyte chemotaxis” were still present in the “no rescue” group (Figure 6G). The persistence of some GO terms associated with inflammation likely reflects residual NE damage in cKO mice despite DN KASH expression (Figure 6B), the multifactorial nature of inflammatory signaling, and transcriptional changes inherent to lamin A/C depletion. Nonetheless, LINC complex disruption substantially mitigated key pathological drivers, as evidenced by the rescue of several key genes involved in cytosolic DNA sensing, including *Rtn4*, *Ifi204*, *Tlr9*, *Il6*, *Irf7*, and *Hspa1a* (Suppl. Figure S22), and a significant decrease in the protein levels of the leukocyte marker CD45, the macrophage/monocyte marker CD68, the inflammatory cytokine IL-6, and RNT4 in cKO hearts expressing DN KASH (Figure 6H-K). Our transcriptomic analysis further revealed that 47% of infiltrating macrophage-associated genes and 65% of myofibroblast-associated genes were completely or partially rescued to wild-type levels in cKO hearts with LINC complex disruption (Suppl. Figure S23).

Supporting a strong connection between inflammatory responses and disease pathology, we observed an inverse correlation between the immune cell burden (presence of CD45-and CD68-positive cells) and ejection fraction in cKO mice across all conditions (Suppl. Figure S24). In further support of this interpretation, “cellular response to interleukin-6” showed the strongest enrichment among GO terms that were completely rescued following DN KASH expression (Figure 6G, bottom), and “negative regulation of cytokine production” and “endocytosis” were part of the GO terms enriched in the ‘partial rescue’ gene sets (Figure 6G). Together, these findings suggest that the therapeutic benefits of LINC complex disruption result from reduced NE rupture and downstream signaling, such as activation of PRR pathways and the production of inflammatory cytokines, thereby limiting immune cell recruitment and other pathogenic processes.

## Discussion

We generated a novel mouse model for *LMNA*-DCM with inducible, cardiomyocyte-specific lamin A/C depletion to gain further insights into the pathogenic mechanisms responsible for the cardiac phenotypes. Using a comprehensive, temporally resolved transcriptomic analysis of cardiac tissue that combines bulk, single-nucleus, and spatially resolved RNA-seq, we characterized dynamic changes in gene expression across individual cell subpopulations to determine which transcriptional programs may initiate or propagate pathology. Finally, due to the high prevalence of NE rupture in lamin A/C-depleted cardiomyocytes, we tested whether LINC complex disruption, which reduces mechanical stress on muscle nuclei, could rescue transcriptional and functional changes in the cKO hearts.

We identified two disease-specific cardiomyocyte subpopulations that were largely responsible for the altered expression of disease-associated genes, termed eDEGs (Figure 2C, Suppl. Figure S2A). These eDEGs revealed a previously unrecognized role of cytosolic PRR signaling that operates independently of cGAS. Generalizing our findings to human *LMNA*-DCM, we found evidence of cytosolic PRR signaling activation in cardiac samples from both *LMNA*-DCM and non-laminopathy DCM patients^38^ (Suppl. Figure S12). Because human samples capture only end-stage disease signatures, these data alone cannot determine the timing and causality of PRR activation in human DCM. In contrast, the time-resolved nature of our cKO model supports PRR signaling as a potential mechanism of DCM pathology.

In the cKO hearts, the disease specific cardiomyocyte subpopulation (i.e., CM3 and CM5) drove many of the transcriptional changes despite representing only a small fraction of all captured cardiomyocytes (CM3: 5.72%; CM5: 5.3%). This fraction of cardiomyocytes is slightly smaller in magnitude than the fraction of cardiomyocytes undergoing NE rupture observed in our study (18%; Figure 6B) and others, which report up to 30% of cardiomyocytes with damage nuclei^10,48^. Two factors likely contribute to this discrepancy: (1) severely damaged cardiomyocyte nuclei may be selectively lost during processing for snRNA-seq, and (2) unbiased clustering discretizes what is in fact a transcriptional continuum, potentially causing a subset of diseased cardiomyocytes to be grouped into subpopulation CM4, which sits between CM3 and CM5 in the transcriptional space. Indeed, pseudo-time analysis based on transcriptomic data (Suppl. Figure S25A), together with the shared enrichment of oxidative phosphorylation in CM4 and CM5 (Suppl. Figure S25B), indicate that CM5 cardiomyocytes likely emerge from the CM4 subpopulation. In cKO hearts, this is reflected by a decline in CM4 cells as the CM5 cluster increases over time (Figure 2G), and by CM4 cells located at the CM4-CM5 interface exhibiting CM5-like transcriptional signatures (Suppl. Figure S25C). In contrast, in cWT hearts, CM4 increases with age (Figure 2G), and its oxidative phosphorylation signature (Suppl. Figure S25B left) is consistent with a physiological cardiomyocyte maturation trajectory following the metabolic shift from glycolysis^80^. Thus, the CM4 subpopulation plays a dual role: in disease, it feeds into the CM5 subpopulation as part of a NE rupture-associated progression, whereas in cWT hearts it expands as a maturing cardiomyocyte state. This continuum view also explains why CM4 cells show eDEG signatures similar to subpopulations CM3 and CM5 at 25 dpi (Figure 2I). Indeed, at 11-dpi, CM3, CM4 and CM5 together constitute ∼30% of the cardiomyocytes, closely matching the fraction of ruptured nuclei in previous studies in similar *LMNA*-DCM models^10,48^.

Although previous studies had reported increased NE rupture in *Lmna*-null and mutant cardiomyocytes^10,45,48^ and that LINC complex disruption improves cardiac function in *LMNA*-DCM mouse models^10,81^, we present the first evidence mechanistically linking NE rupture to altered gene expression in *LMNA*-DCM. We demonstrate that LINC complex disruption reduces NE rupture in lamin A/C-depleted cardiomyocytes (Figure 6B) and results not only in near-complete rescue of cardiac function and survival (Figure 6C-D) in cKO mice, but also rescues over 50% of misregulated genes, despite the continued depletion of lamin A/C. These findings support a role for NE rupture as an early catalyst of disease pathology in *LMNA*-DCM. Roughly 66% of the detected eDEGs (Figure 6E), which appeared downstream of NE rupture, were rescued by LINC complex disruption, suggesting that the ‘gene regulation hypothesis’ and the ‘structural hypothesis’^3,82^ are not mutually exclusive, and are at least in part mechanistically linked. At the same time, the broad impact of NE rupture on cardiac dysfunction may also explain why targeting specific pathways downstream of nuclear damage, such as p53 or STING signaling, has provided only limited success in preclinical *LMNA*-DCM models^83–85^.

It is worth noting that DN KASH acts as a competitor to endogenous KASH-domain containing nesprins, and is thus unlikely to achieve 100% LINC complex disruption in vivo. This incomplete disruption may explain why 10% of cardiomyocyte nuclei still undergo rupture (Figure 6B) and rescue of gene expression is incomplete (Figure 6D).

Alternatively, the latter could also be explained by the fact that *LMNA* mutations can alter the epigenome, including at LADs and at DNA methylation sites^86,87^. These epigenetic changes were not assessed as part of this study but likely contribute to the residual gene expression changes in the cKO hearts expressing DN KASH. Nevertheless, the resulting reduction in inflammation, normalization of numerous transcriptional changes, and improvement in cardiac function are sufficient to support survival comparable to wild-type mice. The minimal transcriptional effect of DN KASH expression in the cWT background indicates that DN KASH itself does not broadly reprogram gene expression and is well tolerated, consistent with previous studies^79,88^, supporting our conclusions that reducing NE rupture is a key driver for the observed rescue in cKO mice. However, we cannot exclude the possibility that LINC complex disruption may also cause changes in chromatin organization, epigenetic modifications^89^, or the cytoskeleton^10^, which could further contribute to the rescue.

Our study revealed activation of innate immunity in the cKO hearts, based on both immunofluorescence labeling (Figures 4B-C) and transcriptomic analyses (Figures 2D, and 4D). As most cardiomyocytes lack cGAS and STING expression, we hypothesize that the activation of innate immunity following NE rupture is instead mediated by other components of the cytoplasmic PRR pathway^90,91^, which we found to be upregulated in eDEGs (Figure 2D) and enriched in the disease-specific cardiomyocyte subpopulations CM3 and CM5 (Figure 3F-I; Figure 4E-F; Suppl. Figure S11). At the same time, DNA released by damaged cardiomyocytes and subsequently taken up by other cells might further activate cytosolic PRR pathways in non-cardiomyocyte cell types, such as macrophages and fibroblasts. Our integration of single-nucleus and spatial transcriptomics demonstrates active cell-cell communication between these disease-specific cardiomyocyte subpopulations, fibroblasts, and myeloid cells. These effects, complemented by cytokine and DAMP releases downstream of cytoplasmic PRR pathways, could then further contribute to pathogenic processes in fibroblasts and immune cells.

Our computational approach, which integrates early and late timepoints to define eDEGs, robustly identified disease-driving transcriptional programs, including PRR signaling, inflammation, and innate immune activation across multiple independent experiments. In the interpretation of the eDEGs, however, it is important to recognize that early transcriptional changes are inherently subtle and therefore more susceptible to biological noise, such as stochasticity in the prevalence of cell subpopulations (e.g., cardiomyocytes with NE rupture or infiltrating immune cells) that affect population level measurements, mouse cohort-specific variability, including the genomic insertion of the Cre-inducible DN KASH transgene, and differences in sequencing platforms. This biological and technical variability is reflected in part in the limited overlap in log2 fold-change directionality (cKO vs. cWT) between genes with *p*-adj < 0.05 at the early disease stage (11-dpi) and the DEGs detected at late disease stages (25-dpi) in the experiments with the DN KASH rescue (Suppl. Figures S20A, and S20C). This reduced directional concordance was largely due to transcriptional variability in the early (11-dpi) cKO condition (Suppl. Figure S26A-B) and accounted for the smaller number of eDEGs in the DN KASH rescue experiment (Suppl. Figures S2A and S20A). Nonetheless, even the genes with opposite directionality showed that DN KASH expression led to normalization of gene expression (Suppl. Fig. 20B), similar to the results observed in the identified eDEGs (Suppl. Fig. 20E). Later-stage transcriptional profiles converge strongly across experiments, both globally (Suppl. Figure S26B) and at the level of eDEGs (Suppl. Figure S26C). Crucially, despite the specific variations in the early timepoint transcriptional landscape, the underlying biological pathways implicated in both datasets were highly concordant. These observations underscore the value of prioritizing pathway-level consistency and increases of transcriptional signals with disease progression, over individual early genes in isolation. Together, our results indicate that the eDEG framework efficiently captures early transcriptional perturbations that mature into robust late-stage disease signatures and highlights pathways that are partially or fully rescued by DN KASH expression, reinforcing the biological relevance and predictive power of this approach.

### Limitations of the current study

Although the cKO model used in our study offers several advantages for identifying early *LMNA*-DCM disease drivers, it is important to note some inherent limitations of our model. The genetic landscape of human *LMNA*-DCM is diverse, encompassing *LMNA* missense mutations, frame-shift mutations, and splice-site variants spanning all exons and the first 10 introns of the *LMNA* gene^92^. Since *LMNA* mutations are congenital and nearly ubiquitously expressed across cell types, they may directly affect cardiac development and the function of non-cardiomyocyte cardiac cell populations, which is not reflected in our cardiomyocyte-restricted knockout model. Because non-cardiomyocyte cells in our model are not depleted for lamin A/C, and thus maintain normal cellular function^93,94^, we hypothesize that their contribution to disease progression may be disproportionally more extensive compared to constitutive mutations, leading to more severe and rapid disease onset. *LMNA* mutations in other tissues might likewise contribute to *LMNA*-DCM disease progression and severity. Indeed, while LINC complex disruption leads to an almost complete rescue of survival and cardiac function in the cKO model, it is only partial in models with global *Lmna* deletion or mutation^81^.

Our transcriptional analysis presented here focused on eDEGs and associated biological processes. This analysis, by design, excludes any DEGs identified at 25-dpi that were not statistically significant at the early disease timepoint (11-dpi) or that were inconsistent with the direction of change between the two timepoints. Although we assume that such genes primarily reflect the consequences of disease progression, rather than disease-driving mechanisms, they may nonetheless contribute to the disease.

Lastly, the goal of this study was to identify early transcriptional changes. We recognize that transcription levels do not always correlate with protein levels, nor does the transcriptional analysis capture post-translational modifications or changes in intra- and extracellular localization of proteins. Thus, although we have validated several key identified genes at the protein level and predictions from the transcriptional analysis align with our histological and functional measurements, further studies are necessary to fully validate the proposed mechanisms and to explore the suitability of the identified pathways as potential therapeutic targets.

### Conclusions

Our transcriptional analyses in cKO mice and littermate controls reveal that early activation of cytosolic PRR and ECM changes likely play major roles in driving *LMNA*-DCM pathology. We propose a model in which NE rupture in cardiomyocytes activates cytosolic PRR pathways, independent of cGAS/STING signaling, and, together with other effects triggered by loss of lamin A/C, leads to substantial transcriptional and functional changes in a subset of cardiomyocytes (Figure 7). Signaling from these cardiomyocytes, along with fibroblast-mediated ECM remodeling, potentially downstream of their own PRR activation, initiate a cascade of pathogenic cell-cell communication that recruits innate immune cells and promotes fibrosis. These immune cells induce inflammation via their own PRR signaling, further driving disease progression. The significant cardiac improvements in mice expressing DN KASH underscores the therapeutic potential of targeting the LINC complex or other strategies to reduce NE rupture in *LMNA* DCM. Besides LINC complex disruption, which currently requires genetic approaches and delivery via viral vectors that can trigger adverse immune reactions, our findings suggest that interventions aimed at restoring proper cell-cell-communication, and/or the aberrantly activated pathways immediately downstream of fragile lamin mutant nuclei, may offer new and promising strategies for treating *LMNA*-DCM, with potential relevance to other DCM forms that engage these same pathways.

**Figure 7:**
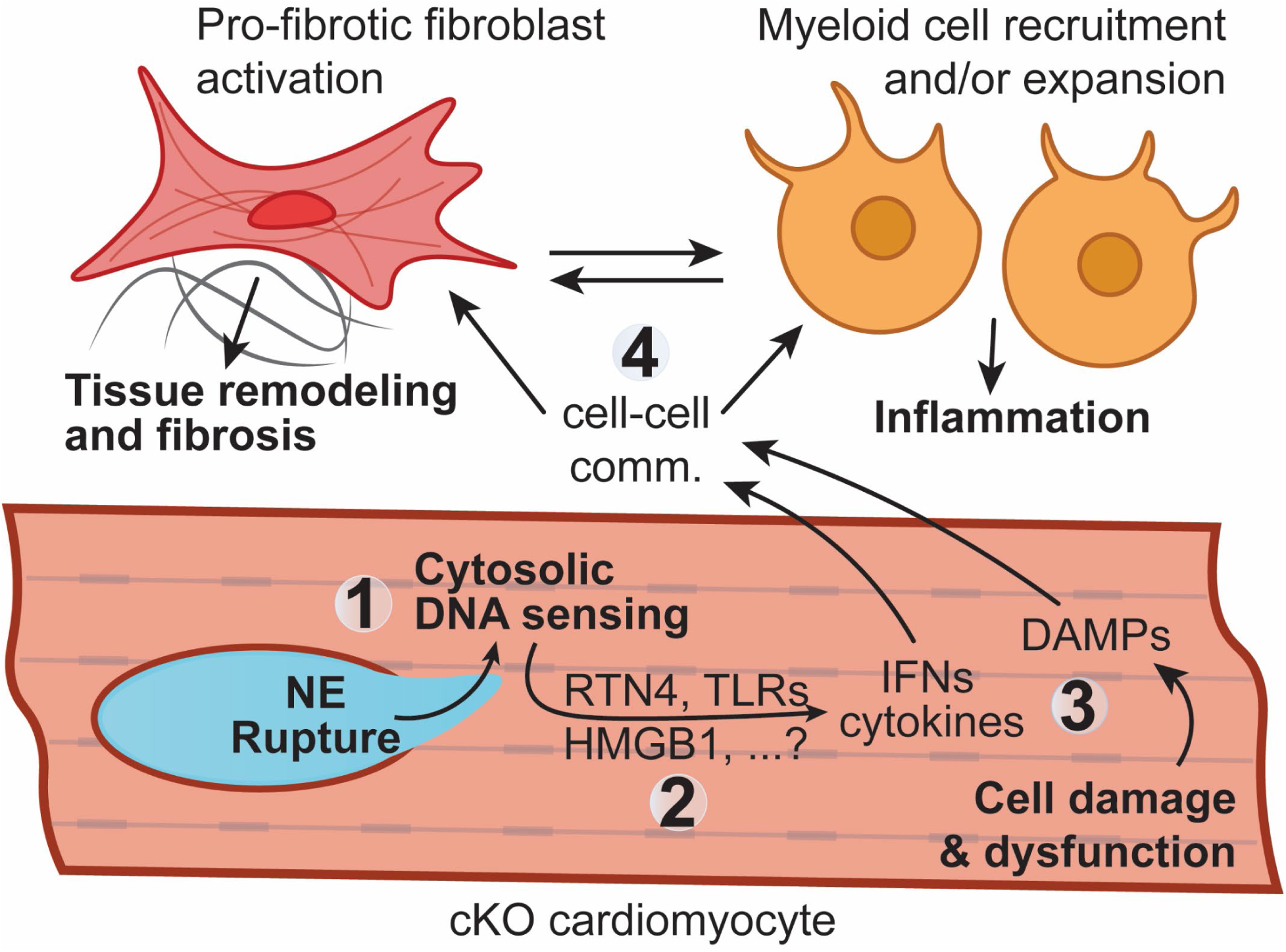
Cytosolic pattern recognition receptor signaling downstream of nuclear envelope rupture drives *LMNA*-DCM pathology. (1) Mechanical stress on the nuclear envelope (NE) leads to rupture in structurally compromised nuclei of cKO cardiomyocytes (notably subpopulations CM3 and CM5), resulting in exposure of nuclear DNA to the cytoplasm, either in the form of chromatin protrusions or cytoplasmic leakage of nuclear DNA fragments. **(2)** Cytosolic DNA is detected by cGas/STING-independent cytosolic nuclei acid sensors, such as Toll-like receptors (TLRs) regulated by RTN4 and HMGB1, thereby activating cytoplasmic pattern recognition receptor (PRR) signaling. **(3)** PRR activation triggers the production of interferons (IFNs) and cytokines by cKO cardiomyocytes. Concurrently, cardiomyocytes release damage-associated molecular patterns (DAMPs) as a consequence of cellular stress and injury. **(4)** These signaling molecules mediate aberrant cell-cell communication to fibroblast (Fib1) and myeloid cells (Mye1), promoting pro-fibrotic and inflammatory signaling cascades, respectively. Crosstalk between these fibroblasts and myeloid cells further amplifies fibrosis and inflammation in the heart, ultimately impairing cardiac function. LINC complex disruption attenuates this pathological cascade by substantially reducing the frequency of NE rupture events in cKO cardiomyocytes.

## Methods

### Animals

Mice expressing a transgene containing the mouse cardiac-specific α-myosin heavy chain promoter (αMHC or Myh6) directing expression of a tamoxifen-inducible Cre recombinase (MerCreMer) in juvenile and adult cardiac myocytes (αMHC-MerCreMer) and mice possessing loxP sites flanking exon 2 of the lamin A gene (*Lmna* floxed or *Lmna*^fl/fl^)^22^, were obtained from The Jackson Laboratory (JAX) in a C57BL/6 background (αMHC-MerCreMer: B6.FVB(129)-A1cf^Tg(Myh6-cre/Esr1*)1Jmk^/J, strain 005657; and *Lmna* floxed: *Lmna*^tm1.1Yxz^/J, strain 026284, respectively) and genotyped as instructed. Mice expressing a recombinant KASH domain (DN KASH) were generated by the Hodzic lab^79^. All mice used in this study were maintained in the same C57BL/6 genetic background and backcrossed for at least three generations. To generate the inducible, cardiac specific *Lmna* deletion mouse model (cKO), αMHC-MerCreMer and *Lmna* floxed mice were crossed to obtain mice hemizygous for αMHC-MerCreMer and homozygous for *Lmna* floxed (*Lmna*^fl/fl^). αMHC-MerCreMer hemizygous expressing wild-type *Lmna* (*Lmna*^+/+^) littermates served as controls (cWT). To generate the inducible, cardiac specific *Lmna* deletion and LINC complex disruption mouse model (cKO + DN KASH), αMHC-MerCreMer, DN KASH, and *Lmna* floxed mice were crossed to obtain mice hemizygous for αMHC-MerCreMer, hemizygous for DN KASH, and homozygous for *Lmna* floxed (*Lmna*^fl/fl^). Littermates hemizygous for αMHC-MerCreMer and homozygous for *Lmna* floxed (*Lmna*^fl/fl^), and mice hemizygous for αMHC-MerCreMer and expressing wild-type *Lmna* (*Lmna*^+/+^) without DN KASH expression served as controls (cWT – DN KASH). To induce lamin A/C depletion and/or LINC complex disruption in cardiomyocytes, tamoxifen (Cayman chemicals catalog#13258) was dissolved in sunflower seed oil (Sigma catalog# 88921) to a final concentration of 30 mg/kg (3 µl/g) and injected intraperitoneally at 9-10 weeks of age for five consecutive days, followed by a one-week-long washout period. Vehicle-only injections served as additional controls. For tissue collection, mice were anesthetized at the appropriate timepoints using Isoflurane (Covetrus catalog# 11695-6777-2). Immediately following cervical dislocation, hearts were harvested, washed with ice cold 1X Phosphate Buffer Saline (PBS), and processed for downstream analysis. All experiments, including breeding, maintenance, and euthanasia of animals, were performed in accordance with relevant guidelines and ethical regulations approved by the Cornell University Institutional Animal Care and Use Committee (IACUC protocol 2011–0099). Genotyping was performed by PCR using genomic DNA isolated from tail clippings following the protocol and sequencing primers suggested by JAX. Mice were housed in a disease-free barrier facility with 12/12 h light/dark cycles.

### Cardiac function

Mice were anesthetized with 2% isoflurane and placed on a stereotactic heated scanning base (37°C) attached to an electrocardiographic monitor. Left ventricular function was determined with a Vevo 2100 imaging system (VisualSonics) equipped with a MS550D transducer (22–55 MHz). Ejection fraction (EF%), fraction shortening (FS%), left ventricle diameter during systole (mm) and volume during systole (mm) were measured for at least five cardiac cycles using AM mode images. The average heart rate for all studied animals was 461 ± 24 beats per minute (BPM). Analysis was performed using the AutoLV Analysis Software (VisualSonics) and conducted by observers blinded to the mouse genotype and/or treatment groups.

### Immunofluorescence and histology of mouse tissues

Immediately following tissue collection, hearts were washed with 1X PBS and fixed in 4% paraformaldehyde (diluted in 1X PBS) at 4°C overnight. Samples were stored for up to one week in 70% ethanol (diluted in 1X PBS) at 4°C before being embedded in paraffin, sectioned, and stained with Masson’s Trichrome using standard methods at the Animal Health Diagnostic Center (College of Veterinary Medicine, Cornell University). For immunofluorescence analysis, samples were washed with PBS and flash frozen in Tissue-Tek® O.C.T. Compound (Sakura, #4583) in an ethanol bath, followed by storage at –70°C. Frozen tissue blocks were cryosectioned using an Epredia™ Microm HM525 NX Cryostat to a thickness of 5 µm, mounted on 25 × 75 × 1.0 mm Superfrost Plus Microscope slides (Fisherbrand® #12-55015) and left to air dry for 1 hour. Samples were stored at –20°C until processed for immunofluorescence staining and imaging. Prior to staining, slides containing tissue were equilibrated to room temperature and rehydrated with 1X PBS for 5 minutes. When using primary antibodies raised in mice, tissues were initially blocked with M.O.M® (Mouse on Mouse) Blocking Reagent (Vector laboratories, Cat# MKB-2213-1, diluted 1:20 in 1X PBS) for 1 hour at room temperature, followed by additional blocking and permeabilization with a solution of 3% BSA, 5% filtered horse serum in PBS-T (0.05% Triton-X 100 and 0.03% Tween (Sigma) in 1X PBS) for 1 hour at room temperature. Tissues were then incubated with primary antibodies diluted in blocking solution at 4°C overnight. The following primary antibodies were used for this study: pcm1, Santa Cruz sc398365, 1:50 dilution; pcm1, Sigma HPA023370, 1:500 dilution; Lamin A/C (E-1), Santa Cruz sc376248, 1:100 dilution; lamin A, Sigma MAB3540, 1:100 dilution; CD68 (E3O7V), Cell Signaling 97778T, 1:400 dilution; BAF, Santa Cruz sc-166324, 1:250 dilution; Rtn4, Novus NB100-56681SS, 1:200 dilution; IL-6, Novus NB600-1131, 1:100; CD45 (D3F8Q), Cell Signaling 70257S, 1:100 dilution. Samples were washed with PBS-T and incubated for 1 hour at room temperature with AlexaFluor secondary antibodies (Invitrogen, 1:300 dilution) and Hoechst Nucleic Acid Stains (ThermoFisher 62249, 1:1000) in PBS-T.

### Proximity Image analysis

Images were processed using a custom CellProfiler^24^ script to identify regions of interest (ROIs) corresponding to DAPI-positive nuclei, Rtn4-positive cells, and CD68-positive cells. DAPI-positive nuclei were identified using an Otsu two-class threshold to raw DAPI-channel images, followed by smoothing and segmentation based on size and shape. Rtn4-positive cells were detected using a three-class Otsu threshold on raw Rtn4-channel images, smoothed, and identified based on size. CD68-positive cells were identified by applying manually set threshold bounds, consistent across all images, to raw CD68-channel images, followed by size-based identification. The distance between CD68-positive ROIs and the nearest Rtn4-positive ROI was calculated using the MeasureObjectNeighbors module. All image data was exported from CellProfiler for post-processing using the ExportToSpreadsheet module.

### Western Blot analysis

Cardiac tissue was placed in homogenization strips (VWR, Cat #TN0946-08RS) in high salt RIPA buffer [12 mM sodium deoxycholate, 50 mM Tris-HCl pH 8.0, 750 mM NaCl, 1% (v/v) NP-40 alternative and 0.1% (v/v) SDS in ultrapure water] and a small metal ball (McMaster-Carr, Cat #1598K23). Tissues were homogenized for three 3-minute cycles of 30 shakes/sec using a TissueLyser II (Qiagen). Homogenized samples were sonicated (Branson 450 Digital Sonifier) for 30 s at 36% amplitude, boiled for 2 min, centrifuged at 4°C for 10 min at 14,000 g and stored at −75°C. Protein concentrations were determined by BCA assay, and 10 µg of total protein from each heart was denatured in 5X Laemmli buffer by boiling for 3 min. Protein was loaded onto 4–12% Bis-Tris gels (Invitrogen NP0322), separated for 1.5 h at 100 V, then transferred for 1 h at 16 V onto PVDF membrane. Coomassie staining of the gel following transfer was used to assess equal protein loading. Membranes were blocked for 1 h in blocking buffer containing 3% BSA in Tris-buffered saline plus 1% Tween 20. Primary antibodies were added individually overnight at 4°C diluted in blocking buffer. Primary antibodies used: rabbit anti-CD68 (Cell Signaling 97778; 1:1000) and rabbit anti-Rtn4 (Novus Biologicals 100-56681; 1:1000). Membranes were washed 3 × 10 min, and secondary antibodies were added for 1 h at room temperature in blocking buffer, followed by 3 × 10 min washed. Secondary antibodies used: Licor IRDye 680RD donkey anti-mouse-IgG (LICORbio 926-68072; 1:5000) and Licor IRDye 800CW Donkey anti-Rabbit IgG (LICORbio 926-32213; 1:5000). Membranes were imaged using Odyssey Licor scanner and then cropped and brightness and contrast was adjusted using Image Studio Lite (version 5.2) software.

### Bulk RNA sequencing (RNA-seq) processing

Left ventricle free wall tissues (excluding the atria and septum) were harvested at the appropriate timepoints, flash frozen in liquid nitrogen, and stored at –70°C until RNA was extracted. Frozen tissue was placed in an Eppendorf tube containing one metal bead and 500 µl of TRIzol® reagent (Invitrogen, #15596018). Tissues were homogenized for five 3-minute cycles of 30 shakes/sec using a TissueLyser II (Qiagen) and transferred to an RNase free microfuge tube (Ambion, Cat #AM12400). 1:5 chloroform was added and mixed by shaking, followed by a 5-minute centrifugation at 4°C 14,000 rpm. The clear supernatant was carefully transferred to a gDNA eliminator spin column (RNeasy Plus Mini Kit, Qiagen, #74136) and RNA purification was performed according to manufacturer instruction. RNA quality control was performed by the Cornell Genomics core facility with a fragment analyzer (Agilent) to assess RNA integrity (RIN); RNA concentrations were quantified by Nanodrop (ThermoFisher Scientific). To generate cDNA directional RNA-seq libraries (New England Biolabs, #E7760), 1 µg of total RNA sample (with RIN ≥ 7) was enriched for mRNA (New England Biolabs, #E7490) for each condition. Each sample received a unique barcode for downstream multiplexing. cDNA library size distribution and concentrations were again assessed by fragment analyzer and Qubit. Final libraries were pooled in equimolar concentrations and sequenced. The 24 cKO and cWT samples at both time points (GSE269451), used to generate data for Figure 2, were pooled 12 per lane and sequenced on a NextSeq 500 (Illumina) with a 75 bp single-end kit by the Cornell Genomics core facility. Libraries generated for the 48 samples (GSE304346) associated with the LINC complex disruption experiment (Figure 6) were pooled into a single lane and sequenced on a NovaSeq 6000 (Illumina) with an S4 flow cell and a 150 bp paired-end kit by Novogene.

### Bulk RNA-seq analysis

We used STAR software^25^ (version 2.7.10a) to generate a genome for read alignment using the GRCm38/mm10 (Ensembl release 102) mouse FASTA and GTF files. Fastq files for the DN KASH experiment (GSE304346) had TruSeq adaptors trimmed using fastp^95^ (versions 0.20.0). All fastq files were aligned to the STAR genome using default parameter settings, resulting in a uniquely mapped read alignment ≥ 79%. Gene-level quantification was performed using the featureCounts function from Subread^26^ (version 2.0.1), with parameters -s 2 (for directional RNA-seq libraries), -p (for paired end only with GSE304346), and the GENCODE GTF annotation file (vM23), to generate a count matrix using all samples. PCA and distance correlation matrix analyses were performed to identify and remove outliers (GSE269451: samples 3 and 19, GSE304346: samples 16 and 43) before downstream processing. Differential gene expression (DEG) analysis was performed in R (v4.1.1) by applying DESeq2^27^ (v1.34.0) to the input count matrix. The design formula “∼genotype*timepoint”, where genotypes were either cKO, cWT (for GSE269451) or cKO – DN KASH, cWT – DN KASH, cKO + DN KASH, cWT + DN KASH (for GSE304346), and timepoints were either 11- or 25-days post-injection (dpi), was used. The ashr^96^ shrinkage estimator was applied to all DEG results. DEGs were defined as genes with |log_2_-fold change| ≥ 1, baseMean ≥ 10, and *p*-adj ≤ 0.05.

### Computation of emerging differentially expressed genes (eDEGs)

eDEGs were identified based on the pipeline described below for both bulk RNA sequencing datasets (GSE269451, Suppl. Figure 2A; GSE304346, Suppl. Figure 20A). We first filtered DEG analysis results for cKO versus cWT samples at 11- and 25-dpi, based on meeting a significance threshold of *p*-adj < 0.05 and a base mean expression ≥ 10. We retained all 11-dpi genes that met the threshold (GSE269451: 1,405 genes, GSE304346: 2,668 genes) for downstream analysis, irrespective of fold-change (FC). We further refined the 25-dpi significance-filtered genes using a |log2 FC| ≥ 1 threshold (GSE269451: 2,951 DEGs, GSE304346: 3,426 DEGs), thus meeting DEG requirements outlined in the “Bulk RNA-seq analysis” section. We compared the 11-dpi genes and 25-dpi DEGs, and identified genes with altered expression at 11-dpi that were also found at the 25-dpi time-point (GSE269451: 508 genes, GSE304346: 995 genes). ∼17% of these genes (GSE269451: 85 DEGs, GSE304346: 175 DEGs) had a |log2 FC| ≥ 1, thus representing 11-dpi DEGs. Of the genes that overlapped with 25-dpi DEGs, 496 genes (GSE269451, 98%) and 83 genes (GSE304346, 8%) had the same directional change at both early and late timepoints (i.e., either both up- or down-regulated) and were termed emerging differentially expressed genes (eDEGs). The remaining genes did not have the same directional expression change at both early and late timepoints (described in Suppl. Fig. S2C-D for GSE269451 and S20C for GSE304346).

### Classification of DEG rescue by LINC complex disruption

To assess the extent to which LINC complex disruption rescues gene misregulation in cKO hearts, we analyzed DEGs identified across three pairwise comparisons: (1) cKO hearts without LINC complex disruption versus cWT hearts without LINC complex disruption (cKO – DN KASH vs. cWT – DN KASH) to identify genes misregulated in the diseased state, (2) cKO hearts without LINC complex disruption versus cKO hearts with LINC complex disruption (*Lna* cKO – DN KASH vs. cKO + DN KASH) to evaluate the impact of LINC complex disruption, and (3) cKO hearts with LINC complex disruption versus cWT hearts without LINC complex disruption (cKO + DN KASH vs. cWT – DN KASH) to determine whether gene expression returned to wild-type levels with LINC complex disruptionTabl.

Genes were classified as rescued if they were significantly misregulated in the disease (comparison 1) and significantly altered following LINC complex disruption (comparison 2). Complete rescue was defined as genes whose expression in cKO + DN KASH hearts was no longer significantly different from cWT (not significant in comparison 3). Partial rescue was defined as genes that remained significantly different from cWT despite changes following LINC complex disruption (significant in comparison 3).

### Single-nucleus RNA sequencing and analysis

Single-nucleus RNA-sequencing (snRNA-seq) was chosen over single-cell RNA-sequencing for this study due to several key advantages that are particularly relevant in cardiac research^97^: (1) snRNA-seq minimizes bias in cell type capture related to cell size, which is particularly important for larges cells such as cardiomyocytes; (2) snRNA-seq excludes cytoplasmic RNA, reducing interference from high mitochondrial RNA content commonly found in cardiomyocytes; and (3) snRNA-seq is compatible with frozen tissue samples, eliminating the need for fresh specimens.

For processing, left ventricle free wall tissues (5-6 left ventricles per experimental group) were harvested from an independent subset of cKO and cWT mice at the appropriate timepoints, flash frozen in liquid nitrogen, and sent to SingulOmics (Bronx, NY, USA) for processing. There, nuclei were isolated from frozen samples and used to construct 3’ single nuclei gene expression libraries (Next GEM v3.1) using the 10X Genomics Chromium system. Libraries were sequenced with ≈200 million PE150 reads per sample on Illumina NovaSeq 6000. Sequencing reads, including introns, were then aligned with the mouse reference genome (mm10/GENCODE vM23/Ensembl 98) using Cell Ranger^28^ (v7.0.1). The Debris Identification using Expectation Maximization (DIEM) algorithm^29^ was applied to remove background, keeping barcodes with debris scores < 0.5, UMI count >800, and number of features (genes) > 250. The resulting dataset was composed of 22,043 total nuclei (5,052 11-dpi and 5,939 25-dpi in cKO; 6,300 11-dpi and 4,752 25-dpi cWT), before downstream filtering. snRNA-seq sample outputs from Cell Ranger were then loaded into R^98^ (v4.1.1) and individually subjected to the standard Seurat^30^ (v5.0.1) pipeline and sctransform. All samples were processed for ambient RNA (SoupX^99^, v1.6.2) and doublet (DoubletFinder^100^, v2.0.3) removal individually. Sample integration was performed with the integrated object subjected to the same standard Seurat pipeline as above, with a cluster resolution of 0.5. To annotate individual clusters with cell types using the established Azimuth reference map^31–33^, mouse genes were first converted to human orthologous genes as only human heart references are available. The converted snRNA-seq dataset was then compared to the Azimuth reference map to assess prediction mapping scores^101^. Cells with high-confidence annotations (prediction scores > 0.75) were used to determine cell types in each identified cluster. To identify cell type-specific subpopulations, each cell type cluster was subjected to the standard Seurat (v5.0.1) pipeline and sctransform with unique cluster resolution (0.9 for cardiomyocyte, 0.7 for myeloid cells, 0.5 for fibroblasts, 0.6 for B cells, and 0.8 for NK/T cells). DEG analyses were performed for each cell type cluster using Seurat’s FindMarkers function: 1) cKO vs. cWT at 11- and 25-dpi across all cells in the cluster irrespective of subpopulation, and 2) each subpopulation vs. all other subpopulations irrespective of experimental condition. Genes were required to meet a threshold of |log2 FC| ≥ 0.1, p-adj < 0.05, and for the fraction of cells expressing the gene to be ≥ 0.01 for at least one of the two queried populations. Gene modules scores were generated using Seurat’s AddModuleScore function and gene lists curated by the gene ontology consortium^54,102^ (GO: 0006954, 0031347, and 0008630), Harmonizome^103,104^ (v3.0) (myofibroblast gene set from GeneRIF Biological Term Annotations), and peer-reviewed papers^74^ (infiltrating and resident macrophage markers).

### Cell-cell interaction prediction

Cell-cell communication prediction was determined using CellChat^34^ (version 2.1.1). The CellChat algorithm computed the strength of specific predicted cell-cell interactions, quantified as the sum of the communication probability of all ligand-receptor interactions, in each experimental condition. CellChat objects were generated from Seurat snRNA-seq objects for each condition individually and processed using the default pipeline to identify enriched ligand-receptor interactions between all cell types and subpopulations. CellChat objects were then merged based on timepoints to perform a comparative analysis of cell-cell communication between cKO and cWT samples over time.

### Pseudotime analysis

Pseudotime trajectories were inferred using Slingshot^35^ (v2.2.1) applied to the PCA embedding of the integrated cardiomyocyte subpopulations. CM1 was designated as the root population and CM3 as the terminal population. These assignments were based on their temporal distribution: CM1 was consistently present across all conditions at 11-dpi, while CM3 emerged only at 25-dpi.

### Functional enrichment analysis

DEGs and eDEGs Log2-fold change-ranked gene symbol lists were submitted to Metascape^36^ (v3.5.20240101) and QIAGEN’s Ingenuity Pathway Analysis (IPA) software^37^ (v111725566) for functional enrichment analysis. All analyses used the gene ontology^54,102^ (GO) biological processes database for enrichment. To perform comparative analyses between gene sets, multiple gene lists were provided simultaneously to Metascape using default parameters (i.e. p-value < 0.01, enrichment ≥ 1.5, overlap ≥ 3). Metascape calculates enrichment using a hypergeometric test. Comparative analyses of predicted upstream regulators between different gene sets were performed using IPA, assuming a |z-score| > 2 and *p*-value < 0.05 as predictive of regulator activation or inhibition.

### Human DCM snRNA-seq dataset

Publicly available snRNA-seq data of genetic dilated cardiomyopathies^38^ were obtained from the CZ CELLxGENE data portal^39^ in h5ad format. We extracted samples annotated as “heart left ventricle” and classified as either “dilated cardiomyopathy” or “normal” using Scanpy^40^ (v1.9.3) in Python (v3.7.12). The resulting AnnData object was exported by separating it into counts.mtx, obs.csv, and var.csv files. These files were subsequently imported into R to create a Seurat object, which was subjected to the standard Seurat (v5.0.1) pipeline and sctransform with cluster resolution of 0.4.

### Tissue collection for Curio Seeker spatial transcriptomics

Isolated hearts were perfused and then coated in Tissue-Tek O.C.T. Compound (Sakura Finetek # 4583), inside cryomolds to be snap-frozen in liquid nitrogen cooled isopentane (Thermo Scientific Chemicals # AA19387AP) and stored at –80°C. Frozen hearts were sectioned with a cryostat at 10 µm thick slices at –20°C.

### Spatial transcriptomic library preparation and sequencing for cardiac samples

Spatial transcriptomic libraries were prepared from heart tissue sections obtained from cKO and cWT mice at 11- and 25-days post injection (dpi). Samples were mounted directly onto barcoded Curio Seeker arrays (Curio Biosciences), corresponding to tile IDs B0040_010 (cKO, 11-dpi), B0040_012 (cWT, 11-dpi), B0040_011 (cKO, 25-dpi), and B0040_014 (cWT, 25-dpi). The protocol followed Curio’s spatial transcriptomics workflow, beginning with RNA hybridization and in situ reverse transcription on the array. Following enzymatic tissue digestion, spatially barcoded beads were released into solution, and second-strand cDNA synthesis and amplification were performed. Sequencing libraries were constructed using the Nextera XT DNA kit, pooled, and sequenced on an Illumina NovaSeq X Plus platform (Read 1/2: 150 bp, Index 1/2: 8 bp). Raw sequencing data were processed using the slide_snake pipeline (https://github.com/mckellardw/slide_snake). Adapter trimming and quality filtering were performed using the seeker_v3.1_internalTrim configuration (cutadapt v4.6)^105^, and reads were aligned to the mouse genome (GRCm39) using STARsolo (v2.7.10b) to generate bead-resolved gene expression matrices^41^.

### Curio Seeker data preprocessing and quality control

For each sample, AnnData objects (v0.10.3) were generated from the raw count matrices, and spatial coordinates were assigned to beads using metadata provided by Curio Biosciences (also known as slide-seq^106^). Preprocessing was performed using Scanpy^40^ (v1.9.3), which included filtering out beads with fewer than 200 detected transcripts, normalizing expression values to a fixed total count per bead, and log transformation. Beads with fewer than 10 neighbors within a 100 μm radius were also excluded, as these often represent smeared or low-quality barcodes. Data were plotted, and regions of interest corresponding to the left ventricles were segmented using Napari^107^. To further reduce noise while preserving spatial structure, expression data were smoothed using the Smoothie package implementation with a Gaussian kernel of 30 μm^108^.

### Cell type deconvolution

We applied cell2location^42^ (v0.1.3) to deconvolve the collected spatial transcriptomics data using the single-nucleus dataset discussed in this work as a reference. Cell-subpopulation-specific expression signatures were estimated from the reference and projected onto the spatial data to infer the contribution of each cell subpopulation category in the spatial beads. Cell2location was run either on full heart sections or focused specifically on the left ventricle annotated regions. Model hyperparameters were set to N_cells_per_location = 1 and detection_alpha = 20.

### Cardiomyocyte neighbor score calculation

To assess the local enrichment of specific cardiomyocyte populations, we implemented a k-nearest neighbors (kNN) approach. For each bead, we computed a Cardiomyocyte neighbor score by counting how many of its

10 nearest spatial neighbors shared the same predicted cardiomyocyte subpopulation (CM1 to CM5). This analysis was performed independently for each of the segmented ventricular spatial transcriptomics objects.

### Neighborhood enrichment analysis on spatial transcriptomics data

Z-score calculations of neighborhood enrichment was calculated using Squidpy^43^ (v1.2.2) for all experimental conditions. A heatmap of this enrichment was generated with Seaborn^109^ (v0.12.2). Finally, visualization of individual subpopulations across the tissue was generated through Squidpy.

### Gene-set score calculation on spatial transcriptomics data

Gene set activity scores were then computed in Scanpy with default parameters for curated programs related to regulation of defense, interferon-beta response, intrinsic apoptosis, and inflammatory response. For each left ventricle sample, scores were compared across five cardiomyocyte subpopulations (CM1–CM5). To evaluate whether these distributions differed significantly within each heart, a Kruskal–Wallis H-test was first performed. In cases where the Kruskal–Wallis test returned a *P* value < 0.05, pairwise Mann–Whitney U tests were conducted to identify specific group differences. All pairwise p-values were adjusted for multiple testing using the Benjamini–Hochberg false discovery rate (FDR) correction, applied independently within each heart sample. All tests were two-sided and implemented using the scipy.stats and statsmodels.stats Python libraries.

### Data availability

The transcriptomic datasets generated from cardiac tissue samples and used to support the conclusions in this article are limited to reviewers and journal editors for the preprint version of the manuscript. All data will be made publicly available in the GEO repository following peer review and publication.

### Statistical analysis

Animals were randomly assigned to treatment groups, and experiments were conducted blinded to treatment and genotype. Since Cre expressed in the heart can be cardiotoxic at high tamoxifen doses, experiments used only low tamoxifen doses (30 mg/kg/day). Furthermore, for maximum stringency, comparisons are focused on groups in which all mice express the MCM transgene but differ in the status of the floxed *Lmna* gene. Tissues isolated from individual mice were counted as a single replicate (unless otherwise specified). Since the bulk-RNA sequencing data did not show any sex difference, male and female mice were combined for downstream transcriptomic analysis. Survival curves were generated using the Kaplan–Meier method and differences in survival tested with a Breslow test using GraphPad Prism 10. For results that follow a Gaussian distribution, the mean and standard error of the mean (SEM) were calculated; comparison between two groups used *t*-tests; ANOVA with Tukey or Dunnett post-analysis was used to test for statistical significance between more than two experimental groups. Data linearity was determined using Pearson correlation coefficients and significance calculated using the R package ggpubr^110^. The non-parametric pairwise wilcoxon rank sum test was calculated to determine significant changes in gene module scores between conditions in R. Pairwise statistical comparisons were summarized using compact letter display (CLD). Groups that share at least one letter are not statistically different from each other, while groups with no letters in common differ at *P* < 0.05. Letters are arranged in aligned rows to simplify visual scanning across multiple groups. Finally, the concordance correlation coefficient (CCC) was calculated using the equation:

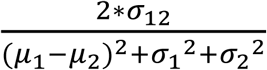

where *σ*_12_ is the covariance, *μ* is the mean, and *σ* is the standard deviation.

## Acknowledgments

The authors thank Stephen Weiss (University of Michigan) and Didier Hodzic (Washington University at St. Louis) for generously providing the floxed *Lmna* mice and DN KASH expressing mice, respectively. In addition, the authors thank Ern Hwei Hannah Fong for help with tamoxifen injections, and Alexander Sopilniak Mints, Hind Zahr, and Julie Heffler for help with weaning and genotyping mice. Bulk RNA sequencing was performed by the Biotechnology Resource Center (BRC) Genomics Facility (RRID:SCR_021727) at the Cornell Institute of Biotechnology.

