## Supplementary Materials for "Nuclear envelope rupture in cardiomyocytes orchestrates early transcriptomic changes and immune activation in *LMNA*-related dilated cardiomyopathy that are reversed by LINC complex disruption"

### Supplemental Information

The Supplemental Materials in this file contain Supplemental Figures S1-S26.

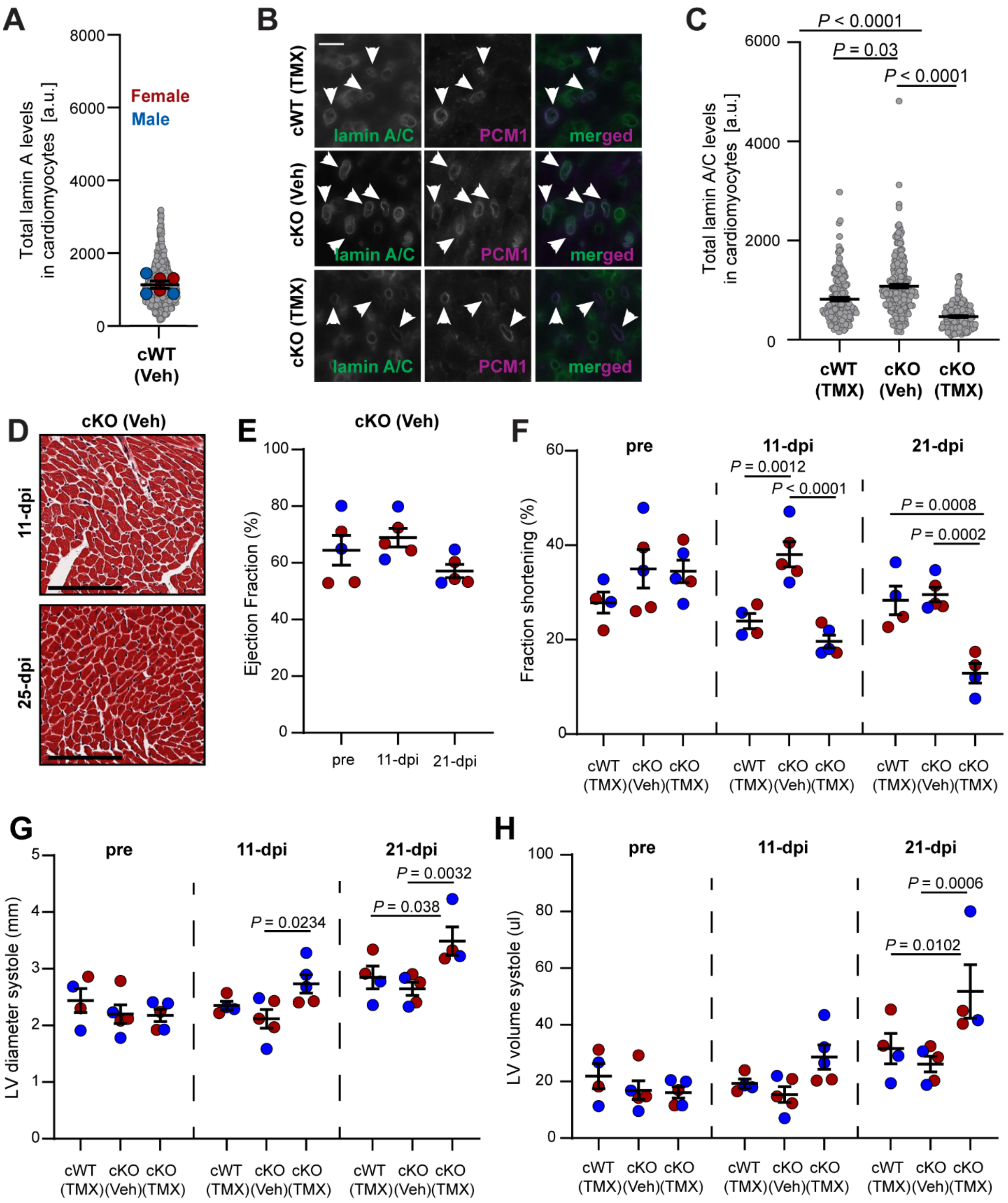


**Supplementary Figure S1: Reduction of lamin A/C in cardiomyocytes in adult mice leads to dilated cardiomyopathy. (A)** Quantification of total lamin A levels in vehicle-only treated cKO cardiomyocytes. Larger colored circles represent average values for individual mice. Smaller gray circles represent individual cardiomyocytes. *N* = 6 mice/condition. **(B)** Representative images of cardiac tissues from cWT mice treated with

30 mg/kg tamoxifen (TMX) (top), cKO mice treated with vehicle (Veh) control (middle), and cKO mice treated with 30 mg/kg tamoxifen (bottom), immunofluorescently labeled for lamin A (green) and PCM1, a cardiomyocyte-specific nuclear marker (magenta). Scale bar: 20 µm. Arrows indicate representative cardiomyocytes. **(C)** Quantification of total lamin A/C levels in cardiomyocytes from vehicle-only-treated cKO and tamoxifen-treated cKO and cWT hearts, confirming the depletion of lamin A/C in cardiomyocytes of tamoxifen-treated cKO mice. Gray circles represent individual cardiomyocytes (1 mouse per condition). **(D)** Masson's trichrome staining of heart sections to detect fibrosis in vehicle-only-treated cKO hearts at both early (11-dpi) and late (25-dpi). Scale bar: 100 µm. Echocardiography-based measurements of **(E)** left ventricular ejection fraction (EF), given as percentage, for vehicle-only treated cKO mice, and **(F)** fractional shortening (FS), given as percentage, **(G)** left ventricle (LV) diameter during systole, measured in mm, and **(H)** left ventricular volume during systole, measured in mm for vehicle-only treated cKO mice, and tamoxifen-treated cKO and cWT mice . Symbols represent the mean values of individual mice. Error bars represent mean ± S.E.M. **(B-C, E-H)** Statistical analysis by Tukey Honest Significant Difference of one-way ANOVA test. **(E-H)** Color indicates sex of individual mice (red: female, blue: male). *N* = 4-5 mice/condition.

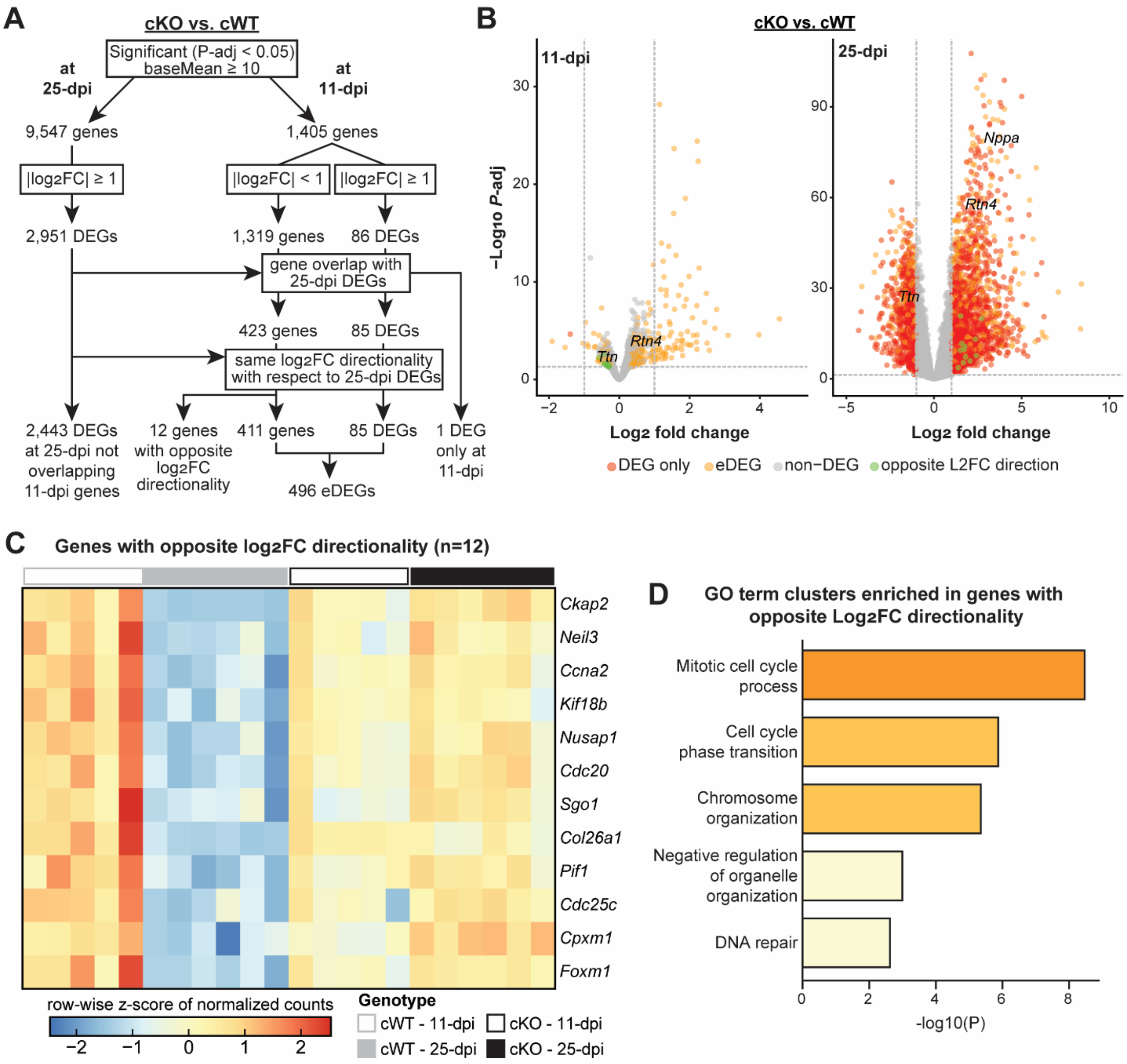


**Supplementary Figure S2: eDEG pipeline and analysis in bulk RNA-seq data of left ventricular tissue. (A)** Flow chart demonstrating the steps to identify emerging differentially expressed genes (eDEGs) from bulk RNA-seq data, based on $\log_2$-fold change ($\mathrm{Log_2FC}$) and adjusted *P*-values (*P*-adj) generated when comparing the gene expression between cKO and cWT samples at each timepoint. **(B)** Volcano plot of the cKO versus cWT DEG analysis at 11- (left) and 25-days post injection (dpi) (right). Genes meeting DEG requirements (*P*-adj < 0.05 and $|\mathrm{Log_2FC} \geq 1|$) but not defined as eDEG are represented in red, eDEGs are shown in orange, the 12 genes with opposite L2FC directionality between 11- and 25-dpi are shown in green, and genes not meeting either DEG or eDEG requirements are shown in gray. Select genes identified throughout the study are highlighted. **(C)** Heatmap of row-wise *z*-score normalized read counts for the 12 genes which had opposite $\mathrm{Log_2FC}$ directionality (cKO versus cWT) at 11- and 25-dpi. Columns, indicating individual samples, were sorted by experimental conditions denoted below by color. Gene names are highlighted. **(D)** Bar plot of gene ontology (GO) term clusters enriched in the 12 genes with opposite $Log_2FC$ directionality between the 11- and 25-dpi timepoints.

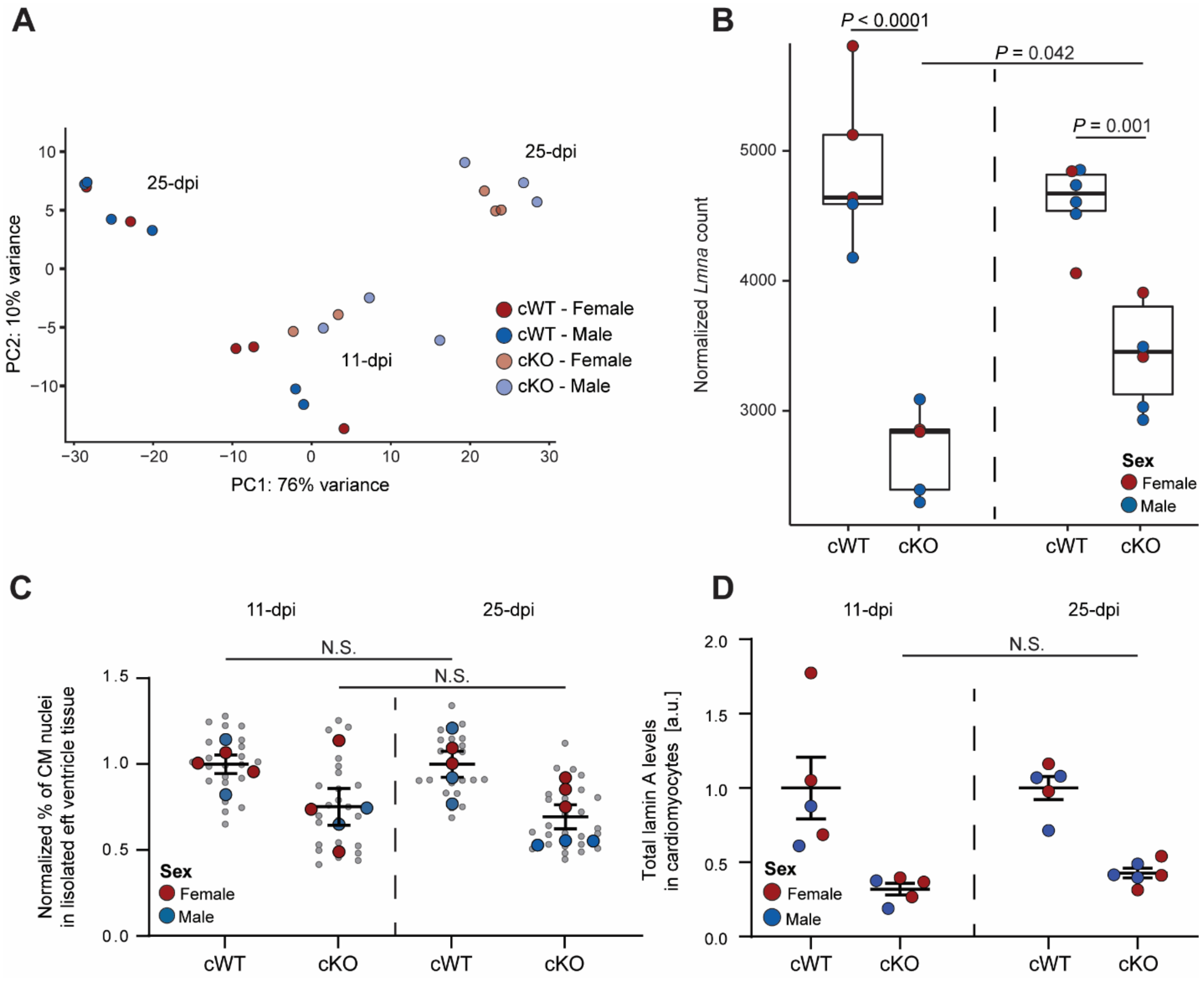


**Supplementary Figure S3: Transcriptomic and immunofluorescence studies of Lmna-cKO and Lmna-cWT hearts did not show significant sex differences. (A)** Principal components analysis (PCA) plot of bulk RNA-seq sample from all experimental conditions. *N* = 5-6 mice/condition. **(B)** Quantification of normalized *Lmna* read counts from bulk RNA-seq samples of cKO and cWT at 11- (left) and 25-dpi (right) . Median (thick line) and first (bottom hinge) and third (top hinge) quantiles are represented. *N* = 5-6 mice/condition. **(C)** Quantification of the normalized percentage of cardiomyocytes (CM) in 11-days post injection (dpi) (left) and 25-dpi (right) left ventricles. Gray dots represent individual fields of view images (10/mouse), large dots represent the average values for each individual mouse analyzed. *N* = 5 mice/group. **(D)** Quantification of lamin A levels in cardiomyocytes at 11-dpi (left) and 25-dpi (right). *N* = 5 mice/group. **(B-D)** Statistical analysis by Tukey Honest Significant Difference of one-way ANOVA test, N.S. = not significant. **(A-D)** Color indicates sex of individual mice (red: female, blue: male).

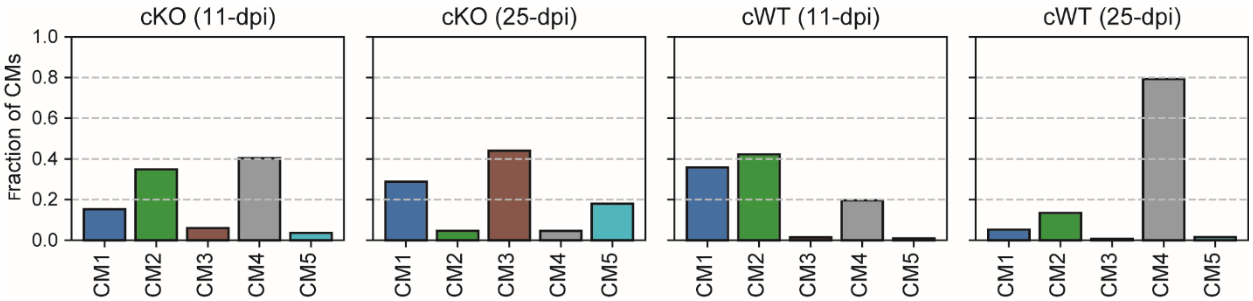


**Supplementary Figure S4: Spatial transcriptomics analysis confirms the presence of disease-specific cardiomyocyte subpopulations.** Fraction of Curio Seeker (Slide-seq) spots, 10 µm spatially barcoded locations on a slide array, assigned to each cardiomyocyte state (CM1–CM5) in the left ventricle of cKO and control cWT hearts at 11-dpi and 25-dpi. Fractions were computed within each sample by normalizing the number of spots assigned to each CM state to the total number of cardiomyocyte-labeled spots in that ventricle.

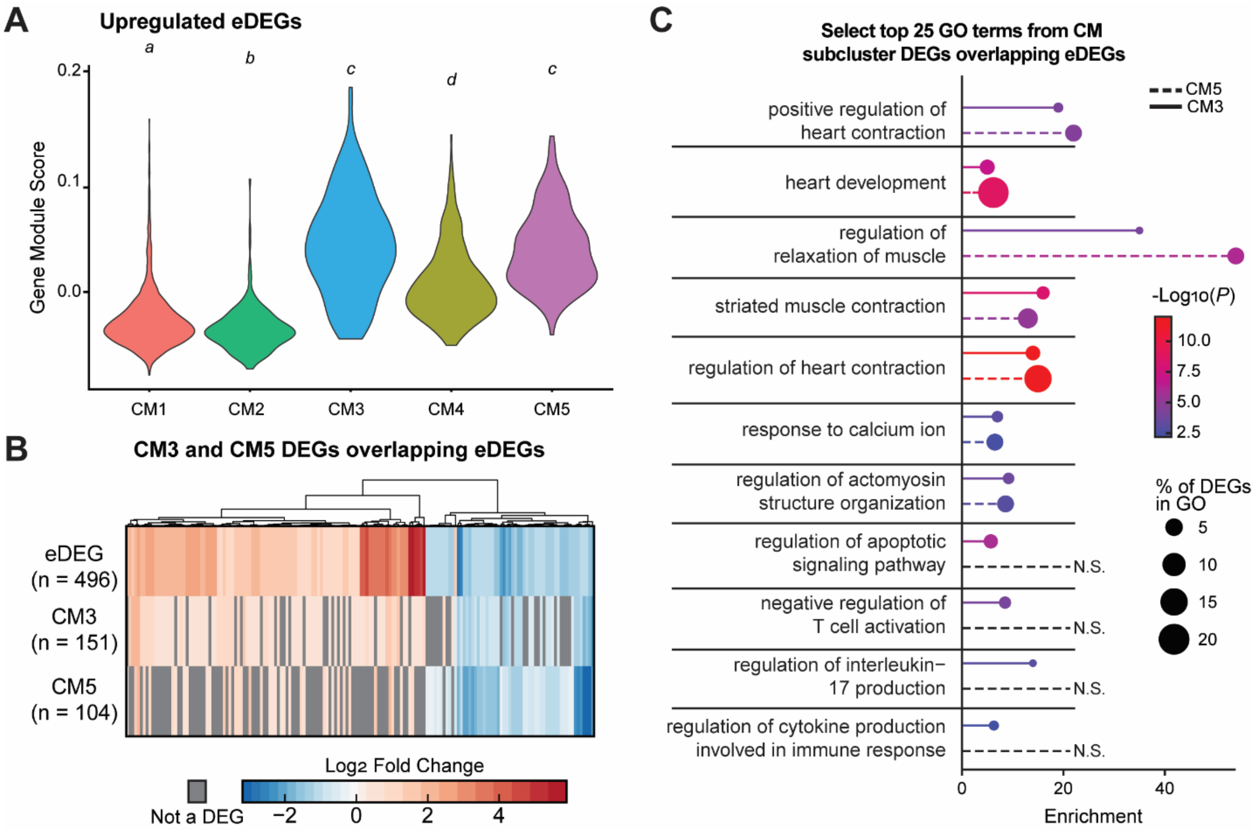


**Supplementary Figure S5: Disease-specific cardiomyocyte subpopulations are enriched for eDEGs. (A)** Violin plot of gene module scores for gene list consisting of emerging differentially expressed genes (eDEGs) upregulated between cKO and cWT at both timepoints across individual cardiomyocyte subpopulations (CM1-5) in the integrated snRNA-seq ventricle dataset. Groups that share a letter are not significantly different; different letters indicate $P$ < 0.05, pair-wise Wilcoxon rank sum test with Benjamini-Hochberg *p*-value adjustment. **(B)** Heatmap of log2 fold change for bulk RNA-seq eDEGs that overlap cardiomyocyte subcluster CM3 and CM5 subcluster-specific DEGs. Gray boxes indicate genes that were not subcluster-specific DEGs for CM3 or CM5. **(C)** Lollipop plot of select top 25 Gene Ontology (GO) terms enriched in disease-specific cardiomyocyte (CM) subpopulations, CM3 (continuous line) and CM5 (dashed line), differentially expressed genes (DEGs) that overlap eDEGs. Bar indicates enrichment score, while dot color shows significance of enrichment, and dot size represents the percentage of subpopulation-specific DEGs contained within the term. Non-significant (N.S.) terms are highlighted.

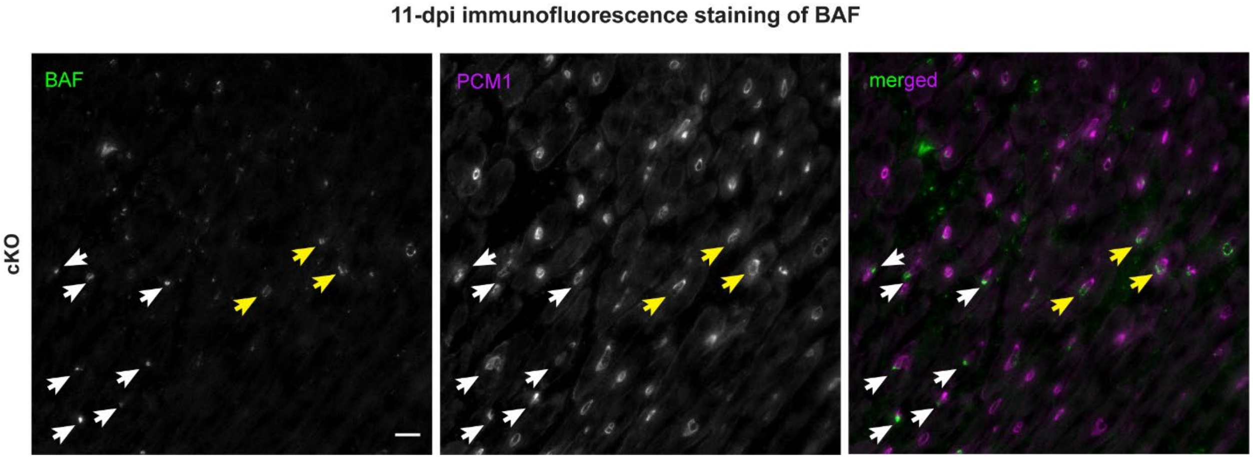


**Supplementary Figure S6: *Lmna*-deficient cardiomyocytes have increased rates of nuclear envelope rupture.** Representative images of cardiac tissue sections from cKO mice at 11 days post injection with antibodies against the nuclear envelope rupture marker BAF (green) and the cardiomyocyte specific marker PCM1 (magenta). Scale bar 20 µm. Arrows indicate representative cardiomyocytes positive for nuclear envelope rupture. Yellow arrows point to the nuclei depicted in the main text figure 3A.

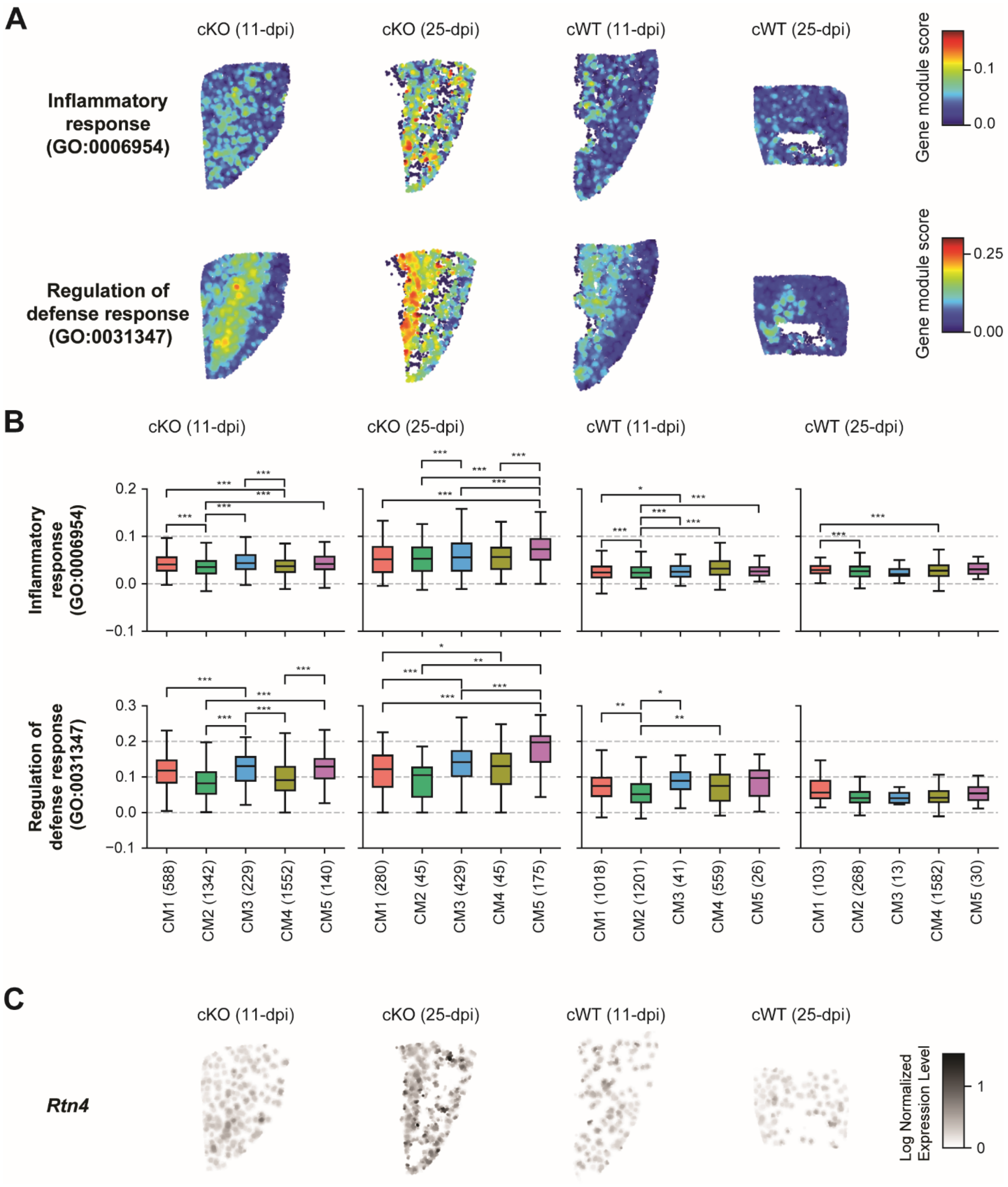


**Supplementary Figure S7: Spatial transcriptomics analysis reveals enrichment of inflammatory and cytosolic DNA sensing pathways in cardiomyocyte subpopulations specific to cKO hearts. (A)** Heatmap and **(B)** boxplot of gene module scores for GO terms “Inflammatory response” and “regulation of defense response” across **(A)** the left ventricle and **(B)** CM1-5 of 11- and 25-days post injection (dpi) of cKO and cWT heart tissue slices based on spatial transcriptomics. Module scores were computed on smoothed, normalized gene expression data. **(B)** Sample size (number of spots per CM group) is indicated below each label. Statistical differences between CM states within each heart were assessed using Kruskal–Wallis tests followed by pairwise Mann–Whitney U tests if the global test was significant ($P < 0.05$). Benjamini–Hochberg FDR correction was applied independently within each heart sample, and adjusted *p*-values are denoted as follows: *, $P < 0.05$; **, $P < 0.01$; ***, $P < 0.001$. **(C)** Heatmap of $\log_2$ normalized *Rtn4* expression across the left ventricle of 11-dpi (left) and 25-dpi (right) cWT and cKO heart tissue slices based on spatial transcriptomics.

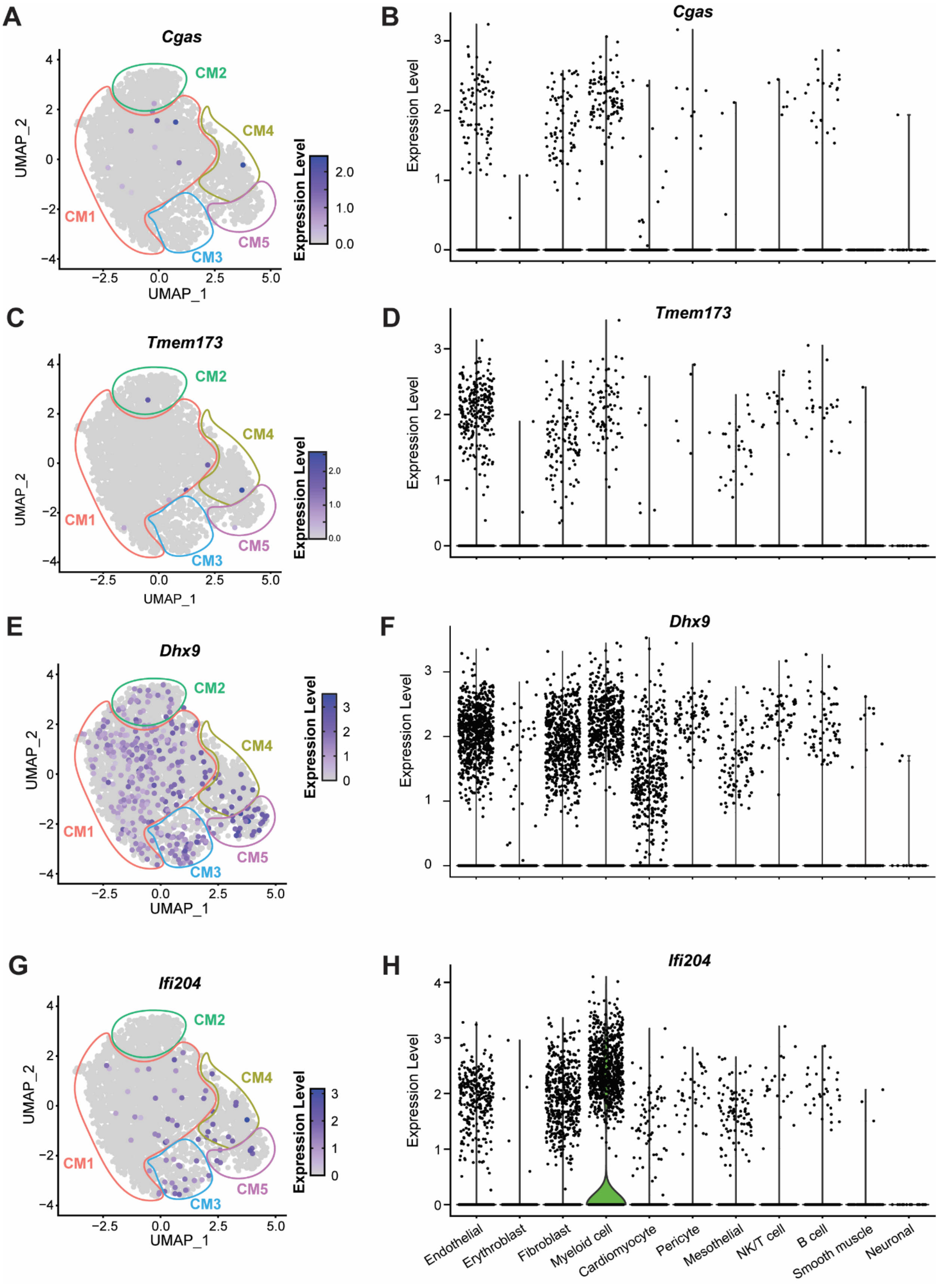
A
Cgas
CM1
CM2
CM3
CM4
CM5
UMAP_1
UMAP_2
Expression Level
B
Cgas
Expression Level
C
Tmem173
D
Tmem173
E
Dhx9
F
Dhx9
G
Ifi204
H
Ifi204
Endothelial
Erythroblast
Fibroblast
Myeloid cell
Cardiomyocyte
Pericyte
Mesothelial
NK/T cell
B cell
Smooth muscle
Neuronal

**Supplementary Figure S8: Cardiomyocytes lack expression of *Cgas* and *Sting1*.** Normalized expression levels for **(A-B)** *Cgas*, **(C-D)** *Sting1* (*Tmem73*), **(E-F)** *Dhx9*, and **(G-H)** *Ifi204* based on single-nucleus RNA sequencing (snRNA-seq) data **(A,C,E,G)** across the UMAP of all captured cardiomyocyte nuclei and **(B,D,F,H)** across all cell types in violin plots. **(A,C,E,G)** Circles of color in the UMAP plots indicate limits of cardiomyocyte subclusters (CM1-5).

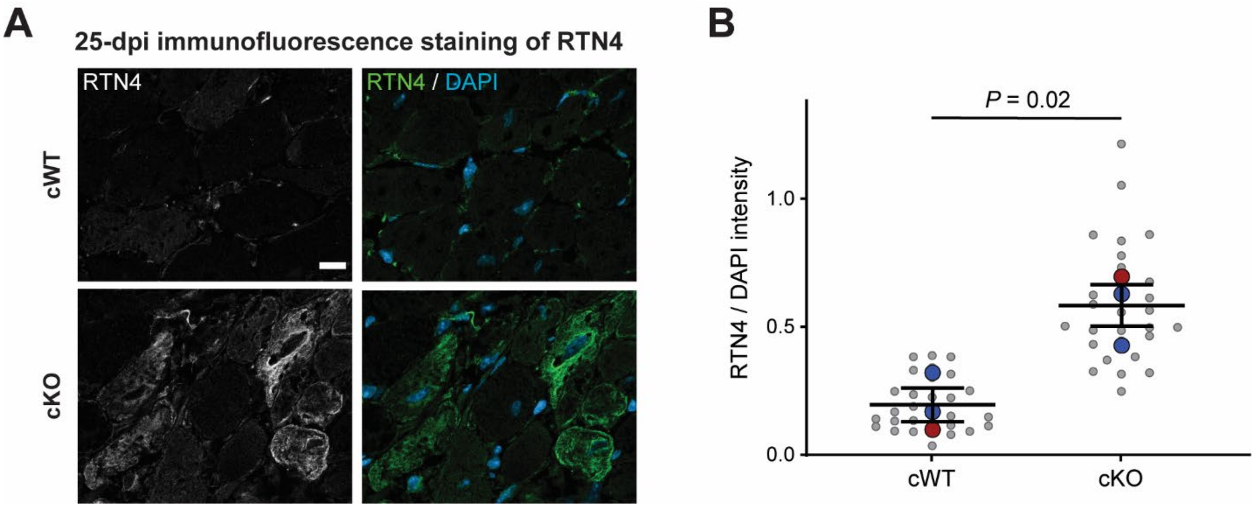


**Supplementary Figure S9: Protein quantifications support transcriptomic findings of increased immune cell activation through cytosolic DNA sensing in cKO hearts.** (**A**) Immunofluorescence labeling for RTN4 (gray on the left, green on the right) and nuclei using DAPI (blue) for cWT (top) and cKO (bottom) hearts at 25-dpi. Scale bar 20 μm. (**B**) quantification of RTN4 over DAPI intensity from representative images. Larger colored circles represent average values for individual male (blue) or female (red) mice. *N* = 3 animals per genotype. Statistical analysis represented performed by *t*-test.

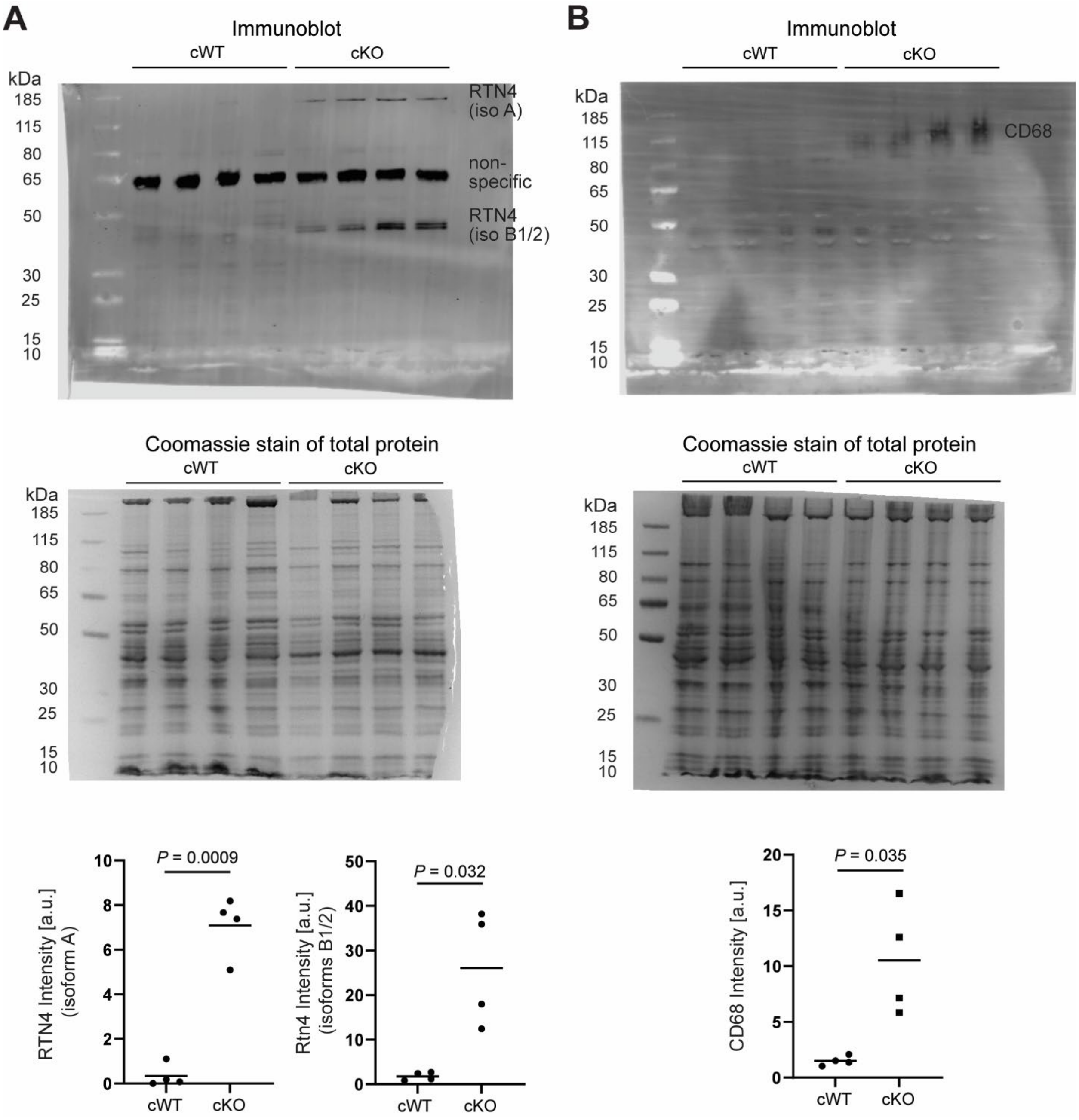


**Supplementary Figure S10: Protein quantifications support transcriptomic findings of increased immune cell activation through cytosolic DNA sensing in cKO hearts.** Immunoblot (top) of heart tissue protein lysates following immunoprecipitation with **(A)** anti-RTN4 and **(B)** anti-CD68 antibodies. Coomassie stains of western gels (middle) were used to ensure even protein loading and for normalization. Quantification (bottom) of band intensity, normalized to coomassie intensity, for replicates showing significant increased protein levels in cKO hearts versus cWT controls. **(A)** Individual isoforms of RTN4 (A and B1/2) are denoted in the gel. *N* = 4 animals per genotype. Statistical analysis by student *t*-test.

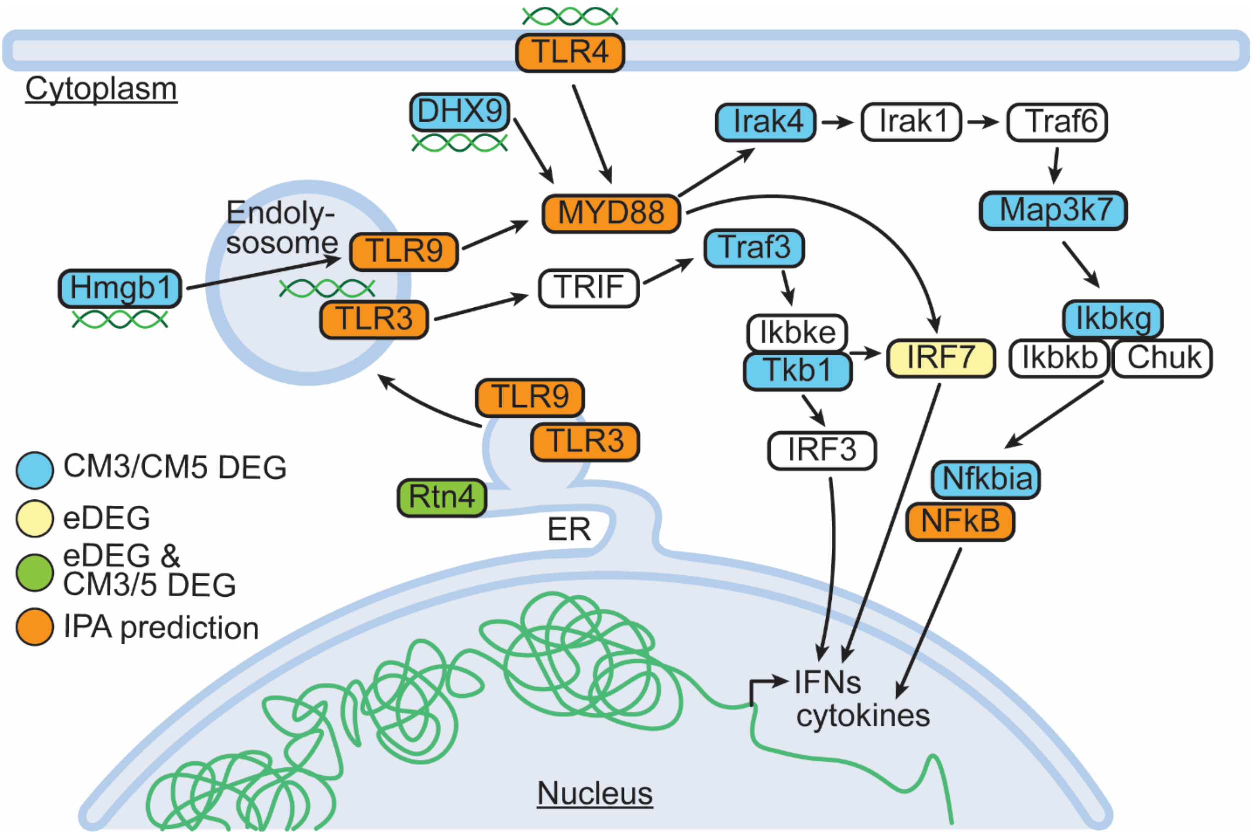


**Supplementary Figure S11: Disease-specific cardiomyocyte subpopulations exhibit evidence for activation of cytosolic DNA sensing independent of cGas/STING pathways.** Diagram of the Tool-like receptor (TLR)-mediated cytoplasmic pattern recognition receptor (PRR) signaling pathway in the cell interior, curated from multiple sources[43–46]. Colors around protein names indicate whether these were detected in the single nucleus RNA-seq cardiomyocyte subpopulations CM3 and CM5 DEGs (blue), appeared as an eDEG from the bulk RNA-seq dataset (yellow), appeared as both an eDEG and CM3/CM5 DEG (green), were predicted as an upstream regulator in Ingenuity Pathway Analysis (IPA) using genes that are both eDEG and CM3/CM5 DEGs (orange), or met none of these criteria (white). Green lines represent nuclear, cytoplasmic, and extracellular DNA.

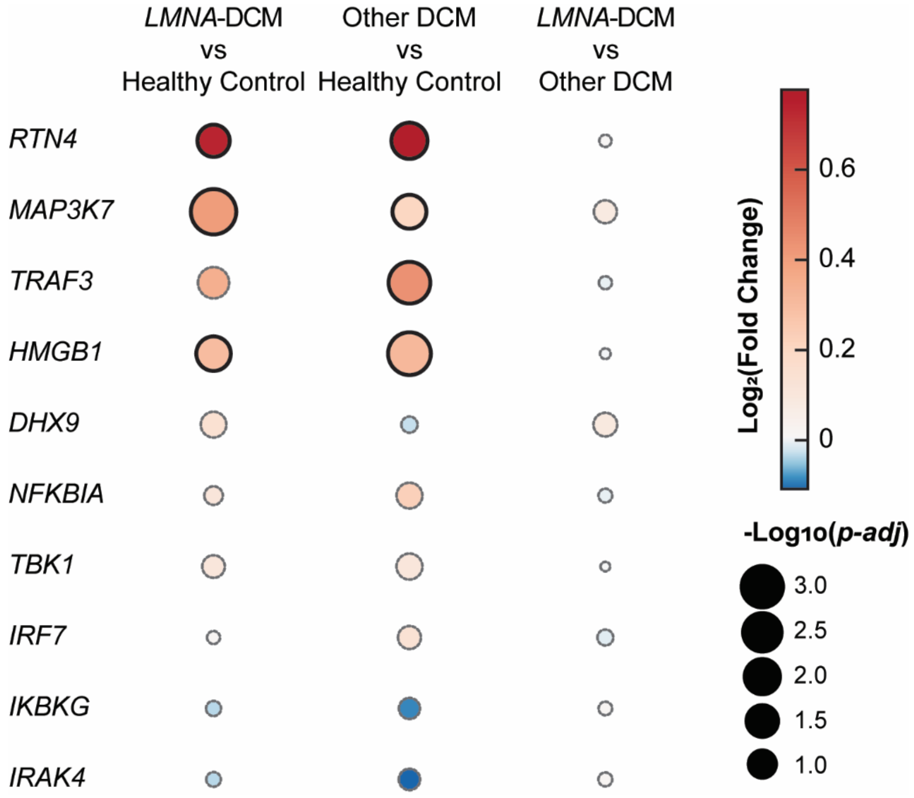


**Supplementary Figure S12: Cytoplasmic pattern recognition receptor (PRR) signaling is upregulated in several forms of human dilated cardiomyopathy (DCM).** Dot plot illustrating differential gene expression from pseudobulk snRNA-seq analysis of human left ventricle cardiomyocytes[29] comparing *LMNA*-DCM, other genetic DCMs, and healthy controls. Shown are cytoplasmic PRR genes identified as either eDEGs or CM3/CM5 DEGs (corresponding to the green, blue, and yellow gene sets in Suppl. Figure S11). Dot color represents effect size ($\log_2$ fold change), while dot size reflects statistical significance ($-\log_{10}$ *p-adj*) for the comparison indicated in each column. Genes reaching significance (i.e. $-\log_{10}$(*p-adj*) $< 0.05$) are outlined with solid black borders.

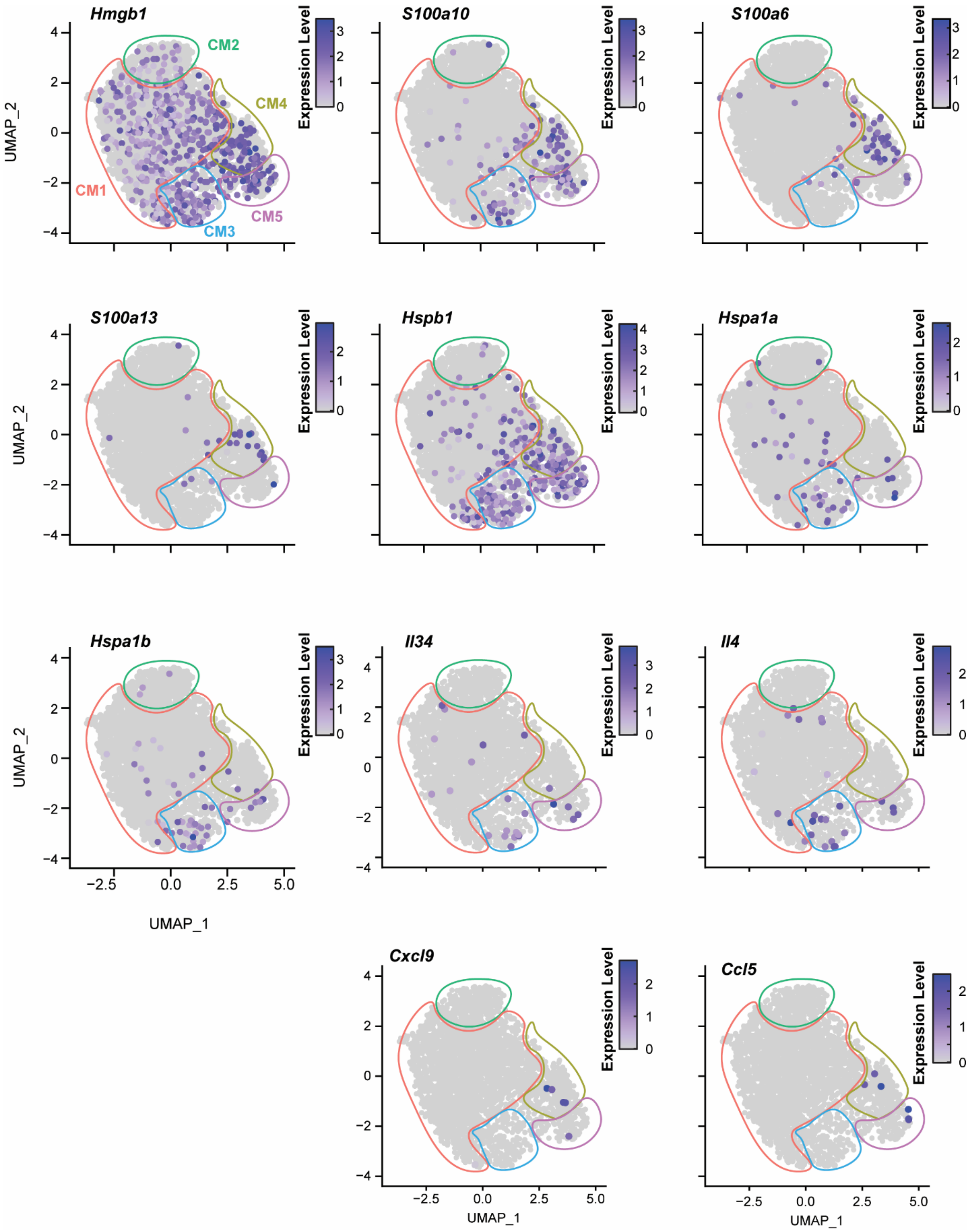

Hmgb1
S100a10
S100a6
S100a13
Hspb1
Hspa1a
Hspa1b
Il34
Il4
Cxcl9
Ccl5
CM1
CM2
CM3
CM4
CM5
UMAP_1
UMAP_2
Expression Level

**Supplementary Figure S13: Disease-specific cardiomyocyte subpopulations have increased expression of DAMPs and cytokines.** Expression of various damage-associated molecular pattern (DAMP) (*Hmgb1, S100a10, S100a6, S100a13, Hspb1, Hspa1a, Hspa1b*) and cytokine (*Il34*, *Il4*, *Cxcl9*, *Ccl5*) genes in snRNA-seq data across the UMAP of all captured cardiomyocyte nuclei. Circles of color indicate limits of cardiomyocyte subclusters (CM1-5).

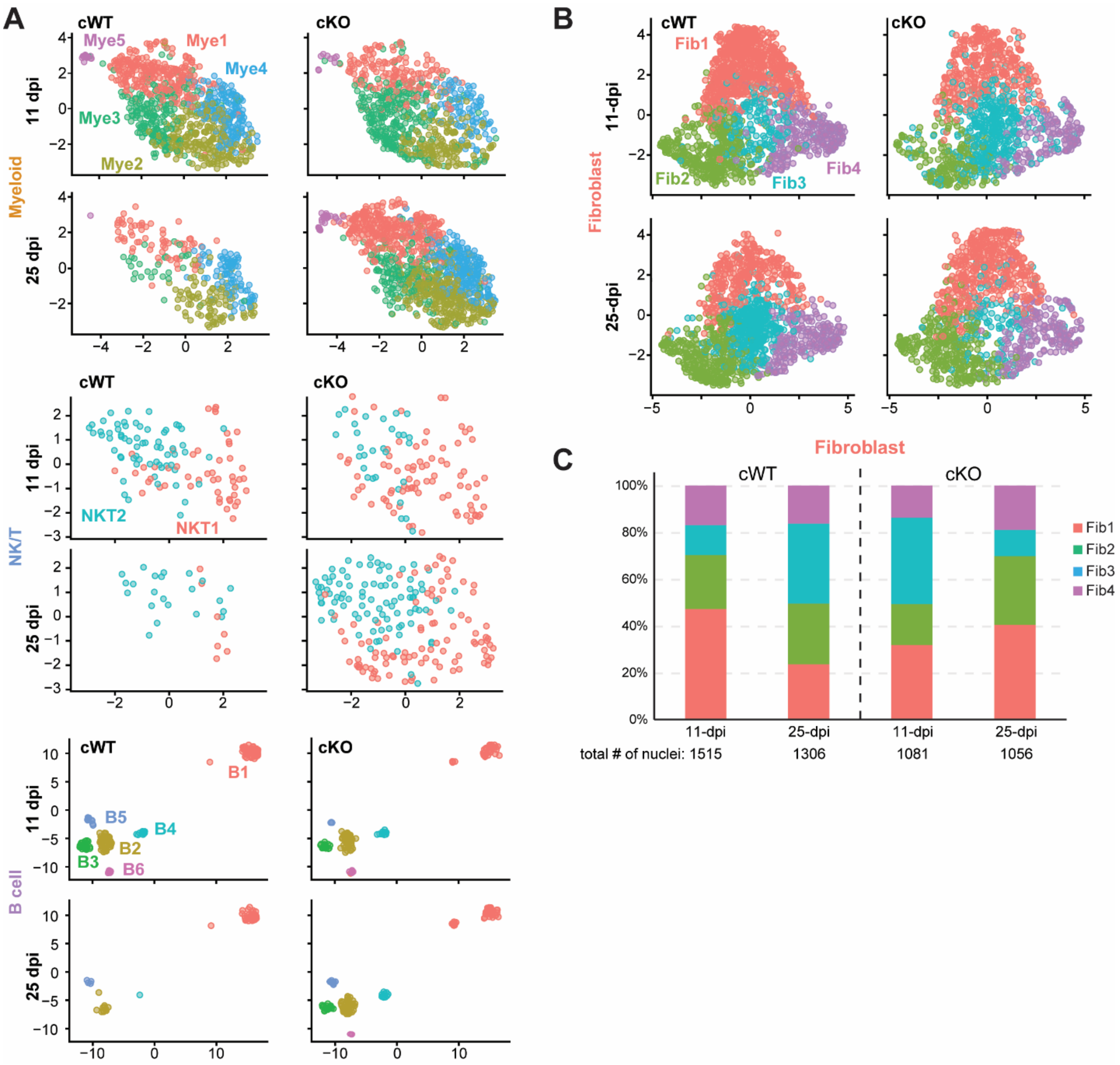


**Supplementary Figure S14: Non-cardiomyocyte cells exhibit distinct subpopulation compositions in cKO hearts. (A,B)** UMAP plots of **(A)** immune cells (Myeloid cells – top, Natural killer T cells – middle, and B cells – bottom) and **(B)** fibroblast-annotated nuclei from integrated snRNA-seq data, across genotype (left and right) and time point (top and bottom). Individual colors represent unique subpopulation identified. **(C)** Stacked bar plot showing the percent composition of each fibroblast (Fib) subcluster for every experimental condition.

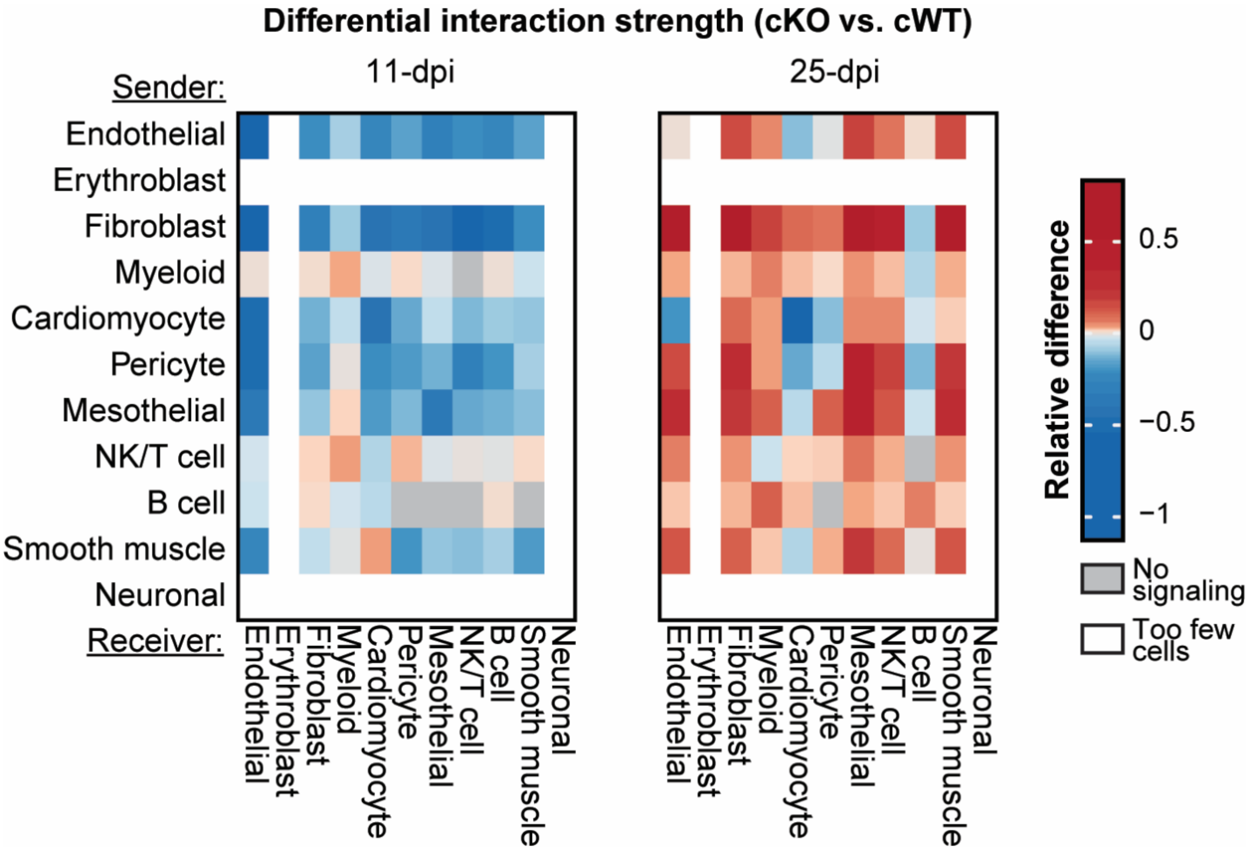


**Supplementary Figure S15: Cell-cell communication across all captured cell types.** Heatmap showing relative difference (cKO versus cWT) in the interaction strength of sender (rows) and receiver (columns) cell types at 11- (left) and 25-dpi (right). Interactions where no communication was identified (gray) and where too few cells were available to perform communication predictions (white) are shown.

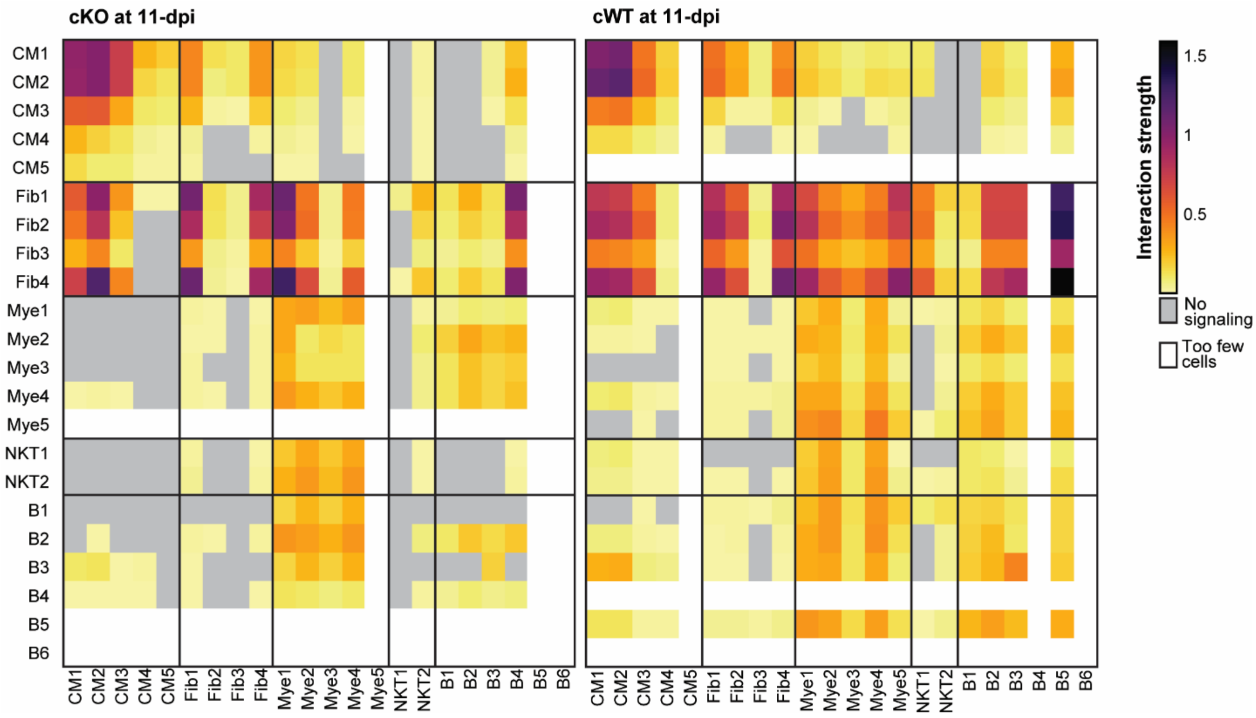


**Supplementary Figure S16: cKO hearts show reduced cell-cell communication in comparison to cWT samples.** Heatmap of the cell-cell communication interaction strength between all subcluster populations of interest at 11-days post injection (dpi) in cKO (left) and cWT (right). Interactions where no communication was identified (gray) and where too few cells were available to perform communication predictions (white) are shown.

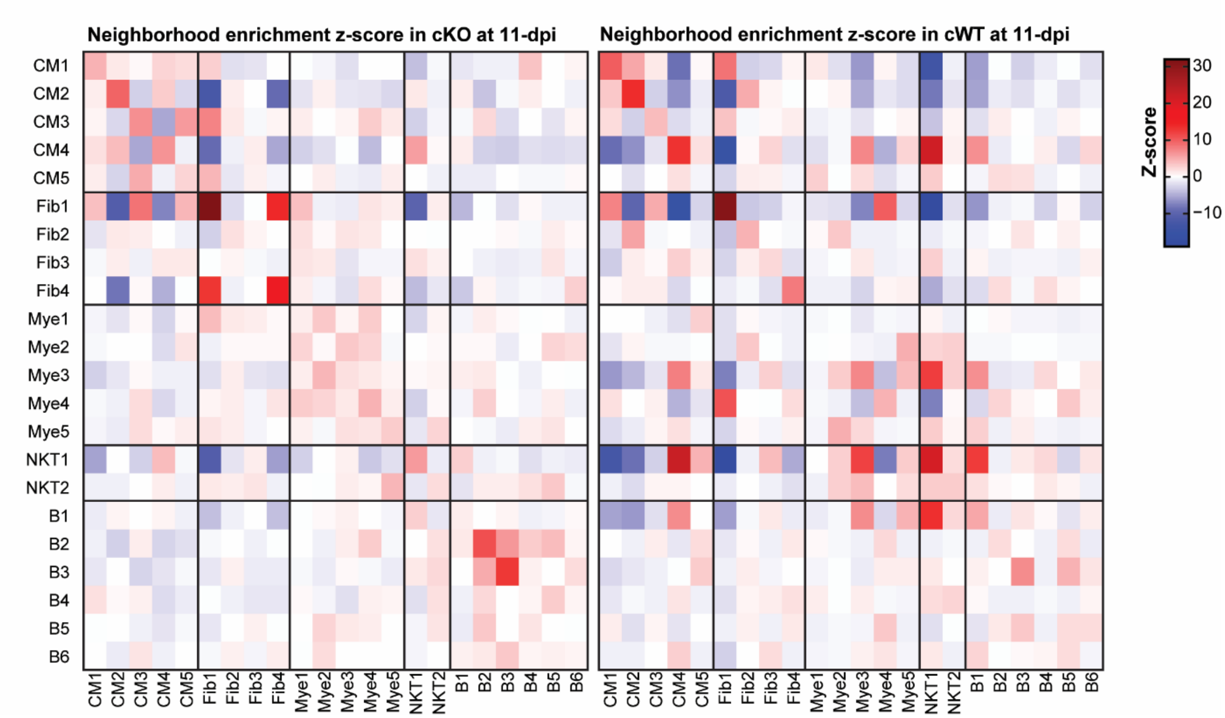


**Supplementary Figure S17: Spatial reorganization of cells in cKO hearts promotes aberrant cell-cell communication. (A)** Heatmap of *z*-score neighborhood enrichment for cell type subpopulations in 11-days post injection (dpi) cKO (left) and cWT (right) ventricles based on spatial transcriptomics.

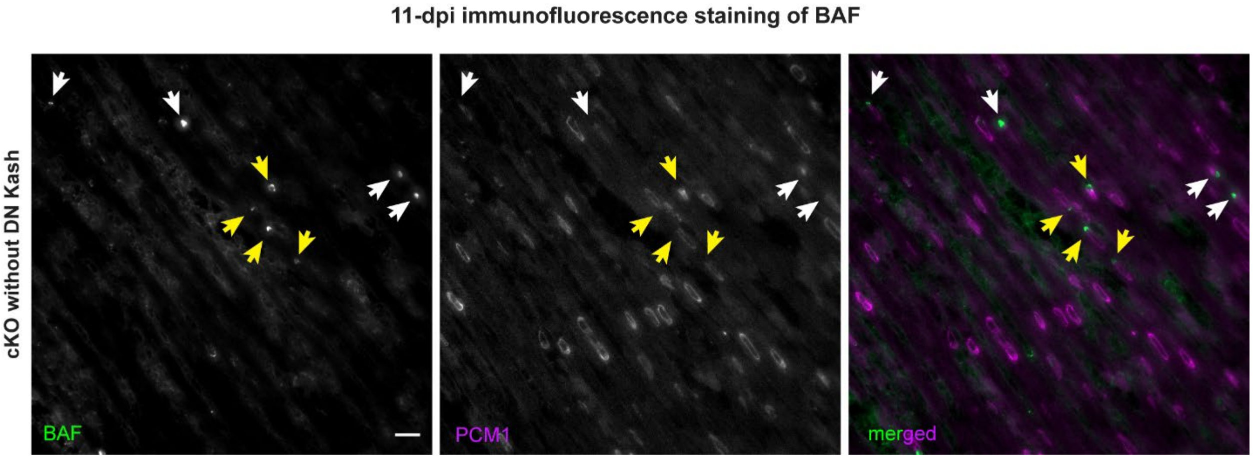


**Supplementary Figure S18: *Lmna*-deficient cardiomyocytes without LINC complex disruption have increased rates of nuclear envelope rupture.** Representative images of cardiac tissue sections from cKO mice without LINC complex disruption at 11-dpi with antibodies against the nuclear envelope rupture marker BAF (green) and the cardiomyocyte specific marker PCM1 (magenta). Scale bar 20 µm. Arrows indicate representative cardiomyocytes positive for nuclear envelope rupture. Yellow arrows point to the nuclei depicted in the main text figure 6B.

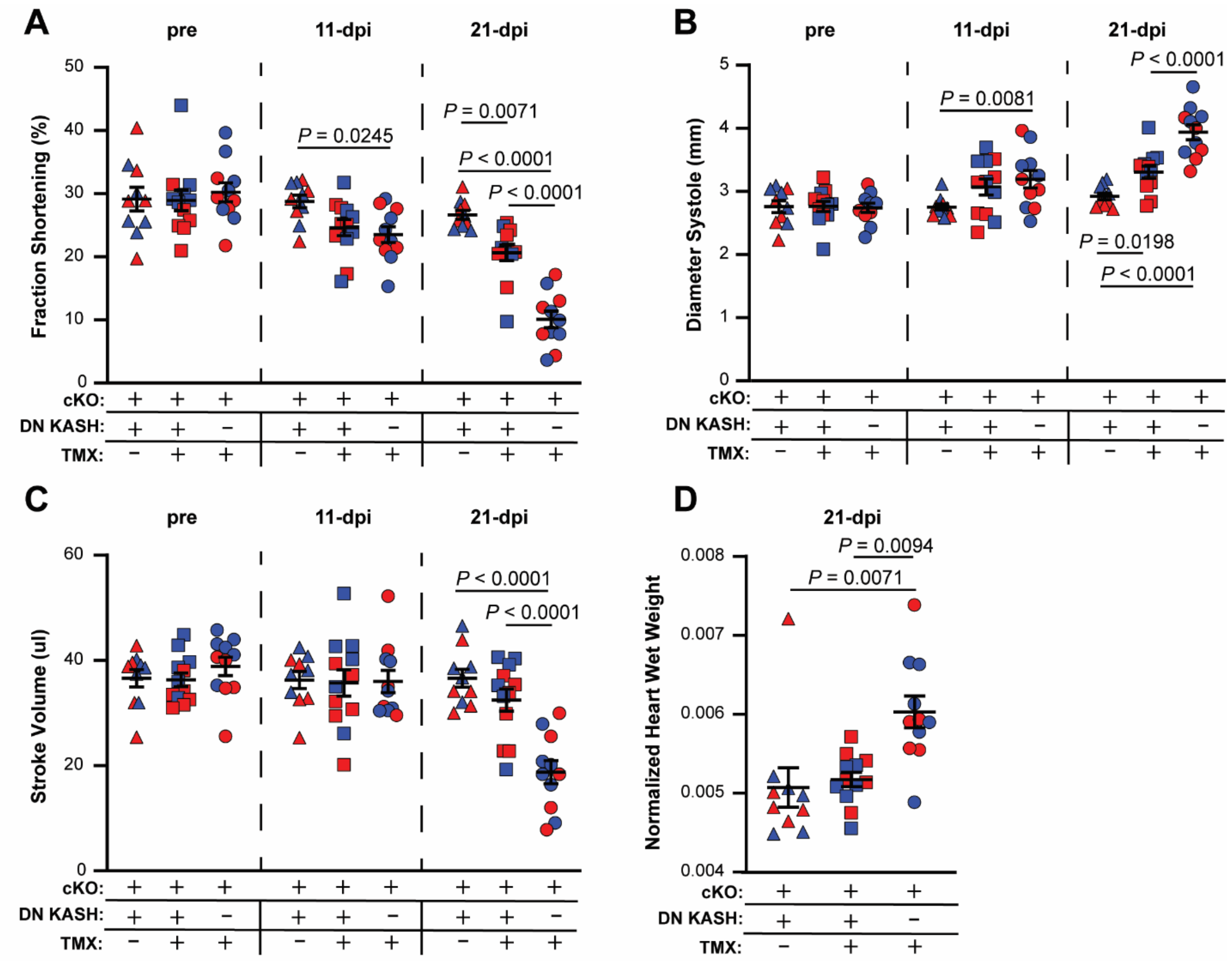


**Supplementary Figure S19: LINC complex disruption rescues cardiac function in cKO mice. (A-D)** Functional cardiac measurements of left ventricles from cKO mice with (+) and without (–) cardiomyocyte-specific LINC complex disruption using a dominant negative nesprin KASH domain construct (DN KASH) induced by tamoxifen (TMX) injection. **(A)** Fractional shortening (given as percentage), **(B)** diameter systole, and **(C)** stroke volume were measured by echocardiography either before TMX injection (pre, triangles), or at 11-days post injection (dpi) (squares) and 21-dpi (circles). **(D)** Heart wet weight normalized to mouse body weight was assessed at 21-dpi. Color indicates sex of individual mice (red: female, blue: male). Error bars represent mean ± s.e.m. *N* = 10-12 animals per genotype. Statistical analysis based on two-way ANOVA test with multiple parameters.

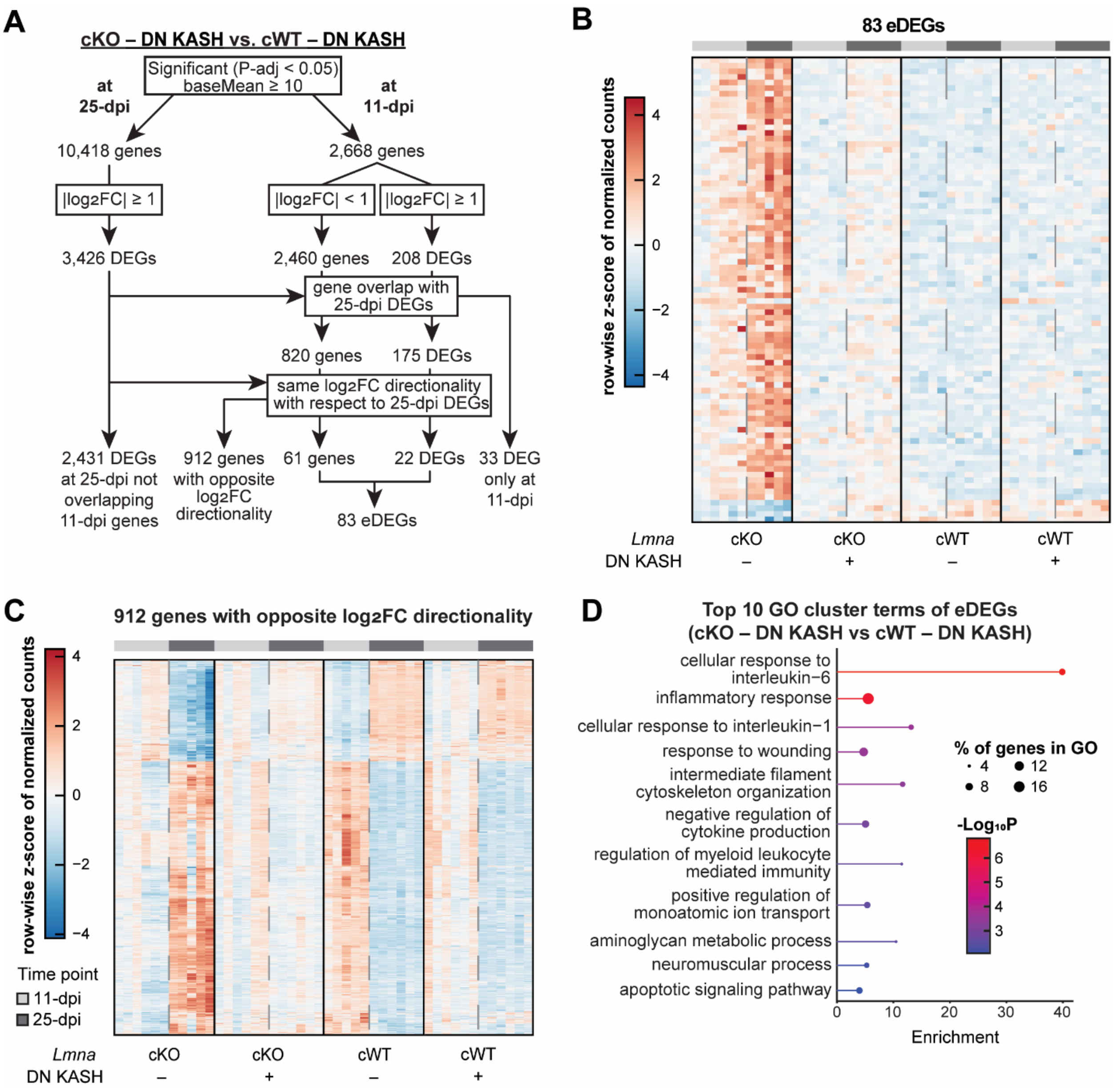


**Supplementary Figure S20: eDEG pipeline and analysis in bulk RNA-seq data of left ventricular tissue for mice without LINC complex disruption. (A)** Flow chart demonstrating the steps to identify emerging differentially expressed genes (eDEGs) from bulk RNA-seq data, based on log2-fold change (Log2FC) and adjusted *P*-values (*P*-adj) generated when comparing the gene expression between cKO and cWT samples both lacking LINC complex disruption (– DN KASH) at each timepoint. **(B)** Heatmap of row-wise *z*-score normalized read counts for the 83 eDEGs (cKO – DN KASH versus cWT – DN KASH) at 11- (light gray) and 25-dpi (dark gray). Columns, indicating individual samples, are separated by condition, where samples with (+) and without (–) cardiomyocyte-specific LINC complex disruption (DN KASH) are specified. **(C)** Heatmap of row-wise *z*-score normalized read counts for the 912 genes that had opposite $log_2$-fold change ($log_2FC$) directionality (cKO – DN KASH versus cWT – DN KASH) at 11- (light gray) and 25-dpi (dark gray). Columns, indicating individual samples, are separated by condition, where samples with (+) and without (–) cardiomyocyte-specific LINC complex disruption (DN KASH) are specified. **(D)** Lollipop plot of the top 10 Gene Ontology (GO) cluster terms enriched in the 83 eDEGs (cKO – DN KASH versus cWT – DN KASH). Color indicates significance of enrichment and dot size represents the percentage of eDEGs contained within the term.

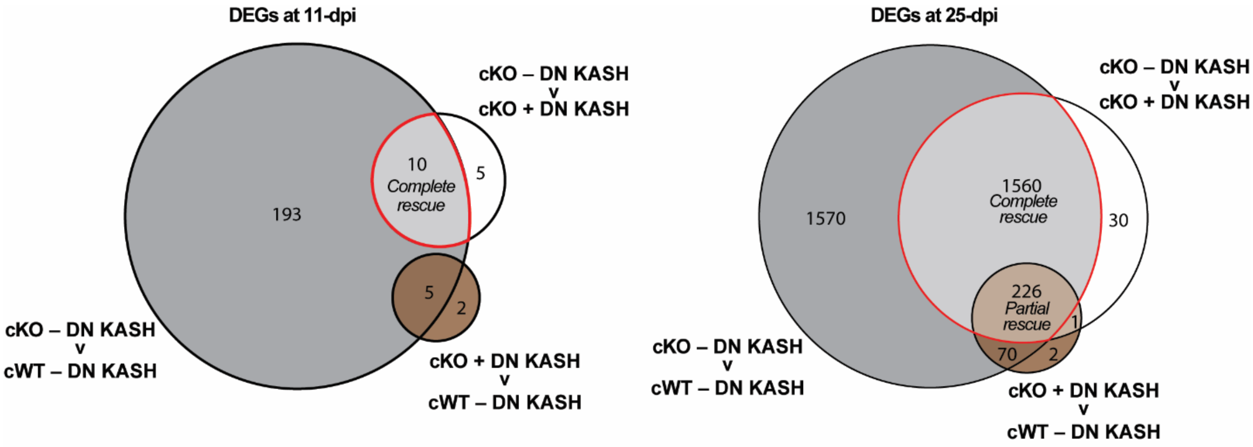


**Supplementary Figure S21: LINC complex disruption partially rescues aberrant cKO gene expression.** Euler plots showing the number of differentially expressed genes (DEGs) generated through three experimental condition comparisons at 11- (right) and 25-dpi (left). Overlaps between the comparisons indicate DEGs rescued by LINC complex disruption (red circle). Distinctions between *Complete* and *Partial* rescue are also denoted.

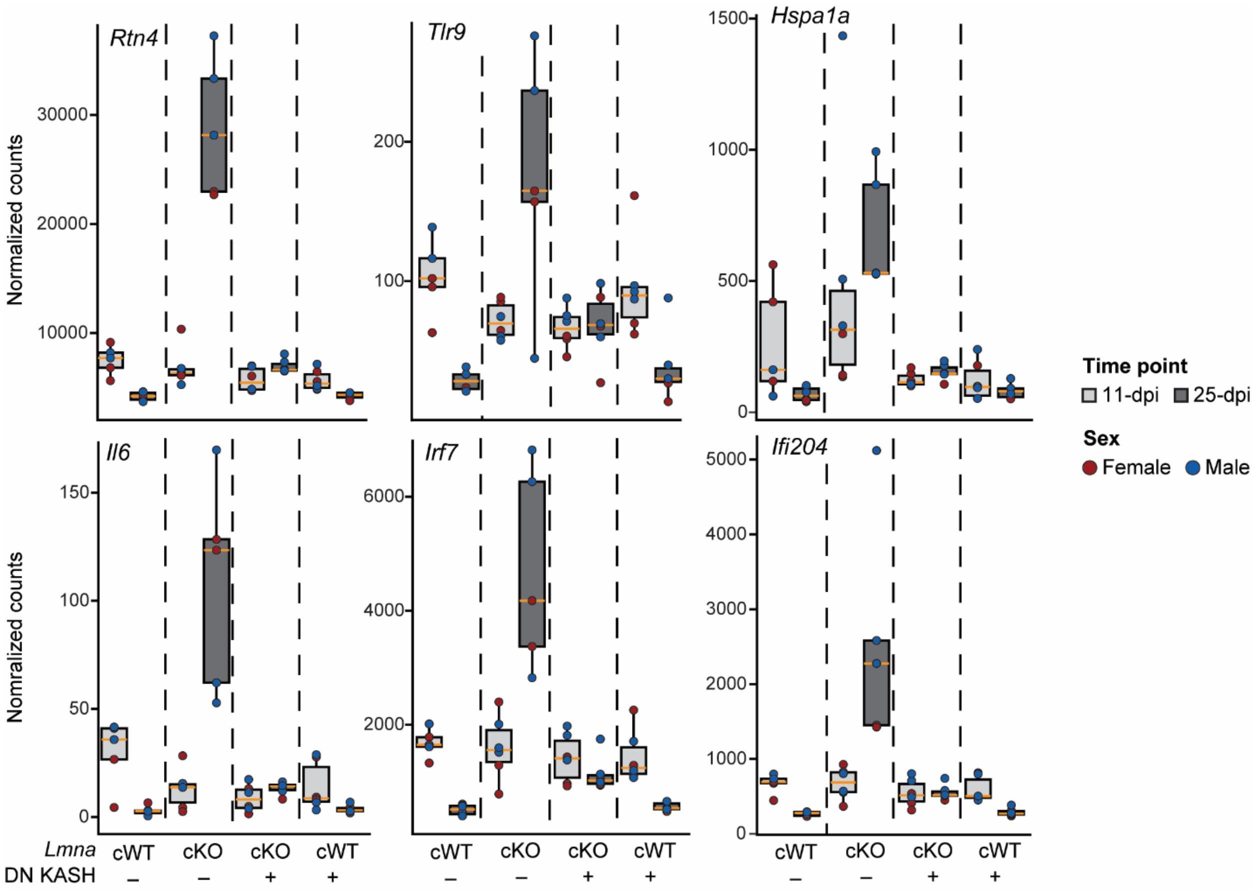


**Supplementary Figure S22: LINC complex disruption restores normal expression of cytosolic DNA sensing genes.** Quantification of normalized read counts for genes of interest involved in cytosolic DNA sensing within bulk RNA-seq samples across each experimental condition. cKO and cWT samples at 11- (light gray) and 25-days post injection (dpi) (dark gray) with (+) and without (–) cardiomyocyte-specific LINC complex disruption (DN KASH) are specified. Median (orange line) and first (bottom hinge) and third (top hinge) quantiles are represented. Color indicates sex of individual mice (red: female, blue: male). *N* = 4-5 animals per genotype.

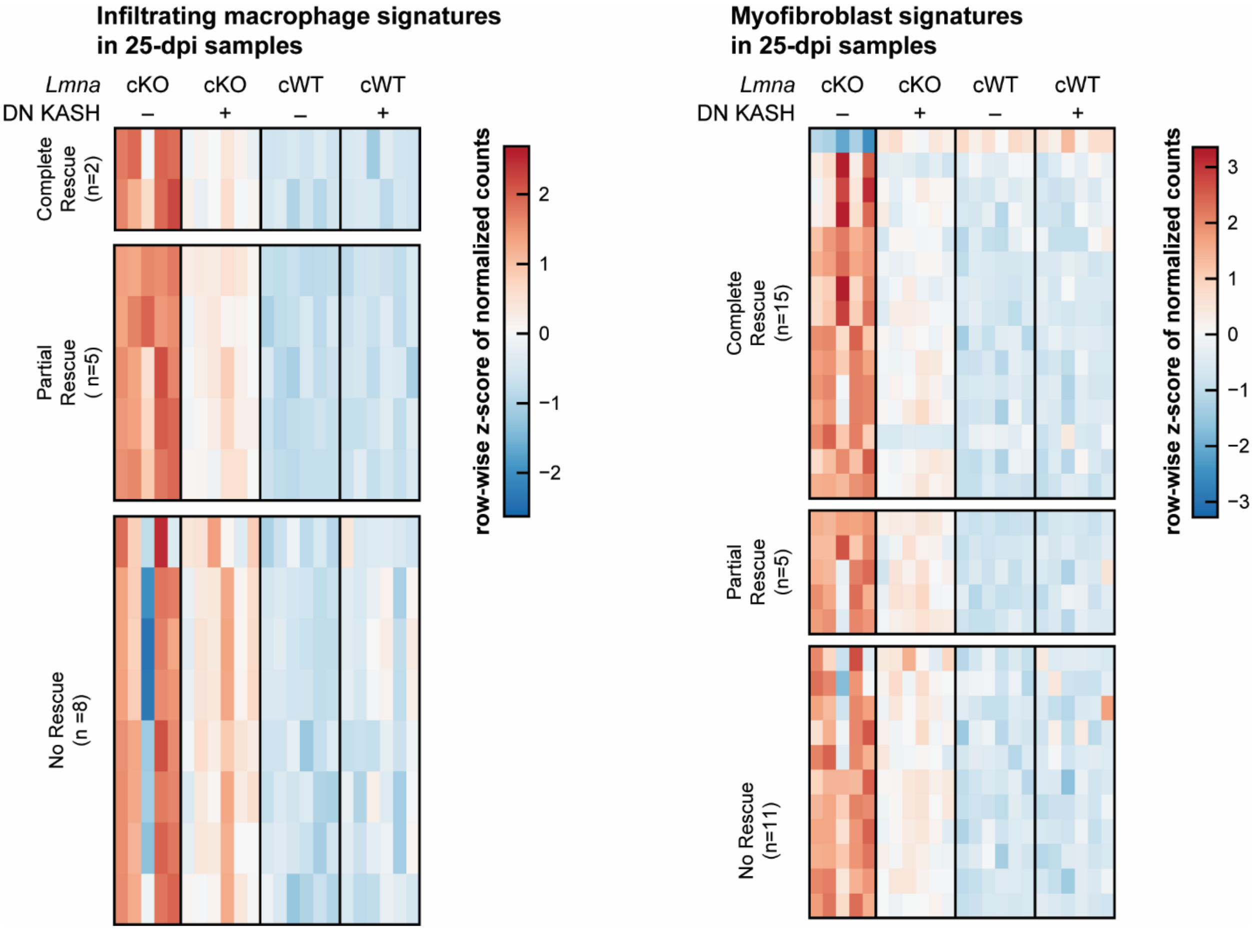


**Supplementary Figure S23: Pathological transcriptional signatures show rescue by LINC complex disruption. (A)** Heatmaps of row-wise z-scored normalized read counts of cKO – DN KASH vs. cWT – DN KASH DEGs representative of infiltrating macrophage (left) and myofibroblast (right) signatures across all experimental conditions. Heatmaps are split based on degree of rescue by LINC complex disruption induced by DN KASH expression.

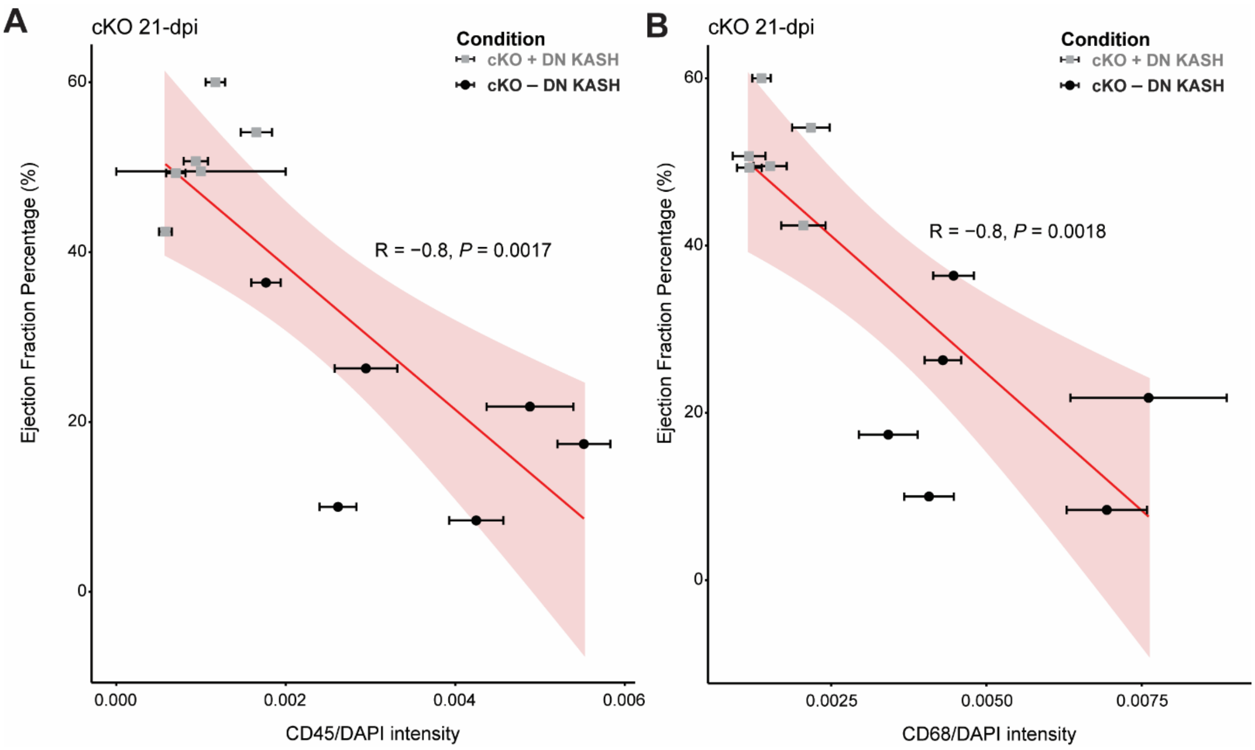


**Supplementary Figure S24: Cardiac function inversely correlates to immune cell activation.** Pearson correlation between the ejection fraction percentage and immunostaining intensity normalized to DAPI stain of left ventricle tissue for either **(A)** CD45 or **(B)** CD68 in cKO mice at 21-dpi. The presence (gray squares, cKO + DN KASH) or lack (black circles, cKO – DN KASH) of LINC complex disruption is indicated. *N* = 6 animals per genotype. Pearson correlation coefficient (R) and p-values for a linear fit (red) are displayed in the graphs.

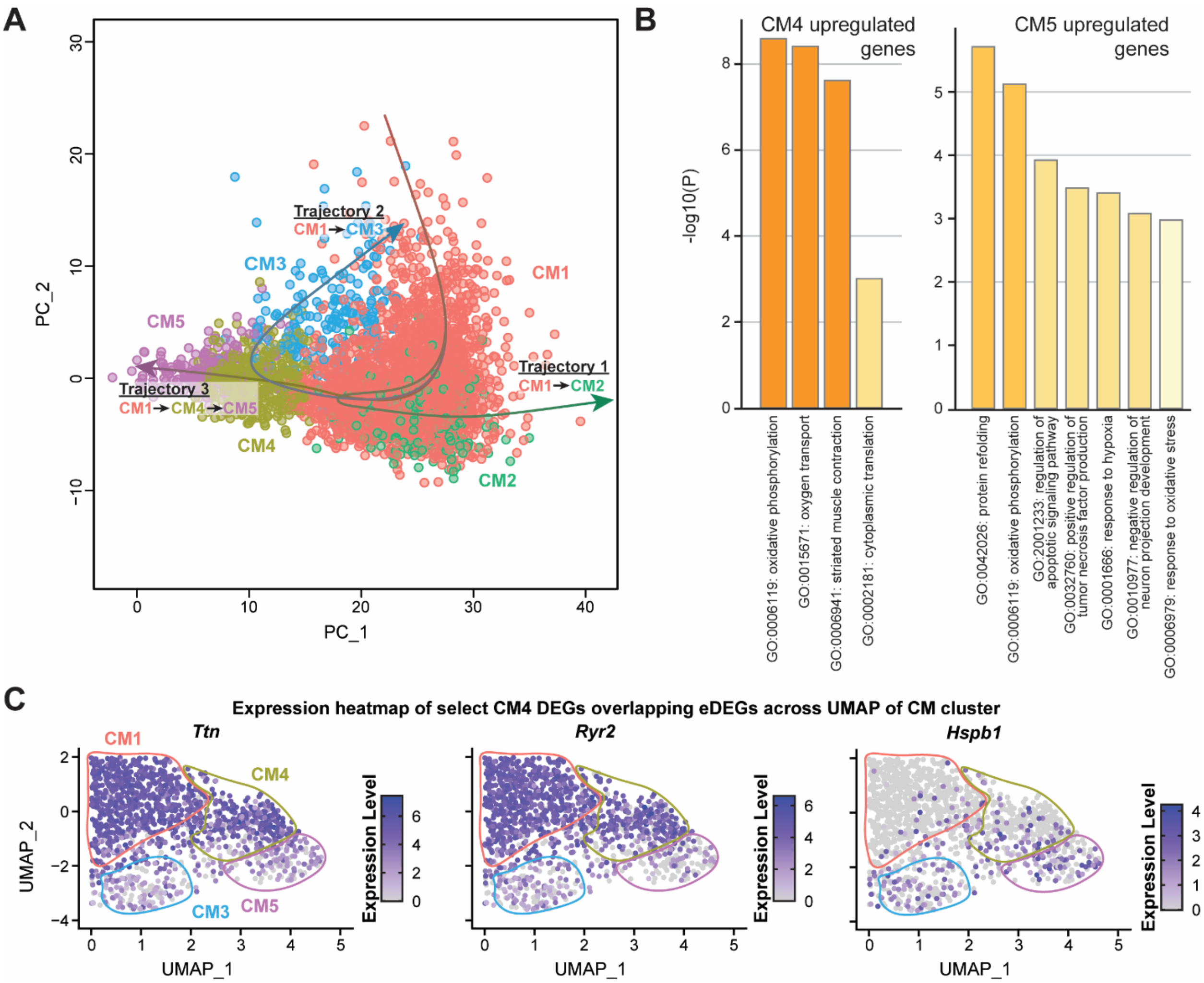


**Supplementary Figure S25: LINC complex disruption restores normal expression of cytosolic DNA sensing genes. (A)** PCA mapping of cardiomyocytes from the snRNA-seq dataset, colors indicate unique subpopulations. Three pseudotime trajectories (CM1 → CM2, CM1 → CM3, and CM1 → CM4 → CM5) are depicted by arrows, indicating that the CM1 population represents an earlier cardiomyocyte state that can progress along three distinct trajectories. **(B)** Barplot showing the significance of gene ontology (GO) term enrichment for genes upregulated in cardiomyocyte subpopulation CM4 (left) and CM5 (right). **(C)** UMAP plot of integrated cardiomyocyte subpopulations (outlined by color) for CM4-specific DEGs overlapping eDEGs (*Ttn* – left, *Ryr2* – middle, *Hspb1* – right). Heatmaps indicate normalized read counts for each gene.

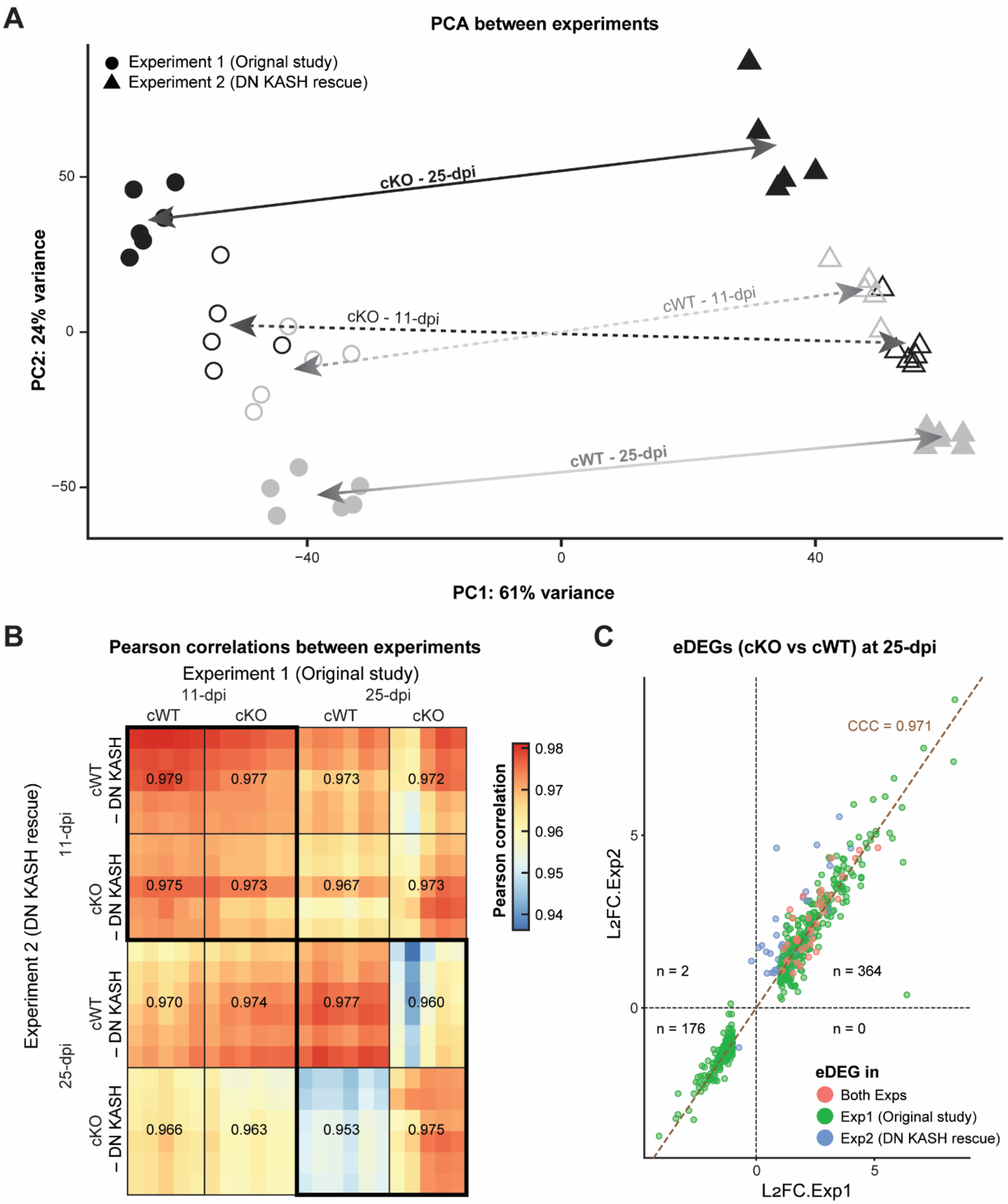


**Supplementary Figure S26: Early disease stages are characterized by greater biological variability.** (**A**) Principal component analysis (PCA) plot of bulk RNA-seq data for cWT (gray) and cKO (black) samples at 11- (open shape) and 25-dpi (filled shape). Samples from both the first (circle, GSE269451 used in Figure 2) and second (triangle, GSE304346 used in Figure 6) experiments are included. Arrows point to the center of centroids for each condition, joining the same experimental condition across experiments. (**B**) Heatmap of Pearson correlations of bulk RNA-seq datasets between the first (x-axis) and second (y-axis) experiments. Rows and columns represent individual samples. Average Pearson correlation values are indicated by condition. Thick borders indicate similar timepoints**. (C)** Scatter plot of $log_2$-fold change (L2FC) for cKO vs. cWT eDEGs in the first (x-axis, used in Figure 2) and second (y-axis, used in Figure 6) experiment. The experiment in which eDEGs were called are indicated by color. The concordance correlation coefficient (CCC) and the $y = x$ identify line are shown in brown.
